# A Pan-Cancer Multi-Omic SuperLearner for Regulated Cell Death Survival Topologies

**DOI:** 10.64898/2026.05.29.728842

**Authors:** Emanuell Rodrigues de Souza, Higor Almeida Cordeiro Nogueira, Victor dos Santos Lopes, Enrique Medina-Acosta

**Author notes:** **Corresponding author**: Enrique Medina-Acosta **Address**: Laboratório de Biotecnologia, Centro de Biociências e Biotecnologia, Universidade Estadual do Norte Fluminense, Avenida Alberto Lamego 2000, Parque Califórnia, Campos dos Goytacazes, RJ, CEP 28015-602, Brazil. **Author Email** Emanuell Rodrigues de Souza, Higor Almeida Cordeiro Nogueira, Victor dos Santos Lopes.

## Abstract

**Introduction:** Regulated cell death (RCD) pathways profoundly influence tumor progression and immune modulation. In prior work, we constructed a comprehensive database mapping 25 forms of RCD across seven multi-omic layers encompassing 33 tumor types (CancerRCDShiny). Despite their robust ability to identify risk populations, translating these prognostic signatures into personalized clinical workflows requires a shift from generalized cohort stratification to individualized risk mapping. This necessitates mapping the complex geometric landscape of patient risk—Survival Topologies—to accurately capture the non-linear dynamics of RCD signatures.

**Methods:** We engineered a Pan-Cancer Multi-Omic SuperLearner pipeline evaluating 33 cancer types. Phase I performed zero-leakage data harmonization and groupwise imputation to prevent cross-cohort amalgamation. Phase II utilized Elastic Net–regularized Cox (CoxNet) regression as an audit-compliant CANARY diagnostic to map mathematical proportional-hazards failures. Admissible strata enforcing a rigid 35% topological missingness barrier entered Phase III, deploying an advanced non-linear Quadripartite Base-Learner Ensemble (Random Survival Forests (*`RSF`*), Extreme Gradient Boosting (*`XGBoost`*), insulated Survival-Boruta, and Multi-Task Logistic Regression (*`MTLR`*))—fused within an Elastic Net Multi-View Meta-Learner (*`MVL`*)—with local interpretability guaranteed via post-hoc *`SHAP`*ley Additive exPlanations (Tree*`SHAP`*) and Local Interpretable Model-agnostic Explanations (*`LIME`*).

**Results:** The CANARY diagnostic empirically proved the structural invalidity of pan-cancer geometric proportional-hazards. Advancing 96 verified matrices into the Quadripartite Machine Learning Ensemble, Phase III executed a structural algorithmic displacement: dense continuous multi-omic topologies computationally suppressed static genomic mutations and Copy Number Variations (CNVs) during multidimensional competition (85.7% vs 0.0% apex retention). Furthermore, the *`MVL`* stabilized global predictions against extreme biological variance, while surrogate *`LIME`* validations (R² < 0.10) confirmed the absolute failure of linear interpretative proxies. Extracting N-dimensional Tree*`SHAP`* interactions natively bypassed generalized risk parameters, mapping exact Survival Topologies. This dynamically exposed multi-omic synergistic (lethal peaks) and antagonistic (protective valleys) rescue trajectories invisible to additive models. We integrated this architecture into CancerRCDPredictor, a Shiny application operating as a digital tumor board.

**Conclusion:** Deploying a Pan-Cancer Multi-Omic SuperLearner to bypass linear topological failures, this study advances beyond generalized cohort stratifications, establishing a deterministically mapped architecture for predicting RCD-related Survival Topologies. Through the CancerRCDPredictor interface, we directly translate multi-omic insights into individualized precision oncology interception.

## 1. Introduction

Regulated cell death (RCD) plays an essential role in tissue homeostasis and cancer treatment response (Galluzzi et al., 2018). Various forms of RCD, including apoptosis, necroptosis, and ferroptosis, distinctly influence tumor progression, immune microenvironment modulation, and therapeutic outcomes. This diverse set of mechanisms provides a unique opportunity to unravel interactions between tumor cells and their microenvironment, particularly within precision oncology frameworks. However, the inherent complexity and interdependence of these processes have historically posed challenges for comprehensive functional characterization and clinical application (Tang et al., 2019).

In a recent comprehensive analysis (Rodrigues de Souza et al., 2025), we constructed an integrated prognostic model that systematically correlated 25 distinct forms of RCD with extensive multi-omic data—including mRNA, miRNA, transcript isoforms, DNA methylation, copy number variation (CNV), mutations, and protein expression—across 33 cancer types from the TCGA study (Goldman et al., 2020, Wang et al., 2022, Li et al., 2024) (see Supplementary Table S1 for specific cancer type abbreviations and full lineage descriptions). In that foundational work, employing a terminological association-based gene selection method, we identified a baseline library of 44,641 multi-omic signatures significantly correlated with various clinical outcomes such as overall survival, mutational burden, and immune infiltration profiles. This model allowed us to distinguish clinically relevant molecular signatures, effectively associating them with immunologically active ("hot") and immune-suppressed ("cold") deconvoluted immune cell and tumor microenvironment phenotypes. Such classification provides critical insights for selecting appropriate immune-based therapeutic strategies tailored to individual patient profiles.

To facilitate practical clinical applications, we also developed a standardized alphanumeric nomenclature system categorizing multi-omic signatures by biological function, specific RCD mechanisms, and key clinical variables, and survival metrics (Rodrigues de Souza et al., 2025). (see Supplementary Note 1 for a detailed methodological framework of the 11-component signature identifier). This approach enhances interpretability, accelerates biomarker identification, and supports the seamless integration of these signatures into clinical workflows, promoting their practical use in precision medicine.

Our prior studies also highlighted the critical importance of transcript-level specificity (Rodrigues de Souza et al., 2025). For instance, *MAPK10* demonstrated significant clinical correlations predominantly through specific transcript isoforms, emphasizing the complexity and importance of alternative splicing and transcriptional control mechanisms in cancer biology. Conversely, genes such as *COL1A1* and *UMOD* exhibited uniform phenotypic correlations across isoforms, suggesting coordinated regulatory mechanisms at the gene level linked to tumor stemness and aggressiveness. We identified 879 clinically meaningful multi-omic signatures that included known immunotherapy targets under clinical investigation, reinforcing their immediate translational relevance. To further enhance the practical exploration and visualization of these comprehensive datasets, we developed CancerRCDShiny (https://cancerrcdshiny.shinyapps.io/cancerrcdshiny/), an interactive analysis tool accessible to the broader research community.

Despite the robust prognostic insights generated by this foundational multi-omic catalog, translating these findings into routine precision oncology workflows requires shifting from descriptive stratification toward precise, individual-level prediction. Prognostic models inherently identify generalized biological risk groups, but they lack the algorithmic complexity necessary to deliver exact patient survival prediction. Achieving this transition requires overcoming profound multidimensional challenges—including extreme data sparsity, missingness, and structural hazards that routinely cause traditional unregularized models to collapse when faced with high-dimensional RCD omic signatures.

To bridge the gap between descriptive prognosis and computational prediction, the present study establishes a Pan-Cancer Audit-Compliant SuperLearner—a formal mathematical architecture for Machine Learning Integration (commonly known as Stacked Generalization (Wolpert 1992, van der Laan et al., 2007))—engineered specifically for rigorous predictive benchmarking of Patient Survival from RCD omic signatures.

Rather than applying predictive algorithms blindly, we deploy a strictly deterministically gated, three-phased architecture. Phase I and Phase II act as formal execution barriers to prevent cross-layer biological data leakage and to mathematically diagnose the geometric survival topologies of 33 distinct tumor types. Only mathematically verified strata advance to Phase III, where an advanced non-linear supervised Quadripartite ML ensemble—combining Random Survival Forests (*`RSF`*) (Ishwaran et al., 2008), Extreme Gradient Boosting (*`XGBoost`*) (Chen et al., 2016), Survival-*`Boruta`* (Kursa et al., 2010), and Multi-Task Logistic Regression (*`MTLR`*) (Yu et al., 2011)—operates concurrently with strict post-hoc interpretability algorithms (*`SHAP`*ley Additive exPlanations (*`SHAP`*) (Lundberg et al., 2017) and Local Interpretable Model-agnostic Explanations (*`LIME`*) (Ribeiro et al., 2016). Ultimately, by resolving the high-dimensional sparsity of the prior multi-omic catalog, this framework successfully shifts the biological logic of the 25 distinct forms of RCD into an actionable, fully transparent predictive architecture.

This transition from identifying prognostic factors to developing actionable predictive models represents a significant methodological advancement essential for personalized medicine. This approach holds great promise for improving patient stratification, clinical decision-making, and therapeutic efficacy, substantially enhancing patient management in oncology.

## 2. Materials and Methods

### 2.1. Overview of the analytical pipeline

To execute robust patient survival prediction via RCD omic signatures, the analysis is organized as a pan-cancer, audit-compliant SuperLearner pipeline. Designed to separate data transformation, eligibility gating, and inferential modeling into distinct operational stages, this framework prevents cross-layer biological leakage and prohibits outcome-driven adaptation.

Phase I generates a library of candidate preprocessing regimes using standard statistical transformations within isolated cancer cohorts. Phase II applies deterministic eligibility and feasibility gates—implemented as Transformation Admissibility Routing (TAR) and a CANARY model—to certify which of these clinical-omic regimes are structurally sound. Only mathematically verified cohorts pass to Phase III, which serves as the exclusive locus for predictive inference. Within Phase III, an advanced non-linear Quadripartite ML Ensemble is derived solely on TAR-admissible regimes, ensuring that any resulting patient survival prediction reflects intrinsic biological structure rather than hidden optimization. A schematic overview of this audit-compliant pipeline, defining phase boundaries and execution gates, is provided in Figure 1.

**Figure 1.**
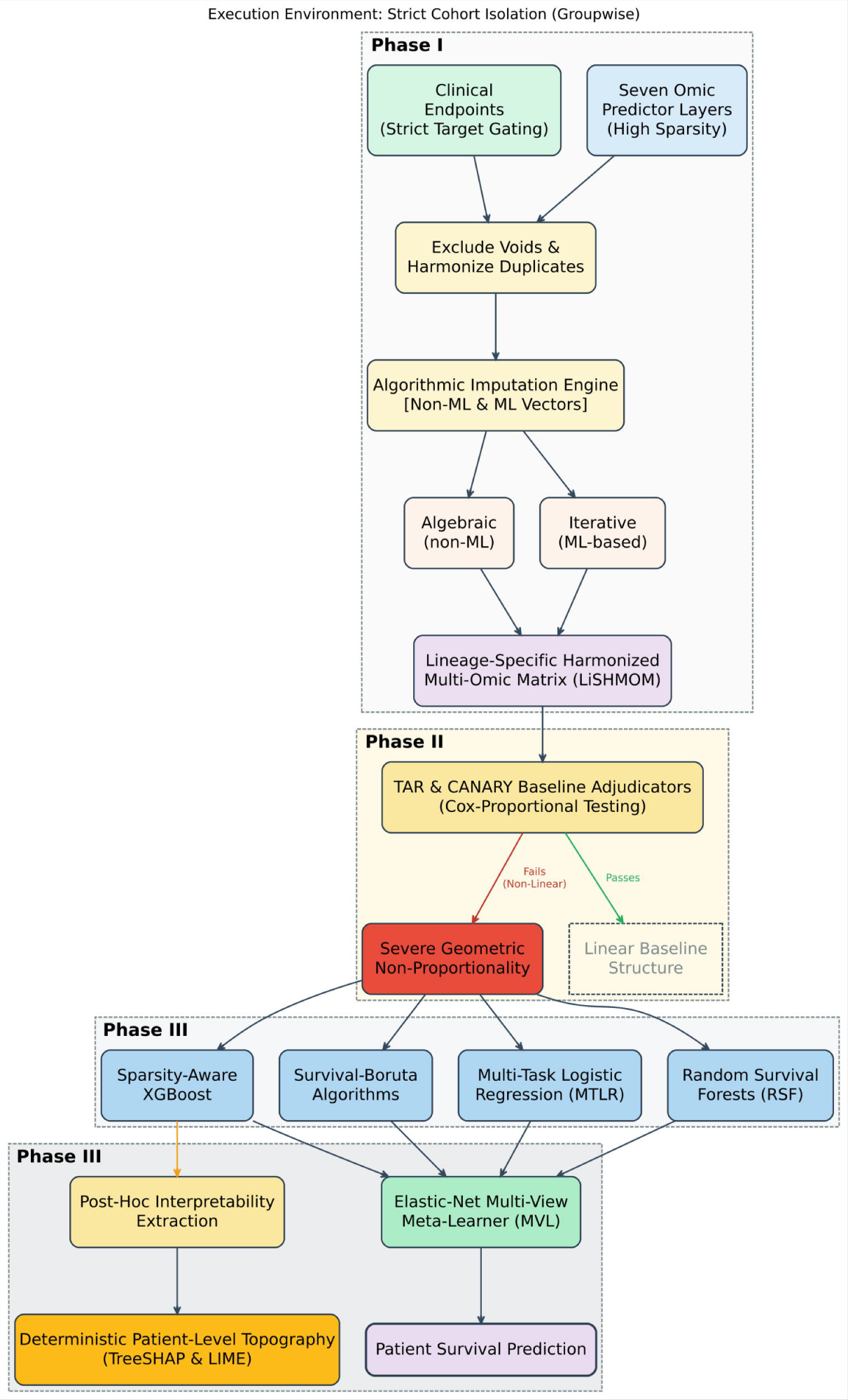
Analytical Pipeline Architecture: Strict Cohort Isolation and SuperLearner Execution Strategies. Phase I: Raw multi-omic inputs (seven highly sparse omic predictor layers) and strictly gated clinical endpoints undergo a domain-specific harmonization process. All computations operate under a mathematically enforced zero-leakage policy; variables are processed within strictly isolated, cancer-specific groupwise layers. Missing feature spaces are addressed using a dual Algorithmic Imputation Engine—deploying both non-ML algebraic techniques and ML-based iterative solutions—to yield standardized, robust Lineage-Specific Harmonized Multi-Omic Matrices (LiSHMOM) for each pan-cancer lineage. Phase II: The CANARY Baseline Adjudicator rigorously tests the resulting matrices for Cox-Proportional topological baseline failures. Phase III: Features exhibiting severe geometric non-proportionality are bypassed into a sophisticated Quadripartite ML Ensemble. This array leverages four discrete, non-linear algorithms: *`RSF`*, Sparsity-Aware *`XGBoost`*, Survival-*`Boruta`* parameters, and *`MTLR`*. Finally, the predictive vectors from these individual base-learners are systematically synthesized via the *`MVL`*. This definitive SuperLearner integration mitigates individual algorithmic bias, establishing patient survival predictions while utilizing Post-Hoc Interpretability Extraction (*`TreeSHAP`* and *`LIME`*) to yield highly transparent, deterministic patient-level topographies for downstream clinical deployment.

All analyses in Phases I–III were executed under a fixed set of constitutional contracts defining admissible operations on cohorts, predictors, and execution units. These contracts enforce strict groupwise analysis by cancer type and survival endpoint, prohibit predictor-driven sample reduction, and require explicit, auditable justification for any predictor exclusion. A full specification of these contracts and their audit requirements is provided in Supplementary Note 2.

### 2.2. Phase I: Programmatic Retrieval, Harmonization, and Algorithmic Reconstruction of Multi-Omic and Clinical Data

The primary function of Phase I is the programmatic harmonization and deterministic reconstruction of high-dimensional prognostic matrices. Crucially, Phase I operates strictly as an outcome-independent data processing boundary enforced by rigorous, multi-layered missingness gates designed to prevent the indiscriminate fabrication of synthetic data.

For clinical survival metrics, we enforced an absolute prohibition on the imputation of survival events (OS, DSS, DFI, PFI) to prevent label leakage and optimism bias. Conversely, the imputation of survival times was permitted but governed by three stringent clinical brakes: (1) a variable-level missingness threshold (≤35%), (2) a type-pair gate requiring adequate intra-cohort event prevalence (0.05–0.95), and (3) a strict row-wise constraint mandating that the event status and foundational clinical anchors (e.g., diagnosis year) be observed.

For multi-omic predictors (both binary/categorical and continuous), imputation was strictly bounded by the same ≤35% intra-cohort missingness barrier. To maintain structural dataframe geometry without massive synthetic data fabrication, omic variables exceeding this missingness threshold were deliberately insulated; their missing values were carried forward untouched to downstream machine learning evaluators. Furthermore, for continuous subsets where extreme sparsity triggered dimensionality failures in multivariate algorithms (which mathematically require ≥2 features), we engineered a deterministic, non-informative dummy-variable injection protocol to satisfy algorithmic constraints without violating biological integrity or resorting to cross-cohort data pooling.

By passing through these gates, eligible molecular values were then inferred in a layer-specific, self-reconstruction mode using a comprehensive suite of algorithmic strategies. This suite ranged from deterministic methods (mean, median, mode) and Bernoulli-based probabilistic sampling for binary features (preserving sample-level heterogeneity), to advanced multivariate approaches including k-nearest neighbors (*`kNN`*) (Cover et al., 1967), Multiple Imputation by Chained Equations (*`MICE`*) (Buuren et al., 2011), Iterative Rank-Restricted Soft Singular Value Decomposition (*`softImpute iSVD`*) (Hastie et al., 2015), and non-parametric architectures like *`missForest`* (Stekhoven et al., 2012), light gradient boosting machine (*`LightGBM`*) (Ke et al., 2017), and *`XGBoost`* (Chen et al., 2016). Throughout this entire architecture, survival endpoints remained fully insulated from the predictor structure, serving exclusively as downstream audit sentinels.

To operationalize this rigorous architectural framework, molecular and clinical data were programmatically retrieved from TCGA using the UCSCXenaShiny resource (Wang et al., 2022, Li et al., 2024). To structure this high-dimensional encoding, we established the unified baseline decoder (Supplementary Dataset S1). Derived from an initial baseline universe of 44,641 pan-cancer multi-omic signatures from our previous work (Rodrigues de Souza et al., 2025), this master decoder was rigorously filtered to isolate features demonstrating robust clinical relevance. Specifically, signatures retained in Supplementary Dataset S1 were required to achieve statistical significance in at least one univariate survival metric (log-rank and Cox proportional hazards indicating definitive risk or protection) and demonstrate significant associations with deconvoluted immune infiltrates (classified as "Hot", "Cold", "Variable", or "NS") and tumor microenvironment profiles ("anti-tumoral", "pro-tumoral", "dual", or "NS"). Through this rigorous selection, we curated a highly stabilized catalog of 14,907 distinct multi-omic prognostic nomenclatures, which collectively encompass 17,875 unique biological target elements (spanning gene symbols, transcript isoforms, miRNAs, and proteins). By formally formatting these targets under the variables Signature, Nomenclature, CTAB (cancer type abbreviation), and Omic_feature, and mapping each abstract alphanumeric token directly to its constituent genetic elements and functional Regulated Cell Death (RCD) mechanics, Supplementary Dataset S1 served as the invisible architectural blueprint dictating the extraction and formatting of the multi-omic features subsequently compiled into the clinical modeling matrix.

Clinical, demographic, and survival metadata were programmatically retrieved from the TCGA using the UCSCXenaShiny package (Wang et al., 2022, Li et al., 2024). To construct the foundational dataset, distinct multi-omic expression profiles—encompassing mRNA, miRNA, protein, transcript isoforms, CNVs, DNA methylation, and somatic mutations—were systematically aggregated into a unified, high-dimensional matrix. To ensure structural integrity during assembly, malformed entries were excluded, and the molecular profiles were merged with the clinical-survival records via unique sample identifiers. Finally, the resulting matrix underwent strict deduplication and was filtered to eject any instances exhibiting total systemic missingness across all molecular and response features. This finalized, cleaned dataset was subsequently serialized as the foundational baseline to feed the downstream harmonization and imputation architectures.

During this deduplication process, repeated patient entries were systematically isolated and examined. For each duplicated group, clinical-survival variables were cross-referenced. Where discrepancies existed, a strict administrative harmonization protocol was applied, replacing fragmented records with values from the most complete row to ensure absolute internal consistency across all binary outcome indicators and survival time variables (Overall Survival [OS], Disease-Specific Survival [DSS], Disease-Free Interval [DFI], and Progression-Free Interval [PFI]). Crucially, this procedure functioned purely as deterministic harmonization for schema auditing; it did not involve any statistical modeling, probabilistic sampling, or stochastic alteration of survival outcomes. These administratively consolidated blocks were subsequently used to overwrite the original duplications, yielding the definitive dataset.

Prior to advanced imputation, foundational predictors were structurally preprocessed. Copy Number Variation (CNV) features (marked by the “.3” nomenclature suffix) were transformed into explicit categorical states (“Deleted”, “Normal”, or “Duplicated”), while somatic mutation features (denoted by “.2”) were binarized to indicate wildtype (0) or mutated (1) status. This natively observed matrix was designated as the df005 pre-imputation reference baseline (Supplementary Dataset S2). Serving as the fixed architectural anchor against which all downstream imputation variants were evaluated, the harmonized df005 baseline comprises 10,489 pan-cancer samples and 14,657 distinct variables (consisting of 62 clinical/demographic anchors and 14,595 multi-omic features).

### 2.3. Omic Token Encoding and Predictor Layer Abstraction

The biological definition of multi-gene signatures and their molecular composition has been described in detail in our prior work (Rodrigues de Souza et al., 2025, Almeida Cordeiro Nogueira et al., 2026a, Almeida Cordeiro Nogueira et al., 2026b). In the present study, we focus on the computational encoding used to operationalize these predictors for large-scale preprocessing and modeling. Each molecular predictor was encoded using a tokenized nomenclature system in which a numeric suffix embedded in the Nomenclature field identifies its omic layer. Seven layers were represented: protein abundance (.1), somatic mutation status (.2), CNV (.3), microRNA expression (.4), transcript isoform abundance (.5), mRNA expression (.6), and CpG methylation (.7). This token-based abstraction enabled programmatic parsing of feature type, enforcement of layer-specific preprocessing and imputation rules, and stratified handling of predictors across cancer types. By decoupling biological identity from measurement modality, the pipeline ensured that transformations and missing-data strategies were applied consistently within each omic class while preserving cross-layer comparability. The token system further supported deterministic audit logging and downstream admissibility evaluation in Phase II.

### 2.4. Clarification on Survival Variable Handling in Phase I

To prevent label leakage and optimism bias, survival variables were strictly insulated from statistical imputation. Instead, a severely restricted administrative completion protocol was applied exclusively to resolve deterministic schema inconsistencies between event indicators and their paired time-to-event axes. Confined to a negligible fraction of records, these logical alignments did not introduce synthetic biological signals or alter the underlying cohort censoring structure. Consequently, all Phase I datasets containing administratively aligned survival fields were treated strictly as tentative preprocessing variants. Their final admissibility for downstream machine learning was adjudicated by the topological baseline adjudicator in Phase II, where survival endpoints served exclusively as quality-control sentinels. This structural audit guaranteed that the tightly gated multi-omic imputation—which was mathematically required to resolve NA values and feed the non-linear Phase III architectures—did not distort the baseline outcome topography.

### 2.5. Distinction Between Survival Harmonization and Time Completion in Phase I

Survival-related processing in Phase I comprised two conceptually distinct operations. First, deterministic harmonization was performed to reconcile endpoint labels and observed time-to-event values across authoritative clinical inputs and duplicate records, yielding a coherent ground-truth survival schema that was frozen prior to downstream processing. Second, limited administrative completion was applied exclusively to time-to-event variables that were missing at baseline, under strict eligibility and consistency constraints, with policy enforcement restricted to imputed cells only. Survival event indicators were not modified after harmonization, and observed (non-missing) baseline times were never altered. These two operations were intentionally separated to preserve censoring structure and ensure that downstream analyses evaluated preprocessing sensitivity auditing rather than inferential survival estimates.

### 2.6. Missing data imputation

To address the challenge of missing data in high-dimensional pan-cancer datasets, we implemented a modular, fault-tolerant imputation pipeline that sequentially integrates CNVs, mutation status, and continuous molecular features across seven omic layers. Survival endpoints were retained throughout the pipeline exclusively as audit and consistency variables and were not subjected to statistical imputation for inferential purposes. Importantly, survival endpoints were never used as predictors, targets, loss functions, or optimization objectives for missing-value reconstruction of omic features.

Limited administrative completion of survival outcome indicators (OS, DSS, DFI, PFI) and their corresponding time-to-event variables was performed under strict, deterministic logical consistency rules to resolve internal schema inconsistencies (e.g., alignment between event indicators and time variables) and to support downstream preprocessing sensitivity auditing. These operations were deterministic, auditable, and affected a mathematically negligible fraction of the global dataset exclusively restricted to the alignment of time-to-event variables (Overall Survival Time [OS.time] = 0.52%; Disease-Specific Survival Time [DSS.time] = 0.94%; Progression-Free Interval Time [PFI.time] = 0.42%). The administratively aligned fraction for Disease-Free Interval Time (DFI.time = 9.96%) was proportionately elevated strictly due to the expected structural missingness of ‘disease-free’ clinical states in specific cohorts (e.g., glioblastoma, where total resection is clinically implausible). Crucially, the fundamental clinical event indicators (OS, DSS, PFI, DFI) were explicitly locked and mathematically forbidden from alteration, ensuring no synthetic biological signal was introduced into the survivorship topography. Administrative completion was applied per cancer type, yielding three preprocessing variants with administrative completion of survival time fields (df006–df008), used solely for preprocessing sensitivity auditing.

Each of the three administratively completed survival variants was then subjected to CNV imputation using three methods—mode (most frequent value), random sampling, and k-nearest neighbors (*`kNN`* via *`VIM::kNN`*)—executed groupwise by tumor type. CNV values were pre-standardized to categorical states ("Normal", "Duplicated", "Deleted"). When *`kNN`* failed because of low sample size or convergence limitations, mode imputation was used as a fallback. This produced nine CNV-imputed datasets (df009–df017).

Mutation features, previously binarized (0 = wild-type, 1 = altered), were imputed in these nine datasets using four strategies: mean, median, mode, and Bernoulli sampling based on empirical alteration frequencies. This resulted in 36 mutation-imputed datasets (df018–df053).

Each mutation-imputed dataset served as input for imputation of continuous omic features—including protein expression (.1), miRNA (.4), transcript isoform abundance (.5), mRNA expression (.6), and CpG methylation (.7). Nine distinct imputation algorithms were applied independently to each dataset: mean, median, random sampling, *`kNN`*, *`MICE`*, *`missForest`*, *`XGBoost`*, *`LightGBM`* and *`iSVD`*. This final step produced 372 Lineage-Specific Harmonized Multi-Omic Matrices (LiSHMOM), 324 of which are continuous-layer–imputed datasets organized by method: mean (df054–df089), median (df090–df125), random (df126–df161), *`kNN`* (df162–df197), *`missForest`* (df198–df233), *`XGBoost`* (df234–df269), *`LightGBM`* (df270–df305), *`MICE`* (df306–df341) and *`iSVD`* (df342-df377) (Supplementary Figure S1; Supplementary Table S2).

At each stage of the pipeline, intermediate matrices were systematically version-controlled and dynamically purged from active memory to optimize computational elasticity. A dynamic checkpoint engine automatically resumes the pipeline from the last successfully validated output in the event of hardware failure. Every generated dataset was indexed and registered in a centralized architectural ledger, enabling absolute deterministic traceability for downstream modeling. This framework yielded 372 version-controlled preprocessing variants (df006–df377), materialized as indexed data objects, which constitute candidate preprocessing regimes generated in Phase I. These variants were not assumed to be admissible for modeling a priori; their admissibility was evaluated exclusively by TAR in Phase II on a cancer- and endpoint-specific basis.

### 2.7. Statistical Rationale for Layer-Specific Imputation Architectures

The multi-layered imputation strategy described above was explicitly tailored to respect the distinct biological and statistical topologies of each omic data type. For categorical CNV states ("Normal", "Duplicated", "Deleted"), imputation was restricted to domain-appropriate strategies to ensure cytogenetic constraints were not violated. Crucially, a programmatic fallback logic automatically defaulted to mode imputation whenever *`kNN`* convergence failed due to cohort-specific multicollinearity or low sample sizes.

For binarized somatic mutations, Bernoulli-based probabilistic sampling was specifically integrated alongside standard estimators. The Bernoulli method was critical for mutation layers because it maintains the empirical alteration frequency across tumors, thereby preserving the stochastic representation of both rare and ubiquitous driver events without introducing fractional artifacts.

Finally, the deployment of nine distinct algorithms across the continuous features (protein abundance, miRNA, mRNA, transcript isoforms, and CpG methylation) was designed to comprehensively sweep the spectrum of modern imputation philosophies: parametric estimators (mean, median), local topological similarity (*`kNN`*), iterative ensemble regression (*`missForest`*), gradient boosting engines (*`XGBoost`*, *`LightGBM`*), and probabilistic chaining (*`MICE`*, *`iSVD`*). By deploying this architectural diversity across five highly heterogeneous continuous layers, the pipeline guaranteed a robust sensitivity audit. This ensured that the predictive Phase III machine-learning models could be evaluated against preprocessing strategies capable of handling extreme data sparsity, non-linear dependencies, and massive dimensionality.

### 2.8. Evaluation Strategy for Imputation Performance Across Output Datasets

To ensure the mathematical validity and interpretability of each imputation method, a systematic evaluation framework was implemented to audit the downstream effect of predictor imputation on survival-related signal preservation. At every stage of the pipeline, intermediate matrices were systematically version-controlled and registered in a centralized tracking ledger. This architectural design ensured deterministic mapping from raw inputs to imputed variants, guaranteeing absolute fault tolerance and reproducibility throughout the pipeline.

The complete set of 372 preprocessing variants was analyzed using a structured diagnostic module that computed concordance indices (C-index) for each survival endpoint (OS, DSS, DFI, PFI) via Cox proportional hazards models. Crucially, these models were deployed strictly as diagnostic probes of signal preservation rather than as inferential endpoints. For each imputation method, C-index trajectories were compared across endpoints and plotted against the baseline percentage of missingness in the original data. This permitted a rigorous evaluation of algorithmic robustness (stability under high missingness) versus informativeness (preservation of prognostic topological structure).

Furthermore, pairwise correlations between data missingness and diagnostic performance were computed per imputation strategy and endpoint. This structural audit enabled the explicit identification of methods susceptible to artifactually inflated performance (e.g., complete-case baseline distortions) versus algorithms that maintained independence from missing data proportions. Ultimately, these diagnostics did not dictate direct model selection; rather, they functioned as absolute admissibility gates to qualify matrices for downstream machine-learning evaluation.

### 2.9. Phase I Imputation Constraint and Hallucination Audit

To explicitly prevent synthetic data hallucination and preserve cohort geometry, the Phase I architecture enforced strict algorithmic gates and downstream topological brakes. Multi-omic parametric imputation was restricted exclusively to predictors exhibiting ≤35% groupwise missingness. Across the pan-cancer matrix, this eligible pool comprised 592,191 global missing cells. However, due to strict secondary algorithmic constraints—such as matrix convergence limitations and multicollinearity brakes integrated within the *`MICE`* and *`iSVD`* engines—the pipeline effectively imputed only ∼11.8% of these eligible cells. The remaining 523,587 cells were safely carried through the pipeline as untouched residual NA values in the final matrices (e.g., df377). Furthermore, to guarantee absolute architectural separation between predictor imputation and clinical outcomes, the pipeline enforced a hard structural lock on all survival endpoints. Tracking the 852 baseline NA cells present across the eligible survival time variables in the raw matrix revealed that exactly 852 NA cells remained in the fully imputed variants. This mathematically proves that none of the nine continuous machine-learning imputation engines ever breached the clinical survival boundary, definitively ensuring that no inferential survival signal was synthetically generated.

### 2.10. Universal Resume Engine and Execution Guarantees

Imputation procedures in Phase I were executed using a fault-tolerant, audit-oriented execution engine implemented in R for large-scale, groupwise multi-omic processing. The engine supports multiple imputation strategies spanning parametric, similarity-based, ensemble, boosting, and matrix-completion approaches. Imputation was performed independently within cancer-type strata to prevent cross-cohort contamination.

The execution framework enforces deterministic checkpointing, full provenance tracking, and version-controlled materialization of each preprocessing regime as an indexed RDS object registered in a centralized audit table. Intermediate failures are isolated at the variable or method level and do not interrupt global pipeline completion. This design ensures that all preprocessing variants entering Phase II are reproducible, auditable, and free of cross-endpoint or cross-cancer information leakage.

Throughout the manuscript, the term *candidate preprocessing regime* refers to any Phase I variant in the df006–df377 series, whereas *TAR-admissible preprocessing regimes* refer specifically to the subset of these variants classified as Unchanged or Improved by TAR for a given cancer × endpoint stratum. Canonical terminology definitions used throughout Phase I–III analyses are summarized in Supplementary Table S3.

Implementation details, operational audit criteria, and software validation procedures for the Phase I imputation engine are provided in the Supplementary Information.

### 2.11. Phase II: TAR-compliant survival modeling feasibility and execution framework

Phase II introduces supervised survival modeling exclusively as a diagnostic mechanism rather than a predictive exercise. Elastic Net–regularized Cox (*`CoxNet`*) regression is deployed as a CANARY model to probe the structural admissibility of proportional-hazards geometry within each cancer–endpoint–preprocessing stratum, under fixed and auditable constraints. Models are executed under fixed, pre-specified identifiability constraints and deterministic regularization ladders, and are evaluated solely for feasibility properties such as convergence, coefficient stability, and absence of degenerate risk scores.

No predictive optimization, hyperparameter tuning, feature selection, or model comparison is performed in Phase II. Survival outcomes are used only to test whether a given preprocessing regime supports a coherent hazard representation under auditable constraints. Model failure is interpreted as a diagnostic signal of data geometry (e.g., nonlinearity, time dependence, information insufficiency), not as a modeling deficiency. Thus, Phase II uses supervised modeling as a structural audit mechanism, not as a predictive exercise.

Phase II was designed as a strictly executional stage that operationalizes survival modeling under the constraints imposed by TAR, which functions as a pre-modeling validity gate that evaluates whether upstream preprocessing regimes preserve survival-relevant structure for each cancer type and survival endpoint. Each preprocessing regime represents a distinct, auditable combination of clinical harmonization and omic missingness handling generated in Phase I (materialized as indexed datasets df006–df377). Using survival endpoints exclusively as external quality-control probes within the preprocessing admissibility stage, TAR classifies each regime, for each cancer–endpoint pair, as Unchanged, Improved, or Degrade relative to a fixed unimputed reference (df005). Only regimes classified as Unchanged or Improved are admissible for Phase II. TAR is explicitly diagnostic and exclusionary; it does not constitute model selection, performance optimization, feature selection, or cohort harmonization, and TAR decisions are never revised during Phase II. Phase II did not generate or modify preprocessing datasets. Instead, TAR evaluated Phase I candidate preprocessing regimes (df006–df377) and annotated their admissibility on a cancer- and endpoint-specific basis.

Phase II respects TAR admissibility as a hard gate and executes supervised survival model forms executed as feasibility probes only on TAR-approved preprocessing regimes. All Phase II operations are endpoint-specific: OS, DSS, DFI, and PFI define independent cohorts. Missingness in one endpoint never excludes a sample from analysis of another endpoint. No global completeness constraints are imposed at any stage; in particular, Phase II prohibits complete-case filtering across predictors or omic layers. The only permissible sample-level exclusion is missing or schema-invalid survival information for the specific endpoint under analysis. Phase II does not modify, tune, or re-evaluate upstream preprocessing strategies and does not select preprocessing regimes based on Phase II outcomes.

All Phase II decisions operate within a single execution unit defined by the tuple (cancer type *c*, survival endpoint *m*, dataset variant *d*, algorithm *a*). No rule, filter, or feasibility decision is permitted to operate outside this unit. This design prevents implicit information sharing across cancers, endpoints, preprocessing regimes, or algorithms.

Survival cohort definition in Phase II is strictly endpoint-scoped. For a given cancer type and endpoint, survival time and event indicators are extracted and validated against a canonical schema: survival time must be finite and non-negative when observed, with time equal to zero explicitly permitted; negative survival times are invalid and trigger hard failure. Event indicators must be binary (0/1) when observed. Samples are retained if and only if both time and event are observed and schema-valid for the selected endpoint. No predictor-driven row deletion is permitted. Cohort masking is therefore defined exclusively by survival feasibility for the endpoint under analysis, ensuring that cohorts are neither intersected across endpoints nor altered by predictor missingness.

Predictor handling in Phase II is algorithm-aware. Algorithms are partitioned according to their tolerance for missing predictor values. Tree-based survival models are permitted to operate directly on matrices containing missing values, whereas linear or finite-matrix models require fully finite numeric predictors.

Residual predictor missingness after TAR-approved preprocessing is resolved exclusively at the machine-learning execution stage through a hierarchical safety mechanism. Feature-level removal is applied first to eliminate numerically inadmissible predictors within a local cancer × endpoint × preprocessing regime stratum. Only if missingness persists and the selected algorithm requires a finite design matrix is deterministic, outcome-independent imputation applied as a last resort. Sample-level exclusion due to predictor missingness is explicitly disallowed.

For algorithms requiring finite matrices, residual predictor missingness after TAR-approved preprocessing is handled through a strictly ordered, local repair hierarchy applied only within the execution unit. First, the dataset is used as provided. Second, predictors with excessive missingness are removed at the feature level only; sample-level deletion is prohibited. Third, remaining missing values are deterministically imputed using outcome-independent, model-independent rules computed locally within the endpoint-scoped cohort. All such operations are auditable and logged. Global predictor intersections, stochastic imputation, outcome-aware imputation, and cross-endpoint cohort enforcement are explicitly prohibited.

A central component of Phase II is the use of *`CoxNet`* regression as a structural feasibility probe rather than as a universal predictive model. *`CoxNet`* operationalizes the CANARY diagnostic under fixed statistical identifiability requirements—including minimum sample size and minimum number of observed events (detailed in Supplementary Note 2)—and under a deterministic μ-ladder that progressively applies increasingly permissive structural relaxations. These relaxations natively include variance filtering, elastic-net path constraint adjustments, univariate prescreening with collinearity pruning, and ridge stabilization. At each μ level, model fitting is evaluated solely for structural feasibility: convergence, stability of coefficients, and non-degenerate risk scores. No performance optimization, hyperparameter tuning, or cross-validation-based selection is performed.

Successful feasibility indicates that the stratum geometrically admits a sparse, approximately linear hazard representation consistent with proportional-hazards assumptions under auditable constraints, without implying superiority over alternative modeling families. Conversely, systematic μ-ladder exhaustion or early data infeasibility is interpreted as concrete evidence of dense, nonlinear, time-varying, or information-deficient survival structure incompatible with sparse PH modeling. Importantly, these outcomes are treated as intrinsic diagnostic properties of the data rather than modeling failures. The CANARY role of *`CoxNet`* therefore provides a principled, model-based structural audit prior to any downstream inference.

Throughout Phase II execution, extensive structural audits are enforced to ensure reproducibility and traceability. These include strict schema validation, row-identity preservation utilizing explicit sample identifiers, alignment checks between predictor matrices and survival vectors, and external-universe audits that verify cancer and endpoint coverage against a reference manifest rather than relying on TAR-filtered inputs. Parallel execution is performed using deterministic, audited multisession workflows, with per-stratum worker provenance permanently recorded to certify actual parallel execution.

The absolute output of Phase II is a stratum-resolved comprehensive feasibility log indexed by cancer type, survival endpoint, and preprocessing regime. This log explicitly records survival cohort sizes, event counts, predictor dimensionality before and after feasibility repair, feature-level retention ratios, sample-level retention ratios, μ-ladder diagnostics, explicit failure codes, and audit metadata. It functions exclusively as a formal eligibility and classification artifact that captures survival data sufficiency and geometric compatibility; it does not encode model performance, parameter estimates, or inferential results. In the subsequent Results section, these feasibility outcomes are summarized descriptively to document the distribution of identified survival geometry classes.

No parameter estimates, risk scores, model coefficients, or performance metrics generated in Phase II are retained, reported, or reused in any downstream analysis.

### 2.12. Phase III: Supervised ML for outcome prediction

Only after preprocessing regimes have passed formal eligibility and feasibility gates does the analysis proceed to Phase III. This phase constitutes the exclusive stage for predictive modeling, in which supervised survival models are applied under strictly controlled, TAR-compliant conditions.

Phase III model-family authorization is determined exclusively by Phase II structural diagnostics. In principle, strata supporting sparse proportional-hazards geometry would be eligible for Cox-based inference. However, Phase II feasibility analysis revealed that no evaluable cancer–endpoint stratum satisfied sparse proportional-hazards compatibility under the specified auditable constraints. All modelable strata were therefore classified as structurally proportional-hazards–incompatible but modelable, indicating the absence of sparse, approximately linear hazard structure despite sufficient survival information for downstream inference (see Supplementary Note 2 for mathematical justification).

Accordingly, Phase III is designed to deploy survival modeling families capable of accommodating nonlinear, interaction-rich, or time-varying hazard structures, including nonlinear ensemble methods and non–proportional-hazards architectures. These modeling families are incorporated sequentially within the same Phase III framework, with each algorithm specified, implemented, and evaluated independently under identical structural and audit constraints.

All Phase III analyses are restricted to TAR-admissible preprocessing regimes, and cohort composition remains fixed within each cancer–endpoint stratum. No preprocessing, feature filtering, or cohort modification occurs at this stage. Phase III does not revisit TAR classifications or Phase II feasibility outcomes, and no Phase III model performance metric is permitted to retroactively alter upstream eligibility determinations.

To preserve the strict biological fidelity of the ensemble predictions and mathematically isolate the models from healthcare administration bias, a rigorous, runtime geometric exclusion filter was deployed prior to ensemble synthesis. During dynamic execution of the Phase III SuperLearner, the topology of missing data was audited row-wise within the boundaries of each cancer-specific predictive matrix (Supplementary Table S4). Crucially, the 35% topological missingness barrier was established as a strict operational policy to intercept severe bimodal sparsity patterns endemic to high-dimensional pan-cancer arrays. This conservative threshold was explicitly designed to capture and surgically excise ‘clinical ghost’ vectors—patient samples where the absence of clinical profiling outweighs verifiable biological mass. By categorically excluding patient samples presenting with missing data spanning ≥ 35% of their explicit lineage-specific omic predictors, the architecture was computationally protected from algorithms falsely mapping extensive missingness (NA) into artificial default trajectories of protective or deleterious risk.

Patients exhibiting partial missingness (< 35%) were retained to preserve the topological density of the cohort matrix. For these retained samples, missing data was systematically managed according to the mathematical structure of the terminal algorithms: tree-based gradient boosting (*`XGBoost`*) actively utilized sparsity-aware split finding to incorporate missingness patterns natively without data alteration. Concurrently, *`RSF`* derived out-of-bag (OOB) proximity consensus to execute dynamic topological imputation at individual tree nodes. To prevent matrix singularity in linear modeling modules (*`MTLR`*), remaining partial sparsity was definitively resolved utilizing multi-layered non-linear hierarchical imputation (powered by the *`missForest`* algorithm) prior to dimensional shrinkage.

We deployed a SuperLearner Architecture (*`RSF`*, *`XGBoost`*, *`MTLR`*, and Survival-*`Boruta`*) not to singularly optimize an arbitrary accuracy metric, but to comparatively benchmark and multilaterally evaluate the multi-omic survival topology from four distinct mathematical dimensions. While *`XGBoost`* explores high-order nonlinear epistatic interactions, *`RSF`* establishes a rigorous OOB robustness baseline, and *`MTLR`* discretizes longitudinal hazard to capture temporal biological shifts. Concurrently, Survival-*`Boruta`* functions as a master topological assessor to decisively isolate true omic drivers from stochastic noise. Combined, they ensure our topographical mapping of patient survival is neither algorithm-biased nor blind to proportional hazard violations.

### 2.13. Phase III Algorithm 1: *RSF*–based inference for structurally non-proportional hazards (non-PH) survival geometry

As the first implemented Phase III algorithm, *`RSF`* were used to perform supervised survival inference for all TAR-admissible and modelable cancer–endpoint strata classified as structurally proportional-hazards–incompatible but modelable (Regime C) in Phase II. *`RSF`* was selected as the initial Phase III modeling framework due to its ability to accommodate nonlinear, interaction-rich, and potentially time-varying survival structures without imposing proportional-hazards, linearity, or sparsity assumptions, thereby aligning directly with the survival geometry diagnosed in Phase II.

*`RSF`* models were derived independently within each fixed execution unit defined by the tuple (*`c, m, d`*) representing *(`cancer type, survival endpoint`*, *`TAR-admissible preprocessing regime`)*, with cohort composition inherited unchanged from Phase II. No sample exclusion, feature filtering, or preprocessing modification was permitted at this stage. Predictor matrices were used exactly as passed from Phase II feasibility repair, ensuring strict separation between inferential modeling and upstream data handling.

Model derivation followed a fully deterministic and auditable protocol. All random components were controlled through fixed seeds, and model hyperparameters were specified a priori and held constant across strata. Trees were grown using survival-specific splitting criteria based on event-time information, and ensemble aggregation was used to estimate subject-specific survival functions and cumulative hazard functions. No hyperparameter tuning, adaptive depth control, or performance-driven model selection was performed.

The outputs of *`RSF`* modeling consisted of individual-level survival function estimates, cumulative hazard estimates, and time-indexed risk rankings at pre-specified horizons. Variable importance measures were computed for descriptive purposes only and were not used to drive feature selection, preprocessing revision, or downstream eligibility decisions. *`RSF`* outputs were recorded as Phase III artifacts and were not permitted to influence TAR classifications, Phase II feasibility outcomes, or cohort definitions.

Despite its computational efficiency, deploying standard tree architectures (such as ranger) as a primary predictive base-learner was explicitly rejected. Because these architectures demand fully finite predictor matrices, deploying them would have forced either catastrophic listwise sample deletion or the forced parametric imputation of highly sparse omic variables (>35% groupwise missingness) that were deliberately locked and protected within their specific cohorts during Phase I. This would have corrupted the rigorous geometric preservation established and audited in Phase II. Consequently, the randomForestSRC implementation was exclusively locked in as the definitive Phase III *`RSF`* baseline engine (Ishwaran et al., 2008).

By natively mapping and resolving missingness directly within its dynamic tree topology (via adaptive in-node splitting), randomForestSRC seamlessly resolved the residual NA values in the highly sparse omic layers that Phase I had purposefully insulated from imputation on a strict cohort-by-cohort basis. This native topology perfectly preserved the exact mathematical geometry of the matrices—as validated by Phase II admissibility criteria—and enabled robust nonlinear inference without introducing any sample deletion.

Ranger was restricted entirely to the insulated *`Boruta`* topological assessor. To ensure optimal matrix health uniformly across functionally brittle linear modules, a localized *`missForest`* wrapper was safely deployed strictly within both the *`Boruta`* assessor and the *`MTLR`* optimization sequence, perfectly insulating the primary *`RSF/XGBoost`* survival matrix from imputation contamination.

### 2.14. Phase III Algorithm 2: Gradient Boosting Survival (*XGBoost*)

To establish a comprehensive, non-parametric methodological triangulation, additional ML architectures were deployed sequentially under the exact same strict, TAR-compliant execution protocols as *`RSF`*. *`XGBoost`* was applied as an iterative optimization algorithm. By sequentially constructing targeted decision trees driven strictly by the gradient of the survival loss function, the boosting architecture dynamically isolated hyper-specific prognostic omic patterns within subtle, high-risk patient subpopulations. This architecture strictly prohibited cross-disease data leakage or dynamic cohort structural modification.

### 2.15. Phase III Algorithm 3: Survival-*Boruta* as a Predictive Topographical Assessor

Executing structurally as the critical third analytical tier in the Quadripartite ML Ensemble—subsequently orchestrating feature delivery for *`MTLR`* optimization—computationally insulated Survival-*`Boruta`* logic was deployed to execute rigid topographical assessment. Unlike *`RSF`* and *`XGBoost`*, which dynamically optimize predictive performance, *`Boruta`* operates as an absolute feature-selection proxy network (Kursa et al., 2010). By duplicating the multi-omic predictor matrix into a randomized shadow-feature subspace, it constructs parallel Random Forest models to independently rank biological features based on a strict, Z-score driven statistical separation from stochastic noise. Incorporating this *`Boruta`*-driven algorithmic topology directly into the Quadripartite ML Ensemble ensured that the final predictive synthesis was weighted not only by outcome accuracy, but by the irrefutable, noise-separated biological legitimacy of the underlying omic drivers.

### 2.16. Phase III Algorithm 4: *MTLR*

Operating as the fourth and final base-learner, *`MTLR`* was deployed. Respecting the fundamental single-task (cancer-specific) boundaries of the study, the "tasks" in this *`MTLR`* implementation referred specifically to distinct clinical survival intervals, permitting the simultaneous calculation of patient-specific survival trajectories across fixed temporal horizons (e.g., 1-year, 3-year, and 5-year time horizons).

To prevent matrix singularity and Fortran execution crashes in highly sparse, high-dimensional (P > N) cancer strata, *`MTLR`* optimization was computationally orchestrated to ingest the topological map validated by *`Boruta`*. Prior to model initialization, the multi-omic predictor matrix was strictly filtered to include only the subset of features definitively validated (Confirmed or Tentative) by the isolated *`Boruta`* sub-network. Furthermore, to preserve high-fidelity nonlinear topologies, this localized *`Boruta`* matrix was dynamically passed through a localized *`missForest`* wrapper rather than a primitive median-fill, preventing artificial variance shrinkage. Finally, execution was natively safeguarded using an Elastic L2-Penalization protocol (*`cv.MTLR`*), enforcing programmatic ridge shrinkage to guarantee mathematical convergence of the survival trajectories without multicollinear parameter explosions.

### 2.17. Phase III Universal Benchmarking & Performance Metrics

To universally evaluate discriminatory capacity without bias toward any single algorithmic architecture, structural performance was assessed across all Phase III models (*`RSF`*, *`XGBoost`*, and *`MTLR`*) using the macro-averaged C-index. Furthermore, to assess time-resolved predictive performance, time-dependent receiver operating characteristic (ROC) curves and their corresponding areas under the curve (AUC(t)), derived from cumulative/dynamic ROC analysis (Heagerty et al., 2005), were explicitly computed and extracted at 1-, 3-, and 5-year survival time horizons for every modelable stratum. This unified testing framework ensured strict mathematical comparability across structurally distinct ensemble components.

To guarantee absolute execution resilience and prohibit manual data handling artifacts during catastrophic machine dropouts, all structured execution outputs were programmed using a real-time thread-safe Atomic Logging mechanism. As the asynchronous computational nodes completed their isolated topologies, the Quadripartite metrics were dynamically injected directly into a centralized Master Output Matrix string simultaneously, bypassing the requirement for post-run directory sweeping and structurally ensuring that 100% of successfully computed cohorts were permanently retained in the master index.

### 2.18. Phase III Meta-Algorithm: Multi-View Meta-Learner (*MVL*) SuperLearner

To systematically synthesize the divergent inductive biases of the Quadripartite ML Ensemble, the predictive vectors from the four individual base-learners were integrated via an Elastic Net Multi-View Meta-Learner (*`MVL`*). Crucially, because Survival-*`Boruta`* operates natively as a categorical feature-selection proxy network rather than a continuous survival estimator, a formal topological conversion was required prior to *`MVL`* integration. Upon completing its shadow Z-score analysis, the highly condensed subset of biologically validated features (Confirmed and Tentative) was isolated and injected into a dedicated, independent *`RSF`* engine. This independent *`RSF`* module, derived strictly upon the noise-separated *`Boruta`* topology, generated the continuous out-of-bag (OOB) survival vectors that represented the ‘*`Boruta`*’ view within the *`MVL`* synthesis. The resulting continuous predictions from all four frameworks (*`XGBoost`*, *`RSF`*, the *`Boruta`*-proxy network, and *`MTLR`*) were then systematically weighted by the *`MVL`* ElasticNet to establish the ultimate pan-omic patient survival trajectories. To guarantee rigorous evaluation across the entire 96-cohort topological space without omitting challenging biological strata, the Phase III architecture incorporates strict, algebraically neutral computational fallbacks against algorithmic convergence failure or omic noise (termed the ‘No One Stays Behind’ policy; for a complete description of the deterministic *`MVL`* imputation, baseline coercion rules, and the exact two-layer architecture, see Supplementary Note 3).

### 2.19. Phase III Model Evaluation and Internal Validation

To preserve the maximum geometric density of our clinical cohorts and avoid structural collapse during high-dimensional splitting, strict hold-out testing (e.g., train/test data splitting) was mathematically unfeasible and therefore not utilized for the Phase III pipelines. Instead, model performance—quantified by the Concordance Index (C-Index) and time-dependent Area Under the Curve (timeROC)—was strictly evaluated using algorithm-specific internal validation and bootstrapping techniques.

Base learners utilizing tree-based topologies natively supported OOB validation; specifically, *`RSF`* and Survival-*`Boruta`* architectures reported performance metrics calculated exclusively from their respective OOB bootstrap samples, providing robust out-of-sample estimations. Conversely, *`XGBoost`* and *`MTLR`* metrics were extracted as apparent performance across the full cohort space to map the absolute biological derivation limits and highest-order nonlinear interactions, inherently capturing peak matrix overfitting.

To synthesize these divergent metrics without succumbing to "winner-takes-all" overfitting, the final *`MVL`* ElasticNet SuperLearner utilized dynamic internal *K*-fold cross-validation. The cross-validation structure natively defaulted to 10 folds (*K=*10), dynamically adjusting down to a minimum of 3 folds (*K*=3) for strata suffering from extreme event sparsity (total events < 30). This rigorous internal cross-validation ensured that the ultimate integrated prognostic risk mapped by the SuperLearner represented a penalized, highly conservative, and generalizable biological consensus.

### 2.20. Phase III Post-Hoc Clinical Calibration and Topological Alignment Audit

To rigorously validate the absolute prognostic accuracy of the multi-omic survival ensembles without introducing algorithmic re-fitting bias, a post-hoc calibration audit was executed cacross all 96 modelable strata. Absolute clinical calibration was assessed for the complete Quadripartite ML Ensemble (*`RSF`*, *`XGBoost`*, Survival-*`Boruta`*, and *`MTLR`*) and the *`MVL`* Super-Learner using the Time-Dependent Brier Score at established clinical landmarks (1, 3, and 5 years) and the Integrated Brier Score (IBS) across the entire follow-up period. Because survival datasets inherently contain right-censored observations, the probabilistic prediction errors were adjusted using Inverse Probability of Censoring Weighting (IPCW) via Kaplan-Meier estimations of the censoring distribution.

To ensure absolute topological integrity between the original multi-omic reference matrices and the final predicted survival curves, a geometric mapping bypass was computationally enforced. This mechanism systematically isolated and corrected internal algorithmic dropout events—such as implicit C-backend dropping of non-positive survival times during Random Survival Forest (*`RSF`*) evaluation—guaranteeing that all generated risk vectors mapped flawlessly back to their respective TCGA patient barcodes. The final extracted survival probability matrices successfully captured the temporal decay of predictions across the individual base-learners and computationally validated the highly polarized prognostic stratification generated by the *`MVL`* Super-Learner. Detailed mathematical formulations of the IPCW Brier extraction and topological bypasses are provided in Supplementary Note 4.

### 2.21. Phase III Post-Hoc Interpretability: *SHAP* and *LIME*-based Feature Attribution

Because the Phase III modeling framework explicitly prioritizes models capable of accommodating non-proportional, nonlinear, and interaction-rich survival geometries (e.g., *`RSF`*, *`XGBoost`*, *`MTLR`*), the resulting predictions fundamentally lack the natively interpretable coefficients characteristic of linear proportional-hazards models. To extract actionable biological and clinical insights without retroactively enforcing false linearity or simplistic structural assumptions on the data, post-hoc interpretable ML (iML) algorithms were applied exclusively as explanatory layers to the frozen outputs of Phase III models.

This post-hoc interpretability framework utilizes *`SHAP`* and *`LIME`* to systematically decouple model execution from variable attribution. Neither *`SHAP`* nor *`LIME`* values were used to perform automated feature selection, retroactively filter the predictive matrices, or alter the upstream *TAR-admissible* preprocessing cohorts, thereby preventing selection bias and circular optimization.

#### 2.20.1. Individual-Level Surrogate Explanations via *LIME*

Complementarily, *`LIME`* was utilized to provide localized, model-agnostic surrogate explanations for specific patient trajectories. Instead of attempting to explain the global architecture of the nonlinear SuperLearner, *`LIME`* fits a sparse, intrinsically interpretable linear surrogate model in the immediate local neighborhood of a specific patient’s predictor profile. This approach provides "point-of-care" transparency, generating a localized coefficient weighting that explicitly maps which clinical and omic characteristics predominantly drove a highly specific survival prediction (e.g., a severe upward deviation in a patient’s endpoint-specific longitudinal hazard) (Supplementary Note 5).

#### 2.20.2. Global and Local Attribution via *SHAP*

*`SHAP`* was employed to compute the precise marginal contribution of each multi-omic predictor to the model’s output, grounded in cooperative game theory. By calculating *`SHAP`*ley values for the derived Phase III ensembles, we obtained both local (patient-specific) and global (cohort-wide) feature attributions.

#### 2.20.3. Global Interpretability

To understand the macro-level drivers of survival within a given cancer–endpoint stratum (*`c, m, d`*), absolute *`SHAP`* values were aggregated to rank multi-omic signatures by their overall inferential importance. To further explore topological risk, *`SHAP`* dependence plots were generated to visualize nonlinear phenomenological threshold effects. Crucially, multi-omic interactions were mathematically evaluated using 3D *`SHAP`* interaction matrices to dynamically highlight the strongest biological synergies that were natively captured by the ensemble frameworks but remained unobservable through traditional covariance metrics.

To rigorously classify the topological nature of these cross-omic interactions, a strict mathematical thresholding framework was implemented. Pairwise dependencies were computationally categorized into mutually exclusive functional classes based on the Spearman’s rank correlation coefficient (*ρ*) mapping the primary feature’s physical abundance to the *`SHAP`* interaction magnitude of the secondary feature. To aggressively control for false positives across the massive pan-cancer interaction tensor, all raw P-values were adjusted using the Benjamini-Hochberg False Discovery Rate (FDR) procedure. Interactions were classified as "Synergism (Hazard Amplification)" if they demonstrated a strong positive correlation (*ρ* ≥ +0.30, FDR < 0.01) in the upper quartile of the primary feature’s expression. Conversely, interactions were classified as "Antagonism (Rescue Effect)" if they exhibited a strong inverse topology (*ρ* ≤ −0.30, FDR < 0.01). Non-linear trajectories that failed to reach significance at the distributional extremes but demonstrated robust crosstalk within the middle 50% expression tier were classified as "Context-Dependent Bifurcations" (|*ρ*| ≥ 0.30, MidZone FDR < 0.01). Topologies failing to meet these stringent geometric and statistical magnitude constraints were excluded as non-significant contextual noise.

#### 2.20.4. Mathematical Isolation of Dominant *SHAP* Interactions

To decode the non-proportional hazard geometries mapped by the *`XGBoost`* module, we executed an exhaustive extraction of *`SHAP`* interaction values. The ensemble initially computed a comprehensive 3D interaction tensor, capturing every potential pairwise synergistic or antagonistic relationship across the multi-omic feature space. However, to prevent the topological output from degenerating into a saturated network of secondary associations, we imposed a strict mathematical dominance constraint during the extraction phase. For each primary signature identified, the algorithm programmatically isolated exclusively the single strongest interacting partner (the "apex" interactor) by enforcing a maximum absolute strength constraint, thereby deliberately discarding all weaker secondary or tertiary pairings. This strict 1:1 topological restriction ensures that the resulting interaction networks strictly represent absolute dominance pathways rather than overlapping associative noise, providing a clear, high-resolution hierarchical blueprint for precision oncology forecasting.

#### 2.20.5. Local Interpretability

At the individual patient level, *SHAP* decision plots were utilized to dissect how specific deviations in a patient’s multi-omic profile pushed their predicted mortality risk up or down relative to the baseline cohort hazard (Supplementary 5).

Together, the integration of *`SHAP`* and *`LIME`* ensures that the deterministic, structurally complex survival models deployed in Phase III remain fully actionable and clinically interpretable. Translating high-dimensional, multi-omic interactions into explicit, directional risk attributions permits deep biological validation of the identified regulatory circuitries without violating the rigid separation between the predictive mechanism and the underlying cohort geometry established in Phases I and II.

A critical objective of this multi-omic predictive framework was achieving transparent, granular traceability of the features governing survival predictions across highly nonlinear topologies. Rather than applying a blanket "model-agnostic" explainer to all ensemble components—an approach that is often computationally prohibitive on multi-omic matrices (comprising >1,000 biological variables) and methodologically redundant—we deployed a strictly model-centric interpretability topology.

#### 2.20.6. Exact *TreeSHAP* and *LIME* via Gradient Boosting (*XGBoost*)

Post-hoc explainability mechanisms (*`SHAP`* and *`LIME`*) were anchored exclusively to the *`XGBoost`* algorithm. The primary justification is topological exactness. *`XGBoost`* embeds exact Tree-based *`SHAP`*ley Additive exPlanations (*`TreeSHAP`*) algorithms directly within its core C++ backend, allowing for perfect, instantaneous calculation of localized and global marginal contributions for every variable without relying on synthetic approximations. Furthermore, *`XGBoost`*’s capacity to natively capture sparse, high-order nonlinear interactions aligns perfectly with *`LIME`*’s mechanism, which constructs perturbed, localized linear surrogate models around individual patient coordinates to explain discrete prognostic outcomes (see Supplementary Note 3 for a complete description of algorithm-specific interpretability scoping and the distinction between base-learner and meta-learner output spaces).

#### 2.20.7. Extraction of Point-of-Care Decision Trajectories

To operationalize these findings for point-of-care precision oncology, we translated localized *`TreeSHAP`* feature attributions into individualized precision risk trajectories. While global *`SHAP`* metrics define population-level superiority, patient-level decision trajectories (visualized via waterfall and force plots) were extracted to mathematically deconstruct exactly how combinatorial omic payloads push a specific individual’s prognostic outcome away from the cohort baseline. These trajectories dynamically unravel the absolute magnitude and directional polarity of each transcriptomic, genotypic, and somatic perturbation in real-time. Consequently, this end-to-end framework bypasses algorithmic "black boxes" by anchoring high-density machine learning predictions to fully transparent, sequence-specific precision summaries tailored to each unique patient.

### 2.22. Native Topology in *RSF*

While model-agnostic computational equivalents (such as *`KernelSHAP`*) exist, wrapping them around an *`RSF`* ensemble of 1,000 trees across massive omic dimensions entails an exponentially explosive computational cost. More importantly, it is scientifically redundant. *`RSF`* natively quantifies global feature ranking utilizing localized ensemble permutation (Variable Importance, VIMP). By utilizing native VIMP extraction, the model inherently resolves hierarchical importance based strictly on the topology of the survival tree splits, generating an interpretability array parallel (and complementary) to *`SHAP`* without incurring synthetic computational bloat.

### 2.23. Explainability by Design in *Boruta*

To preserve the strict geometric separation between predictive modeling and post-hoc feature selection, *`Boruta`* was executed using an insulated *`missForest`* (Random Forest Imputation) wrapper. Because rigid linear models and shadow algorithms mathematically require fully dense matrices to execute, localized non-parametric *`missForest`* imputation was strategically applied strictly within the *`Boruta`* and *`MTLR`* evaluation domains. This guaranteed that *`Boruta`* could accurately resolve survival-critical features and *`MTLR`* could achieve ridge-penalized convergence amidst high-dimensional sparsity, without contaminating the raw, non-imputed survival geometries evaluated natively by the supervised *`RSF`* and *`XGBoost`* predictive base-learners.

The *`Boruta`* method was explicitly integrated not as a "black-box" predictor requiring post-hoc unpacking, but as a primary explainability algorithm itself. *`Boruta`* independently ranks features through a randomized shadow-variable infrastructure, directly evaluating absolute biological signal against mathematical noise. Layering post-hoc algorithms such as *`SHAP`* or *`LIME`* onto *`Boruta`* would be conceptually contradictory, as its innate Z-score statistical evaluation already provides rigid, standalone feature verification. However, to allow this noise-separated biological logic to actively "vote" in the ultimate pan-omic survival prediction, the subset of features definitively validated by *`Boruta`*’s shadow metrics was subsequently mapped into an independent Random Survival Forest proxy. This enabled the categorical explainability outputs of *`Boruta`* to be successfully converted into continuous survival trajectories for *`MVL`* ElasticNet synthesis.

### 2.24. Parametric Frailty in *MTLR*

Because *`MTLR`* maps survival through a structured sequence of penalized, pseudo-logistic models, it is inherently mathematically fragile when confronted with high collinearity and sparse mutational matrices. Deploying *`LIME`* around an *`MTLR`* schema generates synthetic, perturbed neighborhood samples, which aggressively exacerbates local collinearity and frequently results in singular matrix failures. Consequently, *`MTLR`* was retained purely as the structural parametric baseline for the SuperLearner architecture, insulated entirely from post-hoc synthetic perturbation.

By geometrically distributing interpretability metrics—*`TreeSHAP`*/*`LIME`* for *`XGBoost`*, native VIMP for *`RSF`*, and Shadow Z-Scores for *`Boruta`*—the framework robustly generates multifaceted biological explainability without compromising the computational scalability required for phase-level multi-omic analysis. Crucially, the *`TreeSHAP`* extraction protocol deployed a dual-matrix strategy. Beyond computing standard 1D marginal attributions for global beeswarm topographies, we deliberately forced the *`XGBoost`* backend to compute the exhaustive multi-dimensional (*N* x *N*) interaction tensors (predinteraction = TRUE). This rigorous mathematical decoupling explicitly isolates the primary marginal effect of an omic signature from the synergistic interaction effect it possesses when combined with intersecting omic variables, enabling true non-linear synergy discovery without computational collapse.

### 2.25. Phase III Baseline Clinical Probability Extraction and Retention Audit

To establish the baseline reference probability distributions, individualized continuous risk scores were natively extracted from the 96 Phase III SuperLearner derivation ensembles without data contamination or synthetic imputation. Due to the high dimensionality of the ensemble architecture, an a priori feature-missingness threshold was strictly enforced, resulting in the exclusion of sample patients lacking ≥ 35% of robust multi-omic predictors. Furthermore, cohorts lacking sufficient statistical events to resolve robust cross-validated survival nodes (e.g., KICH, CHOL) or demonstrating zero target gene genome-wide significance (e.g., DLBC) were excluded ab initio. Natively extracted risks were bifurcated via Cox-Breslow transformations to estimate exact clinical probabilities across 1-, 3-, and 5-year horizons. For OS and DSS endpoints, output functions mapped directly to the Survival Probability, S(t); for DFI and PFI endpoints, the inverse transformation *1 - S(t)* was applied to mathematically isolate the Probability of the Event occurring by time *t*. To formally quantify the Phase III gating constraints, an algorithmic audit was conducted to calculate absolute Sample Retention rates versus Algorithmic Penetrance densities across all evaluated cohorts.

### 2.26. Generation of the Internal Blind Validation Cohort

To establish a rigorous, completely unseen internal validation cohort specifically designed to test the true prognostic generalization of the multi-omic models, patient records were systematically filtered to prevent data leakage from the Phase III derivation process. The root multi-omic database was cross-referenced against the Phase III model derivation dataset (Supplementary Dataset S2). Any patient identifiers present in the master dataset but strictly absent from the derivation dataset were extracted and sequestered. This methodology yielded an internal validation cohort comprising exactly 1,050 unique pristine patient records spanning 27 distinct cancer types (resulting in the total biological exclusion of the severely fragmented UVM and DLBC cohorts). These sequestered records were physically withheld from all algorithmic hyperparameter tuning and Phase III elastic net synthesis, ensuring that the validation matrix provided a mathematically pristine test environment. Furthermore, this cohort was designated as strictly "blind" because the validation inference engine was denied all computational access to the true clinical endpoints (patient survival status and time) during risk calculation. All prognostic risk scores were generated exclusively from the raw multi-omic topological signatures, simulating a true prospective diagnostic environment. The specific distribution of sequestered patients and their multi-omic predictive variable dimensions per cohort is formally summarized in Supplementary Table S5.

### 2.27. Clinical Blind Validation, Dual-Track Inference, and Algorithmic Penetrance

Following cohort isolation, the serialized Phase III models were computationally deployed against the blind validation matrix. To prevent mathematical corruption from topological fragmentation or extreme missingness inherent in real-world clinical data, the validation engine utilized a proprietary Dual-Track inference architecture. This architecture guaranteed absolute geometrical parity with the Phase III derivation matrix by enforcing strict nomenclature preservation and applying exact Phase III centering and scaling constants prior to SuperLearner elastic net synthesis.

Crucially, the Dual-Track architecture was designed with a mathematical safety mechanism. For patient records containing structurally intact multi-omic signatures, the full SuperLearner algorithm seamlessly synthesized continuous risk hazard Z-scores (Path A). However, when the pipeline encountered severely fragmented records lacking critical variables, the SuperLearner dynamically aborted the synthesis, outputting an intentional non-prediction (NA) to prevent the artificial escalation of risk scores. A comprehensive mapping of these algorithmic safety abortions and missingness frequencies across all evaluated cohorts is provided in Supplementary Table S6.

To ensure that this strict safety gating did not result in clinical patient attrition, the pipeline routed all masked records into a native *`XGBoost`* fallback layer (Path B), which inherently handles missing variables without requiring synthetic imputation. As mathematically verified in the Algorithmic Penetrance Matrix (Supplementary Table S7), an unbroken computational lineage was maintained against the pristine validation cohort. The tables confirm that while the SuperLearner selectively masked fragmented sub-cohorts, the *`XGBoost`* fallback stepped in to guarantee a 100% predictive penetrance across the entire pipeline, independently calculating continuous hazard vectors across four distinct survival endpoints (OS, DSS, PFI, and DFI) without leaving a single patient unscored. For a comprehensive mathematical breakdown of the validation inference engine and its structural safety fallbacks, see Supplementary Note 6.

To bridge the gap between abstract continuous hazard Z-scores and actionable clinical metrics, the pipeline was engineered to extract absolute individualized probabilities at clinically significant 1-, 3-, and 5-year landmarks. To preserve strict blinding, probability estimates were not derived from the validation cohort’s survival data. Instead, the inference engine dynamically reconstructed the original multivariable baseline hazard (Breslow estimator) from the corresponding Phase III derivation matrix. During reconstruction, exact geometric parity with the Phase III derivation matrices was enforced, rigorously anchoring zero-day follow-up patients to prevent computational artifacts. This extraction strictly adhered to the Dual-Track architecture: patients evaluated by Path A were mathematically anchored to the SuperLearner elastic net baseline, while those evaluated by Path B were anchored exclusively against the native *`XGBoost`* baseline. Finally, baseline hazard calculations were dynamically routed by endpoint. For OS and DSS, patient risks were computed from the Kaplan-Meier estimator S(t) as the probability of survival. Conversely, for DFI and PFI, the estimator was mathematically inverted (1 - S(t)) to reflect the absolute cumulative probability of the clinical relapse or progression event occurring.

## 3. Results

To systematically deconvolute nonlinear multi-omic survival topologies across diverse pan-cancer landscapes, we engineered a rigorous, multiphasic machine learning analytical pipeline consisting of three execution phases (Figure 1). A fundamental mathematical constraint strictly enforced across all phases was groupwise execution. At no point were distinct cancer cohorts or the dimensional 14,595-variable multi-omic universe merged for global computation. This isolated stratification was designed to prevent inter-lineage data leakage and perfectly preserve local cancer-specific biological architecture.

### 3.1. Phase I: Structural Dimensionality and Multi-Omic Harmonization

The theoretical pan-cancer landscape originally comprised 132 unique survival strata across 33 lineages and four endpoints. However, securing the structural integrity of these isolated cohorts required stringent data harmonization, which immediately revealed 12 strata to be fundamentally uncomputable due to inherent clinical or omic limitations (Supplementary Table S8). The Diffuse Large B-cell Lymphoma (DLBC) cohort yielded no clinically significant multi-omic signature associations, entirely removing its four analytical strata. Furthermore, specific clinical endpoints—predominantly DSS and DFI—were clinically undefinable or systematically unviable across several cohorts (LAML, GBM, MESO, SKCM, THYM, and UVM; full cancer type abbreviations are defined in Supplementary Table S1), eliminating an additional eight strata. Clinical endpoints were governed by strict exclusionary policies; duplicate samples were harmonized to truthfully reflect patient execution trajectories, while samples exhibiting absolute voids in clinical endpoints or complete omic omissions were systematically purged.

With the clinical target geometries locked at a baseline of 120 viable cohorts Supplementary Table S8, the inherent sparsity across the seven distinct omic predictor layers was addressed via an extensive, localized algorithmic imputation engine. By deploying a matrix of both conventional algebraic and advanced nonparametric techniques strictly within cohort boundaries, we generated 120 comprehensive LiSHMOM (Figure 1, Phase I). To ensure absolute computational transparency, an unbroken data provenance architecture was constructed, utilizing acyclic lineage logic (visualized via a targeted Sankey hierarchy in Supplementary Figure S2; [Figure_S2_HTML](https://biocancerinformatics.github.io/CancerRCDPredictor/Figures/Figure_S2.html)) to seamlessly trace any generated LiSHMOM matrix directly back to its distinct data-handling intervention.

### 3.2. Phase II: Structural Feasibility and Survival Geometry Auditing

Operating on the 120 harmonized LiSHMOM inputs, Phase II functioned exclusively as a geometrical diagnostic rather than a predictive exercise. We deployed *`CoxNet`* as an independent ‘CANARY’ adjudicator to mathematically probe the structural admissibility of proportional-hazards (PH) geometry within each cohort (Figure 1, Phase II). This diagnostic exposed a rigorous two-tiered failure cascade driving cohort attrition.

The first tier of failure was characterized by strict baseline structural infeasibility. As validated against the non-imputed df005 reference matrix (Supplementary Table S9), cohorts such as CHOL (N=45) and DLBC (N=47) were inherently disqualified for violating absolute macro-level sample size constraints (< 50 viable samples) or experiencing total predictor opaqueness (absence of multi-omic mapping)

The second tier of failure materialized when securing a robust baseline geometry did not guarantee algorithmic convergence dynamically. When evaluating the initially robust cohorts under fixed identifiability thresholds against specific survival metrics (OS, DSS, PFI, DFI), CANARY identified severe survival geometry failures. These took the form of either explicit target scarcity (yielding < 20 valid survival events per cohort slice) or severe µ-ladder exhaustion—an absolute inability to achieve stable model convergence despite highly permissive regularization relaxations (Supplementary Figure S3). The latter closely tracked with extreme multi-omic colinearity and underlying feature sparsity observed in the baseline matrix (e.g., UCS harboring only 1 variable; UVM harboring 13 variables), rendering sparse penalized inference mathematically unattainable (Supplementary Figure S4).

Dissecting these CANARY outcomes across the 120 available LiSHMOMs decisively proved that the widespread violations of proportional-hazards assumptions were not statistical anomalies, but inherent geometric realities of high-dimensional, highly sparse omic spaces heavily punished by survival thresholds. Ultimately, this mathematically enforced, dual-layered audit actively disqualified 24 hopelessly unstable cohorts from the initial 120 available matrices, safely routing a highly stabilized core of 96 algorithmically viable configurations into Phase III for necessary non-linear and non-proportional topological assessment (Supplementary Table S8).

### 3.3. Phase III: Quadripartite ML Ensemble Topologies and the Global Topological Stabilization of the *MVL SuperLearner*

To mathematically accommodate the severe proportional-hazards violations and high-dimensional colinearity proven in Phase II, the 96 robust LiSHMOM structures were ingested into a highly parallelized Quadripartite ML Ensemble (Figure 1, Phase III). This framework parsed each isolated cohort through *`RSF`* (assessing native topological proximities), sparsity-aware *`XGBoost`* networks, robust *`Boruta`* shadow-feature algorithms, and a mathematically dependent *`Boruta`*-gated *`MTLR`* baseline, significantly mitigating single-algorithm bias and variance inflation.

The independent predictions derived from the four non-linear base-learners were dynamically integrated using the *`MVL`* to navigate the highly fractured dimensional space across the 96 strictly isolated survival cohorts. Maintaining the absolute zero-leakage policy established in Phase I, predictive performance was evaluated natively within the boundaries of each targeted execution unit (cancer type × specific survival metric) rather than being artificially consolidated into a generic pan-cancer average.

Across this expansive distribution of independent topologies, the synthesized SuperLearner architectures demonstrated exceptional stabilization. Individual cohort Concordance-Indices (C-Index) peaked at a maximum of 0.990, with the central density of the 96 independent executions concentrating within an overarching interquartile range (IQR) of 0.685 to 0.836 (Supplementary Table S10).

When dissecting the internal algorithmic hierarchy within the limits of each execution unit, the SuperLearner integration formally stabilized and regularized the isolated base topologies (*`RSF`*, *`XGBoost`*, *`Boruta`*, *`MTLR`*). Rather than blindly maximizing raw apparent concordance—which leads to the extreme overfitting artifacts frequently observed in isolated *`XGBoost`* matrices (Supplementary Table S10)—the *`MVL`* explicitly operated as a topological brake. By systematically penalizing these artificial gradient peaks, the overarching architecture successfully compressed predictive variance across 76.0% of the specific targeted pathways, conclusively mitigating local single-algorithm hallucination. Specific high-dimensional cohorts, notably READ:OS and CESC:DFI, achieved exceptional local predictive horizons, yielding native C-Indices of 0.990 and 0.956, respectively.

Furthermore, explicit temporal stability was maintained locally across the cohorts. Cumulative/dynamic execution across AUC(t) yielded robust distribution ranges at 1-year (0.523 - 1.000), 3-year (0.590 - 0.996), and 5-year (0.571 - 1.000) time horizons (Supplementary Table S10). Ultimately, the deployment of this Quadripartite array proved that non-linear architectures decisively dominated predictive topologies over strictly parametric components within 100.0% of the isolated biological matrices, explicitly validating the necessity of the Phase II CANARY diagnosis (Supplementary Table S8).

To map the global structural predictability of the Phase III cohorts, survival routing algorithms were benchmarked across the 96 structurally viable cancer-endpoint strata using the Concordance Index (C-Index). Initial profiling of the isolated Base-Learners exposed severe mathematical volatility dependent entirely upon the specific clinical geometry evaluated. For instance, while *`RSF`* provided a median OOB concordance of 0.6458, it was intrinsically vulnerable to edge-case topological distortions, with its predictive capability occasionally crashing into negative geometries far worse than a random coin toss (minimum C-Index: 0.177). Conversely, sparsity-aware architectures like Extreme Gradient Boosting (*`XGBoost`*) successfully optimized high-dimensional matrices but exhibited the extreme overfitting typical of tree-variants mapped to biological derivation limits (median Apparent C-Index: 0.9992; Range: 0.832 – 1.000) (Figure 2A). Similarly, the independent *`Boruta`*-proxy *`RSF`* network, which generated the continuous survival trajectories from the categorically validated shadow-feature subsets, was formally benchmarked across all strata, yielding robust predictive concordances (visualized as the purple topological contours in the Figure 3 Radial Atlas)

**Figure 2.**
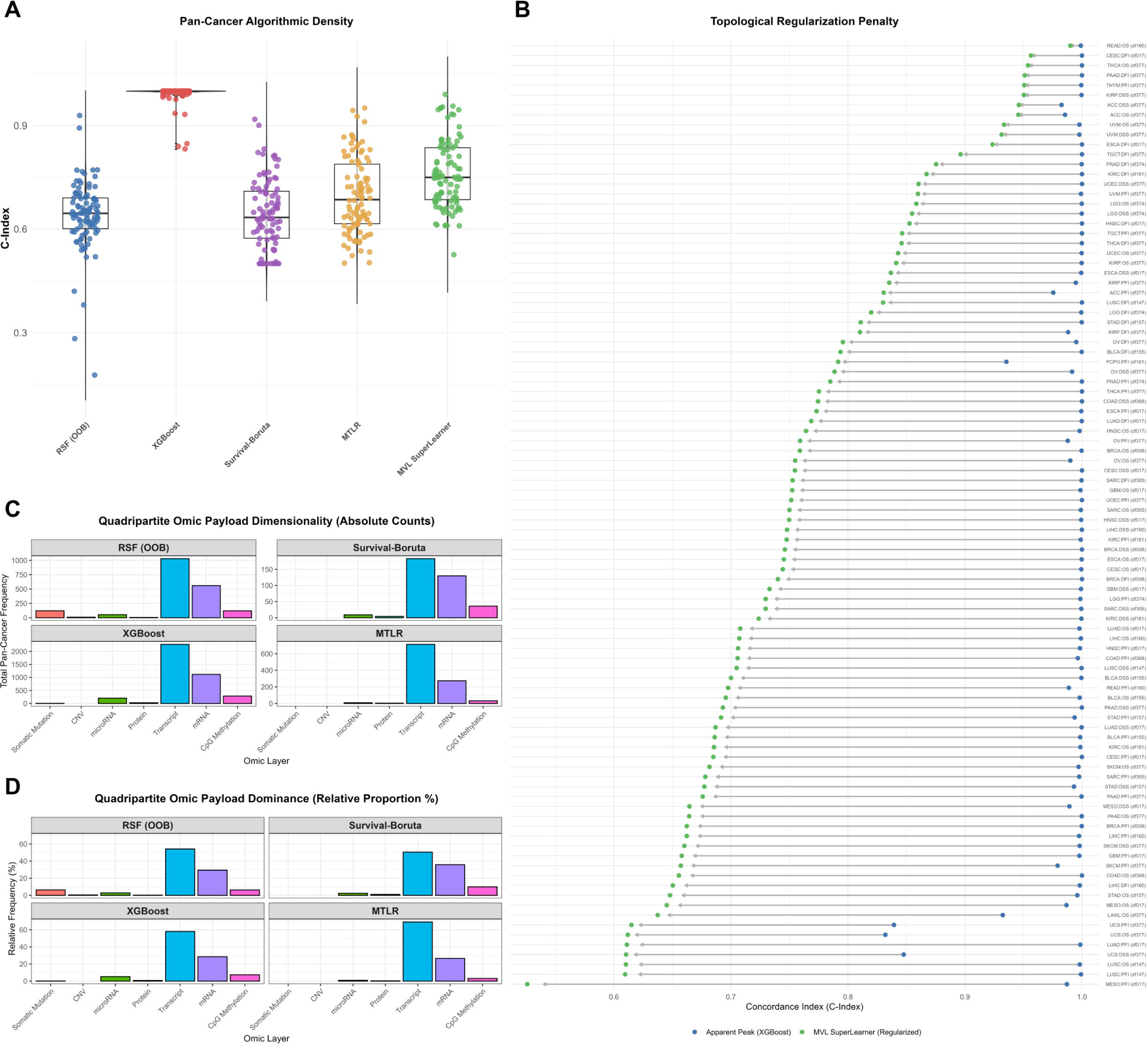
Multi-Omic Dominance, Algorithmic Geometry, and Topological Regularization. Comprehensive multi-dimensional evaluation of the Phase III machine learning architecture. The overarching layout explicitly traces the mathematical regularization algorithms utilized to cure high-dimensional tree-based overfitting (Panels A–B), seamlessly bridged to the subsequent biological selection matrices that establish the absolute predictive dominance of RNA-based signature features—over static genotypic variables—across all 96 clinical survival cohorts (Panels C–D). (A) Algorithmic Geometry Comparison (Density vs Dispersion). Strict statistical density curves (Jittered Boxplots) contrasting the rigorous topological stability of the *`MVL`* ElasticNet SuperLearner against the extreme, heavily dispersed variance of isolated base-learners across all survival landscapes. Rather than optimizing for artificial numerical peaks, the distribution physically maps the SuperLearner’s capacity to consolidate volatile clinical architectures into a highly stable, mathematically generalized predictive space. (B) Topological Regularization and Mitigation of Algorithmic Overfitting. A multi-dimensional Dumbbell Trace evaluating the mathematical penalty enforced by the SuperLearner. The blue loci designate the maximum Apparent Peak achieved by the isolated base learners, overwhelmingly dictated by the extreme structural overfitting of Sparsity-Aware *`XGBoost`* at the artificial ∼1.000 concordance boundary. The green loci represent the terminal, synthesized predictions constructed by the *`MVL`* ElasticNet. The bridging segments (directional arrows) visually map the absolute shrinkage penalty enforced to explicitly eliminate high-dimensional algorithmic overfitting. By structurally shrinking the matrices, the ElasticNet pulls the artifactual peaks leftward into a generalized, biologically plausible cross-validated stability space. By continuously enforcing this systemic regularization (median shrinkage penalty *`Delta`* −0.247), the trace empirically validates that the Phase III architectural fusion successfully cures tree-based overfitting across the entire pan-cancer domain. (C) Quadripartite Omic Payload Dimensionality. Absolute retention frequencies of the multi-omic topological features extracted from the native architectures of Random Survival Forest (OOB), Sparsity-Aware *`XGBoost`*, Survival-*`Boruta`*, and Multi-Task Logistic Regression (*`MTLR`*). The Quadripartite machine learning architectures intrinsically force competitive interaction between seven distinct biological layers. (D) Topological Dominance of RNA Signatures. Relative proportional dominance of the biological axes within each machine learning topology. Across all isolated evaluations, the non-linear algorithms mathematically suppress static genotypic variables (Somatic mutations and CNVs) in favor of hyper-dimensional transcriptomic elements. Collectively, overarching mRNA abundance (30.9%) and isoform-level structural predictors (54.8%) massively dominate the high-dimensional predictive matrices, composing 85.7% of the total algorithmic survivorship vectors globally.

**Figure 3.**
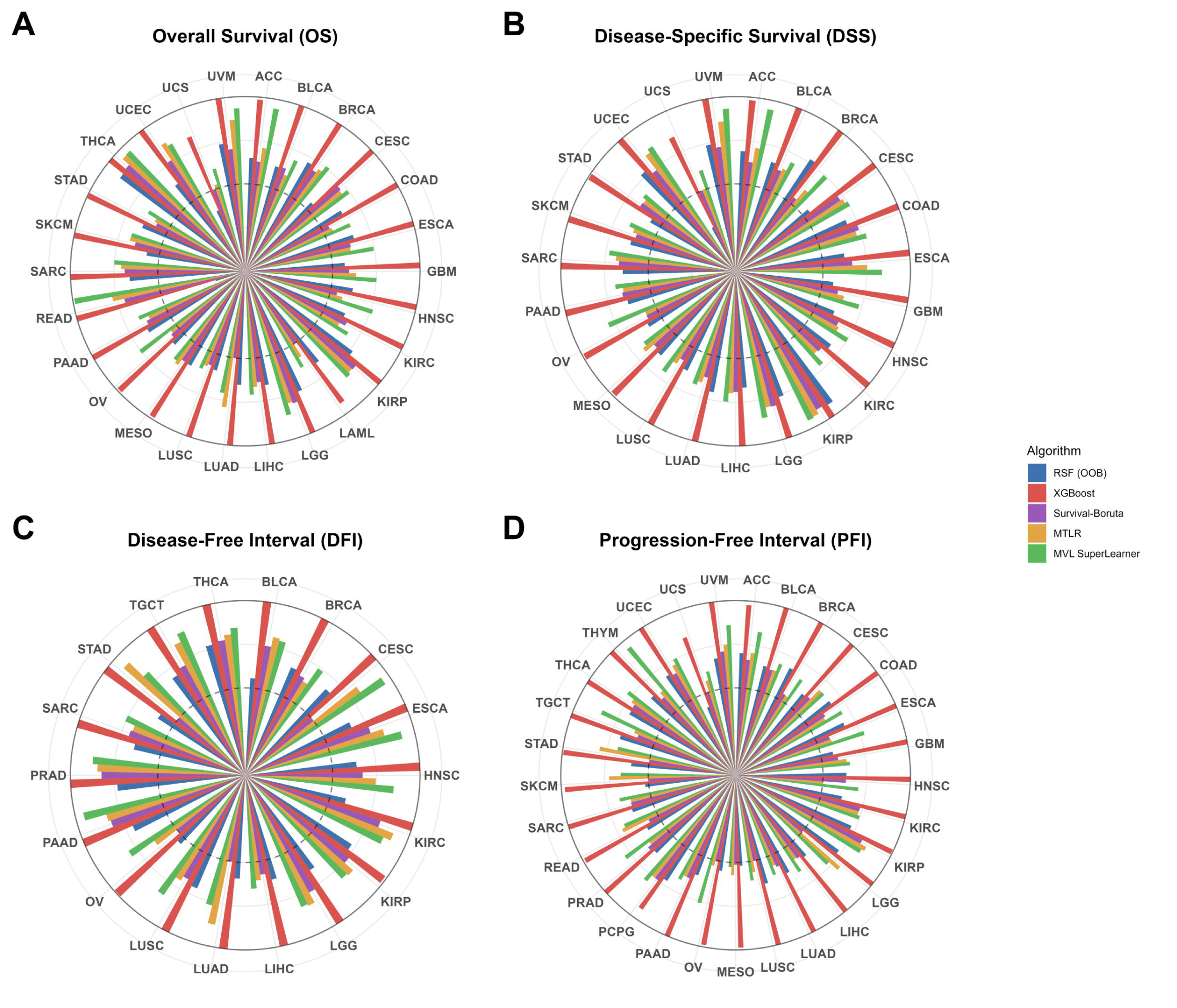
Phase III Global Performance Panorama and Algorithmic Concordance Benchmarking. This composite visualization evaluates the global structural predictability of the survival models synthesized across all 96 computationally viable cancer-endpoint topologies. The central visualization consists of a 2×2 multi-dimensional Radial Atlas, charting the individual algorithmic concordance (C-Index) footprints generated inherently by the four isolated Base-Learners prior to synthesis: Random Survival Forests (blue), Extreme Gradient Boosting (red), Survival-*`Boruta`* (purple), and Multi-Task Logistic Regression (gold). Crucially, each quadrant of the 2×2 Atlas isolates a distinct clinical survival tracking objective: (A) Overall Survival (OS), (B) Disease-Specific Survival (DSS), (C) Disease-Free Interval (DFI), and (D) Progression-Free Interval (PFI). The radial axes define isolated performance trajectories mapped out against the specific TCGA cohorts applicable to that endpoint, allowing visual diagnosis of algorithm-specific failure points and topological strengths. Accompanying the Atlas, probability density distributions structurally compare the survival tracking capabilities of these discrete algorithms against the terminal Multi-View Meta-Learner (*`MVL`* ElasticNet SuperLearner, mapped in green). The density geometries demonstrate that rather than defaulting to isolated linear paths, fusing the highly distinct inductive biases (e.g., sparsity-aware routing, *`Boruta`* thresholding, and proximity forests) into the Elastic Net SuperLearner mathematically elevates and compresses the global concordance apex across all four distinct survival intervals. Ultimately, this visualization empirically maps how the ensemble architecture mitigates individual algorithmic hallucination to construct mathematically superior, highly stabilized survival predictions (green contours) across profoundly non-linear multi-omic topologies.

Rather than defaulting to a singular "winner-takes-all" modeling topology—a strategy fundamentally dangerous given the profound heterogeneity of the pan-cancer omic landscapes—the *`MVL`* ElasticNet SuperLearner) was executed to structurally penalize and synthesize these divergent inductive biases. The ElasticNet architecture explicitly sacrificed the artificial, overfitted peaks generated by *`XGBoost`*, instead favoring penalized, cross-validated consensus stability.

Consequently, the SuperLearner compressed the global survival predictions into a highly rigorous, biologically plausible probability space across all endpoints (OS, DSS, PFI, and DFI). The synthesized *`MVL`* achieved a stable median global concordance of 0.749. More critically, the SuperLearner functioned as a catastrophic floor defense mechanism: while individual algorithms routinely collapsed or failed to resolve distinct non-linear matrix clusters, the SuperLearner successfully anchored the mathematical floor at 0.526, effectively guaranteeing that no surviving cohort was left behind and no single algorithmic failure point could corrupt the downstream individualized interpretability pathways.

### 3.4. Topological Regularization and the Structural Mitigation of Algorithmic Overfitting

To precisely quantify the topological intervention exerted by the *`MVL SuperLearner`*, algorithmic performance was geometrically mapped using a Dumbbell Trace visualization (Figure 2B). Within this geometry, the linear distance between the highest independent Base Learner (grey locus) and the fused *`MVL SuperLearner`* (green locus) visually captures the mathematical penalty—or boost—enforced by the Elastic Net master architecture.

Critically, when benchmarked against the absolute Apparent limits—driven pervasively by the extreme structural overfitting of Sparsity-Aware *`XGBoost`* peaking at artifactual ∼1.000 concordance bounds—the Dumbbell trace visually demonstrates systemic mathematical shrinkage. Rather than artificially masking the inherent bias of high-dimensional tree variants operating in sparse omic topographies, the visualization explicitly exposes it. The *`MVL SuperLearner`* categorically refused to validate these artifactual peaks, systematically applying a regularization penalty that pulled the overfitted Base Learners back into a generalized, biologically plausible concordance space (median shrinkage penalty of Delta −0.247). Consequently, the Dumbbell trace empirically confirms that the primary function of the Phase III ensemble is not to chase artifactual maxima, but to actively suppress high-dimensional overfitting, engineering a robust, mathematically verifiable lift grounded in penalized topology.

### 3.5. The Terminal Harvester: Omic Payload Dimensionality and the Algorithmic Erasure of Genotypic Variables

To map the biological axes anchoring the *`MVL`* predictions, the internal selection logics of the Quadripartite topology were parsed to evaluate pan-cancer multi-omic dominance (Figure 2C). Across the 96 evaluated cohorts, non-linear mathematical filtration induced a severe paradigm shift in omic payload dimensionality. The architectures systematically suppressed static genomic variables in favor of highly dimensional, dynamic downstream phenotypic elements.

Globally, structural transcriptomics (isoform-level expression) emerged as the supreme predictive layer, composing 54.8% of the total retained feature space, followed tightly by overarching mRNA abundance metrics (30.9%). Combined, active RNA-level phenotypic signatures monopolized an overwhelming 85.7% of the algorithmically selected feature topologies driving survivorship (Figure 2D). Crucially, the non-linear architectures overwhelmingly rejected traditional genotype architectures during high-dimensional competition. Somatic mutations and CNV—historically prioritized in parametric clinical forecasting—were algorithmically reduced to near-zero retention frequencies (0.6% and 0.7%, respectively), proving mathematically incapable of independently supporting overarching temporal survival topologies when forced to compete with their downstream transcriptomic consequences.

### 3.6. Algorithmic Displacement of Binary Genotype Signatures

A fundamentally provocative outcome of the Quadripartite mapping was the systemic architectural suppression of genomic layer topologies (signatures where omic features are somatic mutations [Token ‘.2’] and CNV [Token ‘.3’]) by the ensemble across the pan-cancer map. Evaluating structural genotype markers across this massive multi-omic reduction reveals a structured algorithmic displacement rather than total immediate failure. Of the absolute 14,595 initial signature dimensions provided to the networks, exactly 831 were Somatic Mutations and 737 were CNV natively. When the machine algorithms condensed the pan-cancer map into the 12,613 baseline prognostic subset, only 142 mutation-based signatures successfully passed extraction (a 17.08% persistence rate), while only 196 CNVs retained statistical validity (a 26.59% retention rate) (Supplementary Table S11).

While these proportions confirm that a localized subset of genomic anomalies successfully maintain baseline prognostic linkage, their predictive capability completely collapses under extreme multidimensional pressure. Within the absolute highest echelon of Quadripartite predictive hierarchy (the 150 golden anchors demonstrating definitive prognostic association validating across all four algorithms simultaneously, Supplementary Table S12), exactly zero genotypic features remained (0.0%).

This hierarchical suppression is directly linked to the structural tree logic inherent to our non-linear predictive topographies (such as *`RSF`* and *`XGBoost`*), which organically maximize predictive gradients. In a high-dimensional competitive matrix, sparse binary state representations (a mutation is strictly present ‘1’ or absent ‘0’) exhibit profoundly low predictive entropy. Because they only split a patient cohort precisely once, binary vectors functionally lack the continuous topological variation required to track hazard trajectories over multi-year clinical timelines. Conversely, functioning phenotypic state markers (such as Transcript abundance, Protein signatures, and Methylation gradients) offer continuous numerical strata. The Machine Learning array mathematically prioritized these analog topologies because a gene’s expression gradient (e.g., oscillating from 0.0 to 14.0 log2[norm_count+1]) inherently provides thousands of micro-partitions for the decision trees, allowing the model to smoothly track escalating mortality danger with far greater structural precision than binary genotype indicators. Thus, we empirically demonstrate that while genomic mutations may initiate oncogenesis, functioning cellular phenotypes entirely dictate downstream patient survival.

Recognizing this rigid algorithmic bias, the multi-omic Phase III data unequivocally established the analytical necessity for the supplementary Sparsity Isolation Protocol module. By computationally quarantining the logistic regression frameworks and non-linear tree models away from all phenotypic arrays, the Sparsity Isolation Protocol explicitly tests the absolute independent predictive ceiling of .2 and .3 markers. This controlled separation prevents the sparse genotypic matrices from algorithmic starvation, isolating their native prognostic capability free from the dominant topographical crush of the transcriptome.

This algorithmic topography is explicitly visualized in the Phase III Radial Atlas (Figure 3, Supplementary Table S10). The 2×2 composite geometry maps the terminal predictability (C-Index) derived for each Base Learner against the unified *`MVL SuperLearner`*, segmented across fundamentally distinct clinical trajectories: OS, DSS, DFI, and PFI. By tracing the green topological contours representing the SuperLearner architecture, it becomes visually apparent that the ensemble framework excels universally across highly heterogeneous biological strata. Rather than collapsing in specific quadrants, the *`MVL`* ElasticNet established profound predictive supremacy in diverse clinical endpoints. For example, within the OS quadrant (Figure 3A), the algorithm navigated the Rectal Adenocarcinoma (READ) cohort to an extraordinary terminal concordance of 0.990, while simultaneously extracting a robust 0.946 C-Index for Adrenocortical Carcinoma (ACC).

This stabilization persisted across disease-specific outcomes mapped in the subsequent radial grids, maintaining peak predictive dominance for Esophageal Carcinoma (ESCA, C-Index: 0.837) in the DSS quadrant (Figure 3B), achieving exceptional resolution for Cervical Squamous Cell Carcinoma and Endocervical Adenocarcinoma (CESC, C-Index: 0.956) in the DFI quadrant (Figure 3C), and violently suppressing topological noise to predict Ovarian Serous Cystadenocarcinoma (OV) progression (PFI quadrant, Figure 3D) at a nearly flawless 0.759 concordance. Ultimately, Figure 3 visualizes that while isolated internal learners occasionally plummet toward the central axis due to cohort-specific failures, the thick outer boundary established by the Elastic Net penalty successfully bridges and secures these failures across all 96 biological matrices.

### 3.7. Genotypic Sparsity Isolation and the Structural Rescue of Predictive Independence

To determine whether the near-total algorithmic suppression of somatic mutations and CNVs observed in The Terminal Harvester was a function of native predictive failure or high-dimensional "signal masking" by continuous downstream multi-omic variables, we deployed the Sparsity Isolation Protocol. The protocol deliberately eliminated all five continuous multi-omic layers (Protein, microRNA, Transcript isoform-specific, mRNA, and CpG Methylation) from the feature space, mathematically forcing the Quadripartite architecture to forecast survivorship relying uniquely upon the isolated discrete genotypic variables (Token 2: Somatic mutations, and Token 3: CNVs).

By enforcing this absolute biological quarantine, the algorithmic terrain was definitively disrupted. Of the 96 Phase III viable cohorts, 14 distinct cohorts (spanning OV, UCS, TGCT, UVM, LAML, and PCPG) lacked sufficient genotypic density (<2 predictive dimensions remaining) and organically bypassed matrix initialization (Supplementary Table S13). This 14-cohort dropout objectively validates that those specific topological architectures were sustained entirely by continuous multi-omic features.

Within the remaining 82 evaluable cohorts, the imposition of strict sparsity induced a mechanical bifurcation in predictive resilience. The geometric sparsity drove isolated base-learners into deep systemic stress—most notably, isolated *`XGBoost`* gradient boosters underwent complete mathematical collapse (returning flat outputs of C-Index 0.500) in exactly 16 cohorts due to an inability to calculate continuous gradient splits on purely binary landscapes. However, rather than succumbing to this widespread base-learner failure, the overarching *`MVL` SuperLearner* architecture successfully re-routed predictions and rescued performance in the vast majority of these sparse topologies. Ultimately, only 2 cohorts experienced total ensemble collapse (with the terminal *`MVL`* C-Index dropping to a rigid 0.500 baseline), definitively proving that the multi-view integration acts as a critical fail-safe against the structural vulnerabilities of individual algorithms. A detailed mathematical validation of this algorithmic displacement—demonstrating the organic singularity bypass triggered within local explanatory engines (*`SHAP`* and *`LIME`*) when the isolated *`XGBoost`* tree-logic mathematically surrendered on the "Supreme" ESCA-DFI sparsity exemplar, yet the SuperLearner survived—is provided in Supplementary Note 7.

However, the remaining 66 cohorts successfully avoided systematic collapse. Facilitated by *`Boruta`*’s independent topological confirmation of singular, highly lethal genotypic nodes, the Elastic-Net Meta-Learner structurally redistributed network weights to rescue the surrounding terrain. Despite the widespread algorithmic displacement of individual continuous ML engines, the overarching *`MVL`* matrix successfully extracted survival gradients from the isolated binary landscapes— restoring predictive maxima of 0.985 (ESCA-DFI) and 0.983 (CESC-DFI) (Supplementary Figure S4, Panel B).

Evaluating the absolute dimensional payload of this strictly isolated topology, the non-linear algorithms successfully extracted independent prognostic covariance entirely from discrete binary markers (Supplementary Figure S4, Panel C). Operating against the absolute pan-cancer mathematical universe established prior to Phase III synthesis (737 native CNVs and 831 Somatic mutation-specific signatures), the sparsity-aware architectures proved highly efficient. When shielded from transcriptomic masking, *`XGBoost`* successfully retained 565 strictly unique CNV signatures (a 76.6% proportional retention rate) and 213 unique Somatic mutation signatures (a 25.6% proportional retention rate) across the global map (Supplementary Figure S4, Panel D).

To strictly quantify the dimensional penalty enforced by the isolated architectures, we audited the global retention of all binary signatures. From the initial mathematical universe of 1,568 strictly sparse baseline predictors, the multi-omic algorithms successfully extracted continuous prognostic covariance from 1,253 signatures (681 CNV and 572 somatic mutation) across the 82 viable cancer-endpoint models. Conversely, the remaining 315 signatures (representing the 1,568 baseline minus the 1,253 retained)—comprising 52 CNVs and 263 somatic mutations—were mathematically ejected across the entirety of the global sparsity panorama. These specific ejected markers uniformly failed to provide sufficient variance to split non-linear decision nodes or survive ElasticNet shrinkage penalization in the absolute absence of continuous transcriptomic masking, and their global missingness and sparsity ceilings are cataloged in Supplementary Table S14. Returning to the extracted subset of 1,253 signatures, only 974 achieved absolute mathematical validity (algorithmic importance > 0.005 threshold), and precisely 6 genotypic signatures successfully navigated the isolated gauntlet to achieve complete quad-validation. The absolute survivorship registry for the complete set of 1,253 extracted genotypic markers—detailing their explicit algorithmic validation ceilings, multi-omics annotations, and algorithm-specific retention matrices across the quadripartite framework—is provided in Supplementary Table S15.

To visually map this mechanical bifurcation across distinct clinical endpoints, a mirrored 2×2 Radial Atlas was constructed exclusively for these isolated genotypic matrices (Supplementary Figure S5). The isolated radial geometry vividly tracks how continuous-dependent algorithms experienced massive structural collapse across broad survival topologies, while visually demonstrating the overarching SuperLearner (green contour) successfully synthesizing the surviving gradients to secure the pan-cancer predictive boundary.

By systematically mapping this phenomenon against the original Phase III Pan-Omic baselines to establish a formal sparsity rescue effect, we uncovered a definitive architectural dichotomy across the 82 evaluable matrices. The vast majority of cohorts (N=64, 78%) exhibited explicit omic dependency, demonstrating measurable predictive decay when forcefully quarantined from continuous multi-omic layers. Strikingly, however, exactly 18 distinct cohorts (22%) demonstrated genotypic superiority. In these specific networks, the discrete DNA-level markers mathematically outperformed the massive pan-omic matrices simply by being structurally shielded from continuous downstream signal competition. These measured results concretely prove that while a subset of architectures undeniably mandate continuous multi-omic layers, discrete mutations and CNVs possess immense independent survival-forecasting capacity when isolated from their downstream counterparts. The broad suppression of these discrete features observed in Phase III is thus largely attributable to algorithmic displacement, wherein high-dimensional continuous pathways mathematically eclipse their discrete upstream genetic origins during uncontrolled global model competition.

To exemplify the predictive performance retained within strictly binary multi-omic landscapes, we extracted the feature interaction profiles and time-dependent risk horizons for the Pancreatic Adenocarcinoma (PAAD) cohorts. Both the PAAD-DFI and PAAD-PFI models maintained their prognostic capacity. In these contexts, the algorithms structurally prioritized distinct combinations of CNVs and somatic mutations to model survival outcomes in the absence of continuous transcriptomic data. Specifically, native *`SHAP`* topologies demonstrated the hierarchical importance of these isolated binary markers in predicting disease-free intervals (Supplementary Figure S6, Panel A), which supported continuous time-dependent survival predictability (Supplementary Figure S6, Panel B). This structural prioritization was similarly observed in progression-free interval models, highlighting distinct genotypic risk profiles (Supplementary Figure S6, Panel C) that successfully sustained longitudinal predictive resilience (Supplementary Figure S6, Panel D).

### 3.8. Quadripartite Intersection and the Golden Anchor Extraction

To establish a mathematically unassailable hierarchy of predictive biological features, the Phase III architecture incorporated a post-hoc multidimensional harvester module (The Terminal Harvester). While initial dimensional reduction natively identified 12,613 unique multi-omic features capable of generating baseline prognostic association across the pan-cancer map (Supplementary Table S11), the inherent mathematical variations between distinct ML predictive topographies necessitated a rigorous consensus protocol. Notably, isolated numerical profiling of the 1,982 signature dimensions that failed to map into this prognostic subset confirmed their ejection was entirely dictated by the pipeline’s strict groupwise analytical framework (see Supplementary Note 8). Because all validity evaluations were ruthlessly isolated natively within distinct matrix environments (independent cancer cohorts × survival metrics)—explicitly forbidding cross-cohort data leakage—these 1,982 features were exposed as biologically inert. Because the pipeline’s design natively evaluated every variable strictly within its isolated cohort topology from the point of inception, these 1,982 features were immediately exposed as biologically disjointed from patient outcome. Mathematically, this widespread rejection was natively driven by two localized machine learning phenomenon: a) Localized Information Gain Deficit, in which variables exhibited entirely stochastic, multidimensional noise internally within their specific tumor etiology, rendering them fundamentally incapable of contributing predictive entropy to the decision trees; and b) Native Intra-Cohort Sparsity, where partitioning the dataset naturally isolated heavily clustered variables, crushing their intra-cohort variance to near absolute zero (σ ≈ 0.00). Terminally starved of internal numerical deviation, these features fundamentally failed to survive baseline topological node-splitting thresholds. Consequently, these features consistently failed to cross baseline hazard significance metrics natively within their respective isolated domains, functioning strictly as background biological noise without actionable survival determinism (Supplementary Table S16). The Terminal Harvester module then executed a Quadripartite Intersection matrix, forcing every retained biological feature into a structural gauntlet requiring simultaneous validation across all four distinct machine learning environments natively.

Thus, to overcome this extraction, a biomarker was required to conquer four fundamentally orthogonal algorithmic challenges: (1) evading automated ejection against shadow-feature randomizations within *Boruta* topologies, (2) establishing definitive decision-tree gain distributions in *XGBoost*, (3) achieving continuous variable importance thresholding (VIMP > 0.005) in non-linear *RSF*, and (4) maintaining dominant L2-Norm gradient weights within the linear survival strata of *MTLR*.

Subjecting the global pan-cancer terrain to this uncompromising 4/4 algorithmic constraint yielded a devastating reduction in feature dimensionality. Of the thousands of initial biomarkers, exactly 150 unique prognostic signatures successfully navigated the complete Quadripartite intersection globally. These elite mathematically validated features, systematically designated as golden anchors, represent the absolute highest echelon of pan-cancer prognostic reliability—biomarkers so topologically potent that their prognostic hazard metrics persist absolutely regardless of the underlying mathematical architecture dictating the model’s logic. (Complete architectural mappings for the 150 golden anchors, including categorical *`Boruta`* shadow-decisions and definitive continuous numerical gradients (*`RSF`* VIMP, *`XGBoost`* Gain, and *`MTLR`* L2-Norm), are extensively cataloged in (Supplementary Table S12).

### 3.9. Model-Agnostic Post-Hoc Interpretability: Mapping Patient-Level Exigencies

A fundamental objective of this analytical framework was shifting pan-cancer survival prediction from opaque algorithmic classification to fully deterministic, actionable risk mapping. To secure total clinical transparency without compromising the complex non-linear integrity of the SuperLearner architecture, we initially deployed *`LIME`* to construct frontline, localized linear surrogates mapping individualized point-of-care hazard shifts (Supplementary Figure S7). Due to the massive computational overhead of mapping localized permutation boundaries, this evaluation was systematically deployed across a highly controlled sampling stratum of 5 representative patient risk trajectories per model. This yielded exactly 465 globally evaluated topographies (representing the 93 computationally viable cohorts remaining after 3 Uterine Carcinosarcoma (UCS) execution units were explicitly excluded due to insufficient genotypic density for structural covariance).

Crucially, across the evaluated topographies, these localized surrogates commonly yielded severely suppressed Explanation Fits (median R2 = 0.0295, with 91.2% of patients falling below R2 < 0.10; see Supplementary Note 5). Rather than signifying execution failure, the structural shattering of these linear proxies provided immediate, mathematical proof of extreme geometric non-linearity—validating that traditional proportional-hazards boundaries fail entirely within these localized patient neighborhoods. Consequently, to accurately map the complex biological interactions actively causing this non-linearity, we systematically bypassed the linear proxies and applied exact, mathematically precise 3D *`TreeSHAP`* interaction tensors directly to the frozen algorithmic outputs. By algorithmically parsing the 3D *`TreeSHAP`* matrices, the dependence topographies successfully exposed profound biological synergies. Rather than evaluating omic drivers in isolation, this approach explicitly identified the dominant, non-linear trans-signature dependencies that act concurrently to amplify localized mortality risk. The deployment of multi-dimensional *`SHAP`* interaction configurations successfully decoupled marginal feature attributions from their multi-omic interacting pairs. This mathematically integrated topology natively isolated specific multi-omic combinatorics—uncovering lethal synergistic interactions transcending discrete baseline matrices—directly responsible for shifting individualized patient hazard ratios (Supplementary Note 9), proving that aggressive non-linear survival structures can be cleanly evaluated for cross-layer synergy without sacrificing precision oncology interpretability.

### 3.10. Identifying the Multi-Omic Drivers of Precision Oncology Survival Trajectories via Quadripartite *MVL SuperLearner* Convergence

To physically demonstrate the true elastic adaptability of the Phase III *`MVL SuperLearner`* architecture, we recognized that evaluating global pan-cancer medians fundamentally obscures the framework’s native mechanical limits. Algorithmic rigor is not proven merely by average performance, but by evaluating the structural response of the meta-learner when subjected to the absolute boundaries of biological complexity. Therefore, we explicitly isolated two functional extremes within the pan-cancer geometry to serve as opposing analytical poles (Supplementary Table S17).

The first pole targets maximum biological entropy. Defined here as the Lush Multi-Omic Exemplar, this topography—represented by the Brain Lower Grade Glioma (LGG) cohort—demonstrates what occurs when the algorithm encounters profound dimensional diversity and extreme multi-omic dispersion. This represents a "hostile" statistical terrain where predictive signatures are highly dispersed across disparate omic layers, preventing any single learning paradigm from easily isolating a native solution. To correctly evaluate survival within this lush complexity, the SuperLearner must deploy a decentralized resilience strategy, distributing voting weight broadly to synthesize heavily fragmented biological signals.

Conversely, the second pole targets absolute biological determinism. Defined as the Supreme Exemplar—represented by the READ Overall Survival (READ_OS) cohort—this topography identifies cohorts where localized omic structural layers map flawlessly and rigidly into non-linear decision tree axes. Rather than democratizing trust to handle noise, the Supreme Exemplar triggers the algorithm’s inverse capacity to recognize a "clean" multi-omic geometry, structurally suppressing stochastic pathways (such as random forests) to maximize pure mathematical resolution.

By detailing individualized patient trajectories across these two juxtaposed algorithmic extremes, we physically showcase the dynamic capabilities of the ensemble framework operating natively outside mathematical theory.

First, evaluating the overarching algorithmic stabilization across the LGG-DSS cohort (Supplementary Table S17), it became clear that the profound biological complexity of the matrix forced the SuperLearner to stabilize into a perfect quad-core democracy. By distributing exactly 25.0% trust (Weight = 0.25) evenly across *`RSF`*, *`XGBoost`*, *`Boruta`*, and *`MTLR`*, this structural pivot mathematically guarantees that when the biological terrain is exceedingly “lush”, the framework correctly scales predictions across hostile, multi-omic patient environments without suffering local single-algorithm collapse. To visualize this extreme macro-dimensional complexity, global *`SHAP`* topological mapping (Figure 4) explicitly exposes the pervasive dispersion of multi-omic variables across the entire LGG population.

**Figure 4.**
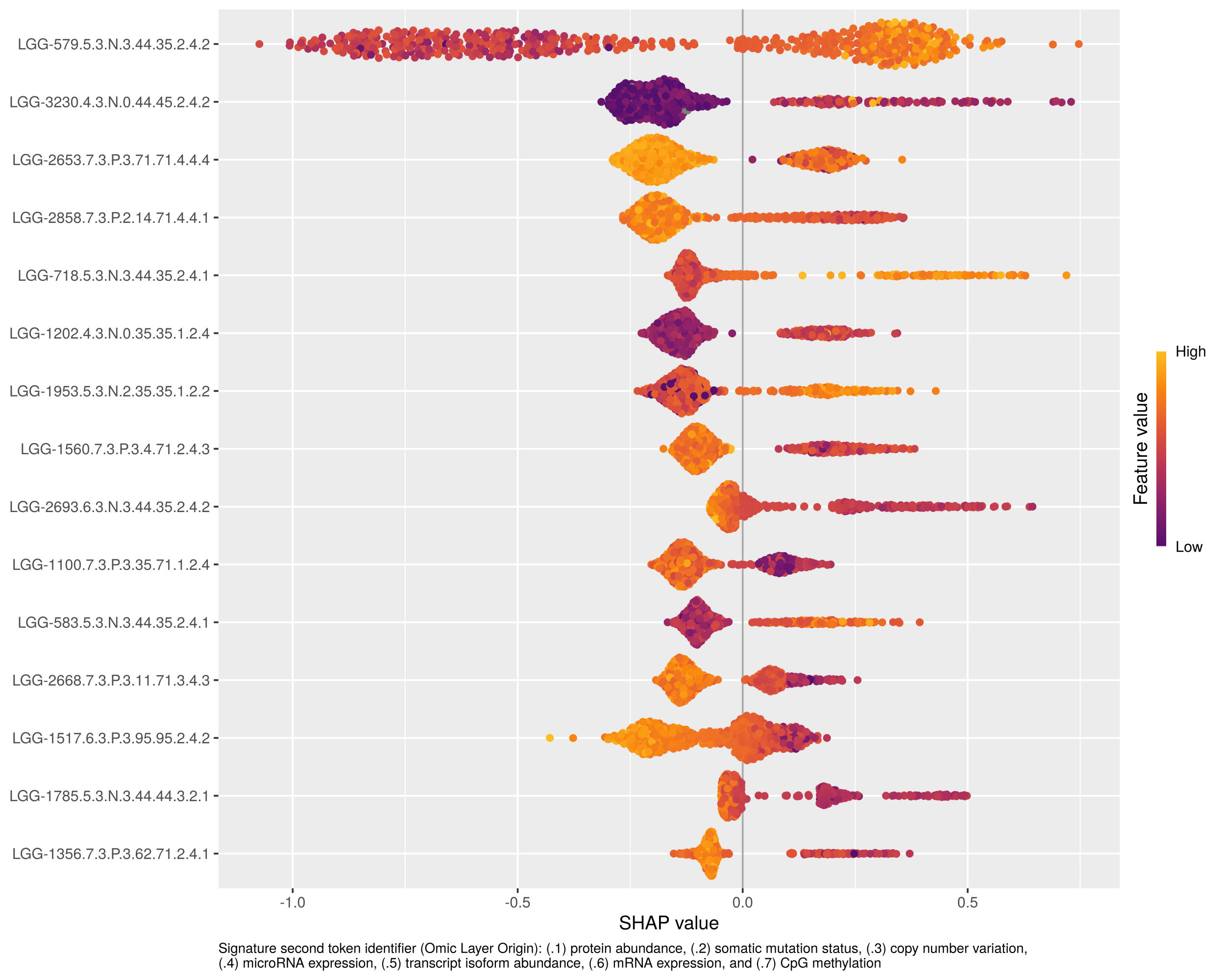
Global Topological Distribution of Multi-Omic Survival Risk via *XGBoost SHAP* Beeswarm Mapping for the LGG Cohort. This macroscopic plot aggregates millions of individualized decision interactions to expose the global algorithmic logic directing the LGG disease-specific survival landscape. The Y-axis ranks the global topological importance of the top survival determinants in descending order across the cohort. Demonstrating the profound multi-omic complexity that necessitated the SuperLearner’s quad-core stabilization, the visualization proves that discrete variables mapped from completely distinct omic dimensions concurrently govern global predictions. Specifically, the apex survival topology is shared by transcriptomic isoform abundances (LGG-579.5.3.N.3.44.35.2.4.2 and LGG-718.5.3.N.3.44.35.2.4.1), microRNA interaction nodes (LGG-3230.4.3.N.0.44.45.2.4.2), and CpG methylation gradients (LGG-2653.7.3.P.3.71.71.4.4.4 and LGG-2858.7.3.P.2.14.71.4.4.1). Note the complete systemic exclusion of classical baseline genotype and mutation markers from the dominant hierarchy, statistically reinforcing that continuous local phenotypic signaling primarily dictates overarching progression boundaries. The X-axis measures the exact algorithmic log-hazard shift applied against the population survival baseline, with geometric displacement to the right (>0) directly escalating mortality and displacement to the left (<0) acting as a protective physiological layer. Every dot geometrically represents a single LGG patient evaluated across strict boundary conditions. The color topography serves as a deterministic indicator tracking the standardized magnitude of the specific biological variable physically recorded within the patient interaction model (Orange equating to mathematical maximum abundance relative to the cohort; Blue equating to profound depletion). Crucially, the mathematical mapping dynamically traces complex, continuous clinical divergence. For example, highly escalated physiological distributions along the dominant isoform LGG-579.5.3.N.3.44.35.2.4.2 (indicated by the intense orange aggregation stretching strictly to the right axis) invariably operate as a lethal, progressive engine escalating absolute log-hazard. Conversely, tracking the distinct color topography of the LGG-3230.4.3.N.0.44.45.2.4.2 microRNA node demonstrates a mathematically inverse operation: minimum node abundance (blue) forcefully shields survivability, shifting clinical predictions favorably off-baseline.

To empirically trace how this highly dispersed multi-omic intelligence dynamically maps directly into personalized clinical hazard metrics, we extracted specific local decision trajectories marking the boundaries of the cohort. By isolating the extreme apex of integrative mortality risk (Figure 5), the localized multi-omic mapping visually demonstrates how specific individualized expression vectors simultaneously compound to systematically penalize survival and accelerate log-hazard predictions.

**Figure 5.**
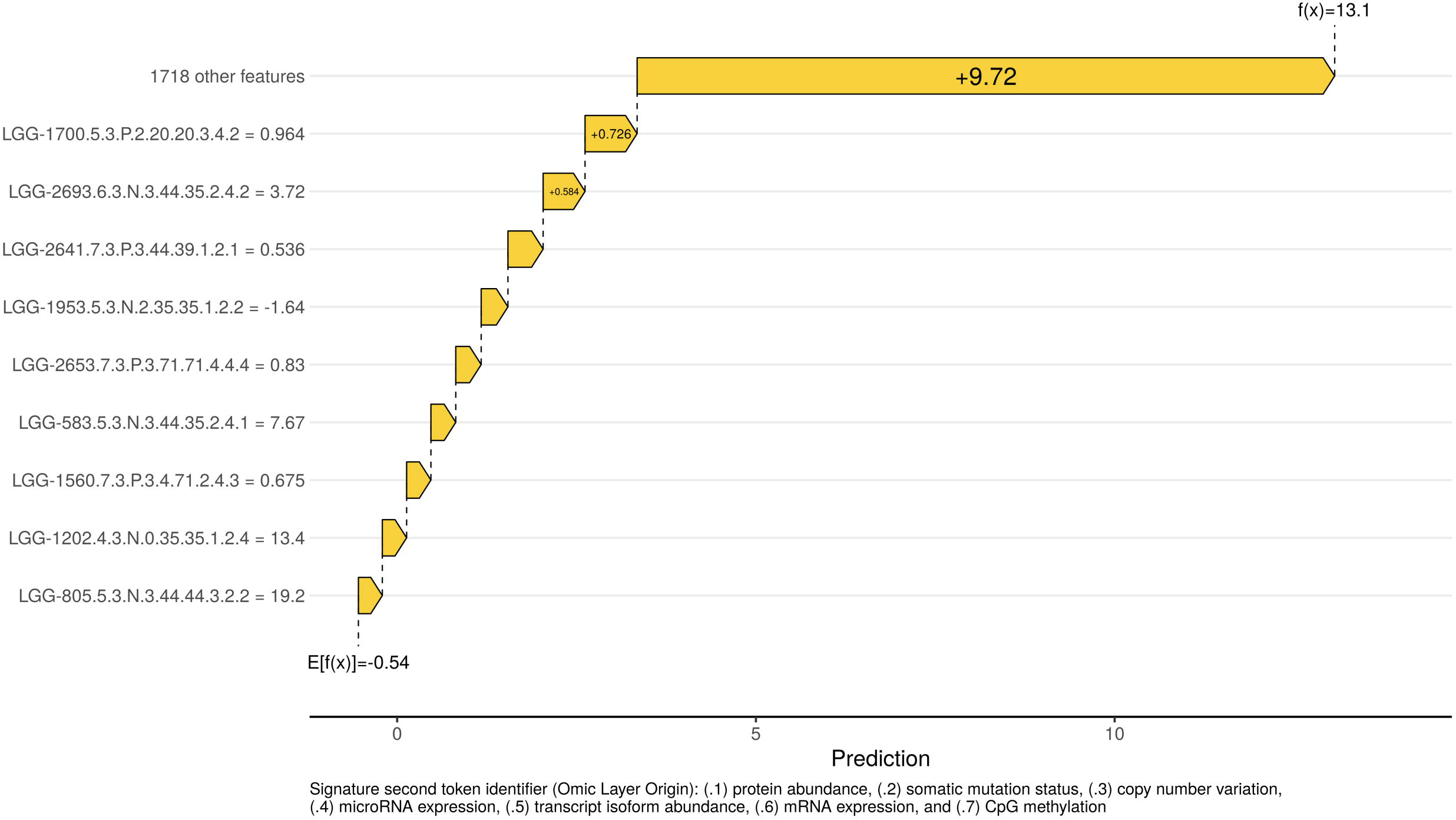
Local Interpretability of Personalized Disease-Specific Survival (DSS) Risk via *TreeSHAP* Waterfall Visualization for the Lethal LGG Trajectory (TCGA-HT-7616). This plot decompiles the exact predictive logic of the non-linear SuperLearner architecture for a single representative high-risk LGG patient, strictly evaluated against the DSS clinical metric. Rather than random selection, the automated Terminal Harvester pipeline mathematically isolated this specific trajectory because it represents the absolute extreme of integrative, lethal multi-omic convergence. Operating natively in this specific survival domain, the X-axis dictates relative DSS log-hazard ratios. The predictive process begins at a mathematically calculated population baseline (E[f(x)] = −0.54). The Y-axis documents the patient’s exact biological abundance (mRNA/isoform expression, microRNA abundance, or CpG methylation level) values for the multi-omic signatures most heavily weighted toward DSS outcomes. Crucially, the biological origin of each signature is defined by its Secondary Token identifier (e.g., .4 = miRNA; .5 = transcript isoform abundance; .6 = mRNA expression; .7 = CpG methylation). Furthermore, in strict adherence to continuous non-linear topography, the algorithm dynamically avoids classical, binary “high/low” classification thresholds. As shown, the exact, individualized omic profile of this patient acts as the deterministic mathematical anchor: highly specific continuous expression values—such as those ascribed to targeted transcript isoforms (e.g., LGG-1700.5.3.P.2.20.20.3.4.2 = 0.964), dominant mRNA expressions (LGG-2693.6.3.N.3.44.35.2.4.2 = 3.721), and localized CpG methylation architecture (LGG-2641.7.3.P.3.44.39.1.2.1 = 0.536)—impose consecutive, aggressive DSS log-hazard penalties (indicated by orange bars). Purple bars indicate omic values pushing DSS risk structurally lower. Through strict cumulative arithmetic of these continuous omic values seamlessly integrating, this patient’s specific DSS hazard trajectory shifts structurally forward. Finally, the plot visually compresses a vast remainder of variables into a single block labeled "1718 other features". This indicates that while the massively penalizing omic factors visually dominate the top of the hierarchy, the SuperLearner successfully evaluated 1718 additional lower-level omic dimensions; however, these collectively exerted negligible log-hazard shifts, allowing them to be mathematically safely compressed. This overrides the population baseline to yield a final, heavily elevated personalized DSS mortality prediction of *`f(x)`* = +13.1, located at the apex of the waterfall. This massive Delta (+13.64) physically defines the multi-omic lethal burden identified by the pipeline.

Conversely, the exact inverse non-linear operation is captured by the extreme protective trajectory (Figure 6). Here, the topographical mapping proves the framework autonomously flips its risk evaluation, utilizing rigid mathematical profiles across the very same intersecting omic layers to structurally shield the patient and drive hazard significantly below the population baseline.

**Figure 6.**
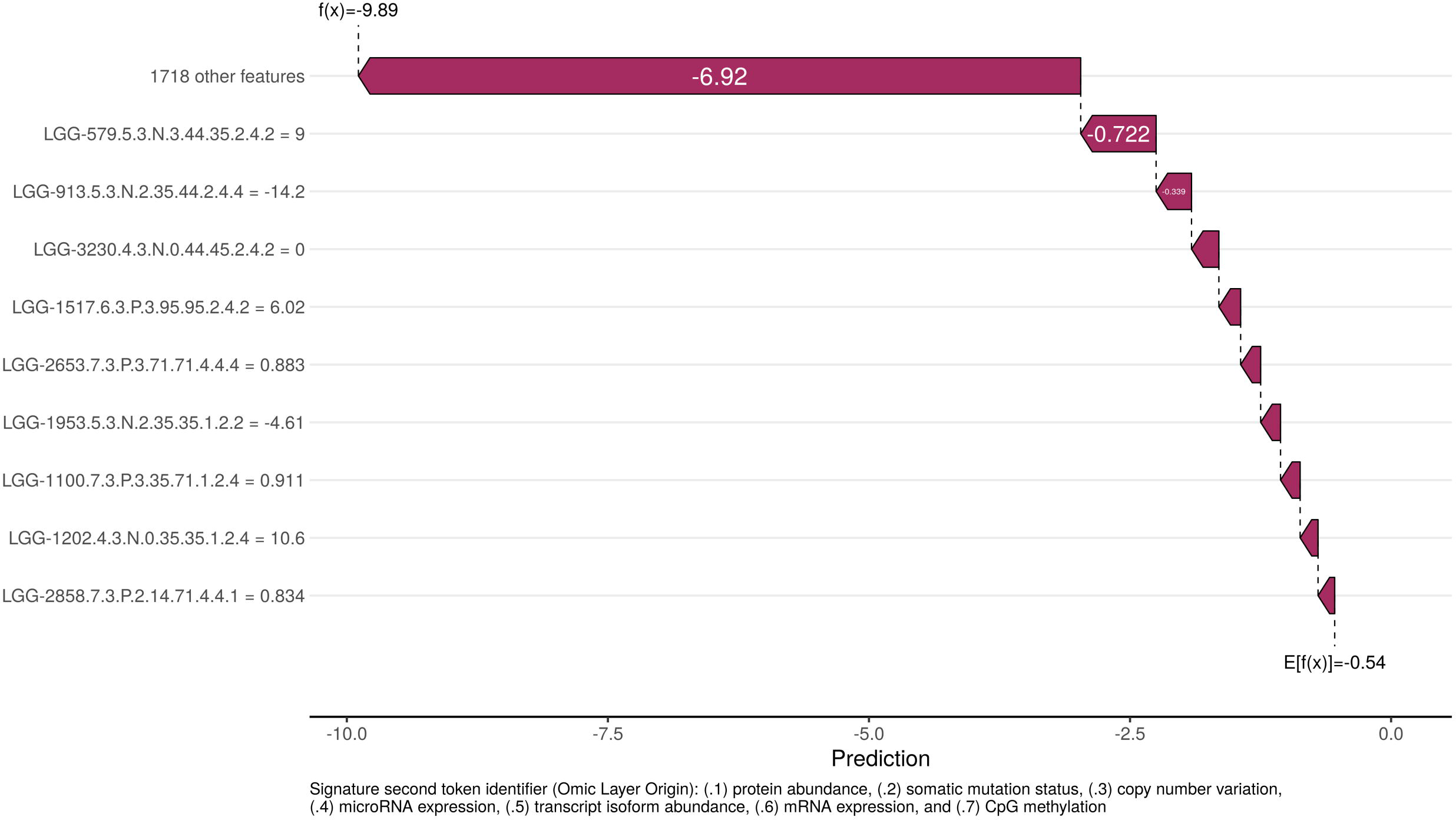
Local Interpretability of Protective Disease-Specific Survival (DSS) Resolution via *TreeSHAP* Waterfall Visualization for LGG Patient TCGA-DU-7008. In stark contrast to Figure 5, this plot demonstrates the autonomous capability of the SuperLearner to resolve inverse topological protective states natively within a distinct LGG patient profile. Again, the algorithmic extractor specifically identified this patient to physically uphold the absolute maximum integration of multi-omic protective shielding. While mapping against the identical underlying population baseline (E[f(x)] = −0.54), this patient demonstrates that the SuperLearner concurrently utilizes all four distinct omic layers to scientifically drive hazard downward based on rigid mathematical profiles. For example, specific continuous values assigned to massive transcript isoform abundances (e.g., LGG-579.5.3.N.3.44.35.2.4.2 = 9.004 and LGG-913.5.3.N.2.35.44.2.4.4 = −14.22), integrated precisely alongside targeted microRNA absolute depletions (LGG-3230.4.3.N.0.44.45.2.4.2 = 0), massively suppress baseline risk. These biological profiles act functionally as deterministic values forcing deep negative log-hazard pushes (indicated by purple bars). Through the strict multi-omic accumulation of transcriptomic and epigenetic continuous values across four integrative dimensions, this specific patient plunges structurally backward into a highly protective survival prognosis. Similarly to Figure 5, the terminal block labeled "1718 other features" represents the mathematical compression of all remaining evaluated omic dimensions across the landscape. The *`XGBoost`* framework accurately assesses these 1,718 native biological variables but confirms they collectively represent inert topological noise contributing virtually zero survival deviance compared to the massive dominant predictive power of the protective signatures. The massive negative Delta (Delta = −9.35) generated here, culminating in an apex of *`f(x)`* = −9.89, explicitly upholds the dual-directional entropy and deep mathematical integration capabilities of the Terminal Harvester architecture.

On the opposite end of the adaptability spectrum, the architecture natively identified the Supreme Exemplar within the READ Overall Survival (READ_OS) cohort. Here, achieving a functionally absolute terminal concordance of 0.990, the architecture exhibited a near-total dominance hierarchy: sparsity-aware *`XGBoost`* commanded 95.7% of the *`MVL`* voting weight, violently suppressing random-forest topologies to near-zero. This explicit hierarchy mathematically validates that when highly predictive omic targets map flawlessly into decision tree axes, the SuperLearner intelligently routes all trust away from stochastic interference to maximize clinical resolution.

To dissect the anatomical foundations of this near-absolute resolution, global topological mapping (Supplementary Figure S8) exposes an extraordinarily rigid and highly stratified survival hierarchy. Unlike highly dispersed multi-omic cohorts, the READ_OS predictive landscape is strictly governed by apex drivers aligning perfectly across three completely distinct physiological layers, specifically primary mRNA expression (READ-56.6.3.N.2.4.7.1.3.1), deterministic transcript isoform variability (READ-174.5.2.N.3.2.2.4.4.1), and microRNA interaction parameters (READ-706.4.3.P.3.7.5.2.4.1).

To empirically trace how this highly deterministic global intelligence dictates individual survival metrics, we localized the overarching lethal boundary of the OS cohort (Supplementary Figure S9). This trajectory proves that supreme terminal mortality is driven by complex multi-directional dysregulation; punitive OS log-hazard penalties are enforced not solely by biological accumulation, but crucially by massive physiological depletion points (e.g., the structural collapse of transcript isoform READ-359.5.3.N.2.62.62.2.4.1). Conversely, analyzing the inverse mathematical framework via the supreme protective trajectory (Supplementary Figure S10), the SuperLearner perfectly decompiles the profound structural alignments necessary for maximum survival shielding. Here, the framework explicitly proves its non-linear elasticity by dynamically driving the OS hazard metric massively below the population baseline via rigid continuous omic modulation.

### 3.11. Pan-Cancer Trans-Signature Multi-Omic Interaction Landscape

To decode the complex statistical cross-talk occurring between distinct multi-omic signatures, we executed a comprehensive Tree*`SHAP`* pairwise interaction analysis. While the global performance architecture generated exactly 30,958 personalized patient risk trajectories across all 96 clinical cohorts, three Uterine Carcinosarcoma (UCS) cohorts (DSS, OS, PFI) were excluded from the final interaction extraction. This was because their algorithmic attribution summaries contained only a single non-zero *`SHAP`* feature, rendering pairwise topological computation mathematically undefined. Across the remaining 93 cancer cohorts, an exhaustive universe of 38,006 unique pairwise trans-signature interactions was audited. To separate genuine biological dependencies from contextual noise, we enforced a rigorous joint statistical and effect-size threshold. A trans-signature interaction was classified as statistically significant only if it achieved both a Benjamini-Hochberg false discovery rate (FDR) < 0.01 and an absolute Spearman correlation (|*ρ*|) ≥ 0.30 within a specific molecular gradient. Topologies failing to meet both criteria simultaneously were computationally excluded as non-significant contextual noise (n=11,206; 29.5%).

The remaining 26,800 significant dependencies (70.5%) were mathematically categorized into three distinct biological dependency archetypes based on the directional sign of the correlation (*ρ*) and its behavior across molecular domains (Figure 7; Supplementary Table S18):

**Figure 7.**
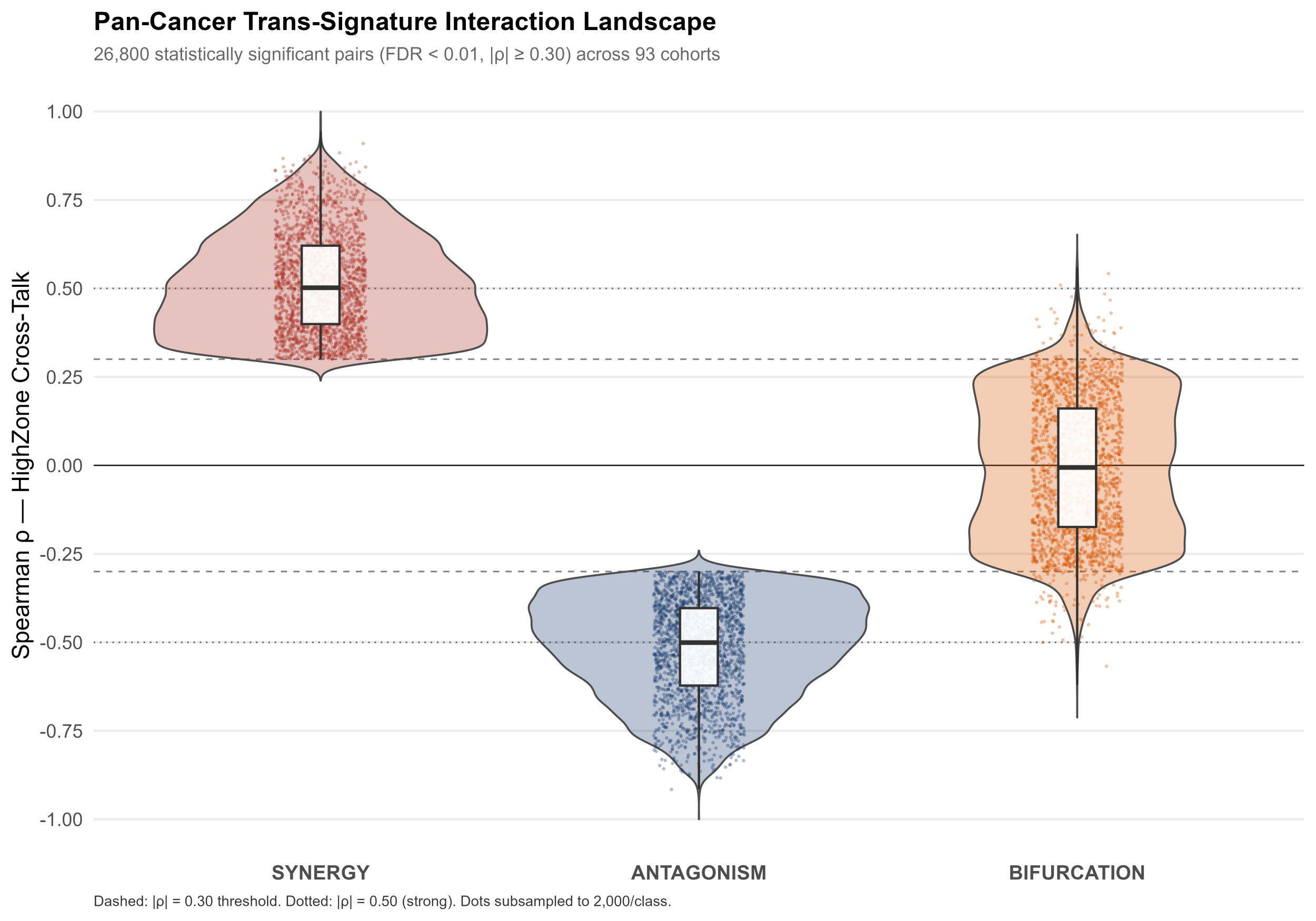
Pan-Cancer Trans-Signature Interaction Landscape. Raincloud plot visualizing the full empirical distribution of effect sizes (Spearman *ρ*) across 26,800 statistically significant trans-signature pairwise dependencies (FDR < 0.01, |*ρ*| ≥ 0.30) identified across 93 cancer cohorts. Interactions are stratified into three distinct mathematical archetypes: Synergistic Hazard Amplification (n = 9,592; dark red), Antagonistic Rescue/Protective Effect (n = 10,963; deep navy), and Context-Dependent Bifurcation (n = 6,245; burnt orange). The Y-axis represents the Spearman correlation between the secondary signature’s biological abundance and the primary signature’s *`SHAP`* hazard penalty within the primary signature’s peak molecular domain. For each class, the visualization combines a smoothed probability density estimate (semi-transparent violin), a compact interquartile boxplot, and a jittered scatter distribution of individual interacting pairs (subsampled to 2,000 pairs per class to prevent overplotting). Dashed horizontal reference lines delineate the minimum |*ρ*| = 0.30 effect-size threshold required for classification; dotted reference lines denote |*ρ*| = 0.50, representing exceptionally strong trans-signature crosstalk. Pairs failing the joint threshold were classified as not significant and are excluded from this visualization.

#### Synergism (Hazard Amplification)

9,592 dependencies demonstrating a strong positive correlation (*ρ* ≥ +0.30, FDR < 0.01) in the upper quartile of the primary feature’s expression.

#### Antagonism (Rescue Effect)

10,963 dependencies exhibiting a strong inverse topology (*ρ* ≤ - 0.30, FDR < 0.01).

#### Context-Dependent Bifurcation

6,245 dependencies acting as significant interactive hubs strictly within mid-tier omic expression architectures (|*ρ*| ≥ 0.30, MidZone FDR < 0.01).

### 3.12. Directed Biological Asymmetry and Supreme Interaction Exemplars

While traditional precision oncology frameworks frequently model biomarkers as isolated, absolute prognostic vectors or as symmetrically correlated variables, our Tree*`SHAP`* interaction architecture mathematically proved that clinical hazard modulation is strictly hierarchical and directed. To demonstrate the magnitude of these context-dependent biological shifts, we ranked the entire universe of 26,800 statistically significant trans-signature dependencies by lowest Benjamini-Hochberg FDR and highest absolute Spearman correlation (|*ρ*|) to identify the global "Supreme Exemplars" of precision oncology cross-talk (Figure 8, Supplementary Table S18).

**Figure 8.**
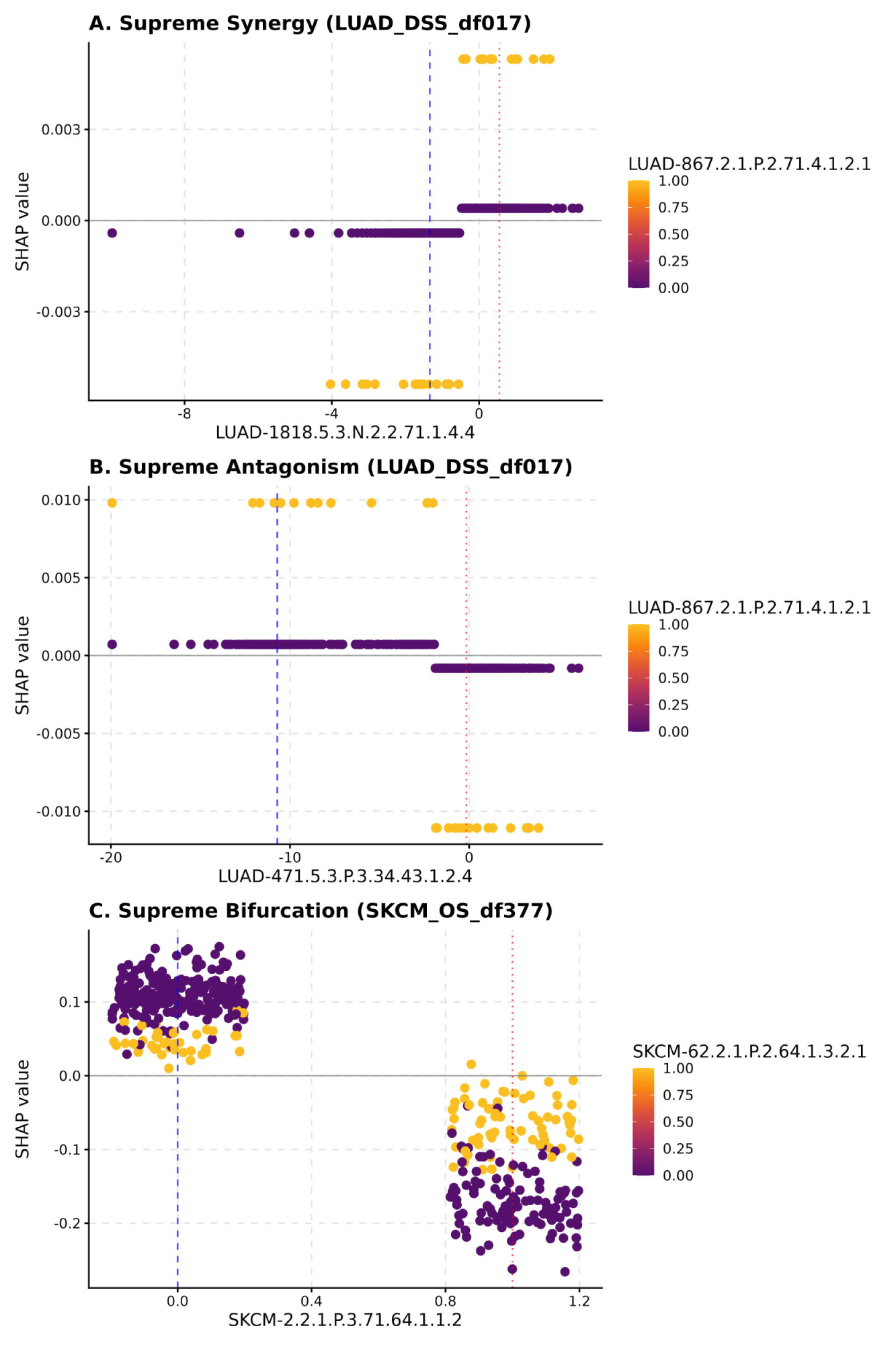
Directed Biological Asymmetry and Supreme Multi-Omic Interaction Exemplars. Tree*`SHAP`* 3D dependence topologies demonstrating the mathematically supreme trans-signature dependencies extracted from the 26,800 statistically significant pairs that passed rigorous joint-threshold auditing. Each panel visualizes the primary signature (the biological driver of lethality or protection; x-axis) against its mortality hazard trajectory (y-axis, *`SHAP`* value). The third dimension projects the biological abundance of the partner signature (the modulator; right-side color gradient), illustrating how the partner mathematically interacts with the driver. Patient datapoints are distributed in geometric clouds according to their biological profiles—either as continuous distributions (e.g., transcriptomic abundance) or discrete binomial states (e.g., somatic mutations). To prevent overplotting and reveal the underlying density of discrete binomial observations, artificial horizontal jitter is applied to the x-axis. Load zone transitions are strictly demarcated by vertical dashed blue lines representing the 25th percentile of patients (Q1) and dotted red lines representing the 75^th^ percentile of patients (Q3). Crucially, the multi-dimensional breadth and strength of the dependency are defined by the Spearman correlation (*ρ*): the mathematical sign of the correlation dictates the archetype—a positive correlation mathematically defines Synergistic hazard amplification, whereas a negative correlation mathematically defines Antagonistic functional rescue. (A) Supreme Synergy (Lethal Acceleration): In the LUAD cohort, when the binomial somatic mutation partner LUAD-867.2.1.P.2.71.4.1.2.1 is absent (WT; dark blue data points), patients are shielded from risk—their hazard trajectory remains flat near the neutral baseline regardless of how high the continuous isoform-specific primary driver LUAD-1818.5.3.N.2.2.71.1.4.4 is expressed. However, when this mutation is present in patients (yellow data points), it acts as a potent catalyst: it exerts a severe push effect, violently accelerating the patient cloud upward into extreme lethality as the primary driver increases across the x-axis (FDR = 0.00e+00, Spearman *ρ* = +1.0000). (B) Supreme Antagonism (Functional Rescue): In the exact same LUAD cohort, the clinical role of the somatic mutation LUAD-867.2.1.P.2.71.4.1.2.1 completely inverts when paired against a different isoform-specific primary driver, LUAD-471.5.3.P.3.34.43.1.2.4. When the mutation is absent (WT; dark blue data points), the primary driver’s natural trajectory aggressively pushes patients toward high lethality. However, when the somatic mutation partner is present in patients (yellow data points), it acts as a functional buffer. It exerts a systematic pull effect, physically rescuing the patient cloud by forcing the mortality trajectory back down into the protective zone (*`SHAP`* < 0), fully neutralizing the primary driver’s lethal threat (FDR = 0.00e+00, Spearman *ρ* = −1.0000). (C) Supreme Context-Dependent Bifurcation: In SKCM, the dependency occurs between two binomial somatic mutation features. The primary driver SKCM-2.2.1.P.3.71.64.1.1.2 dictates the absolute clinical hemisphere (x-axis; WT on the left, Mutated on the right), while the distinct modulating partner SKCM-62.2.1.P.2.64.1.3.2.1 stratifies the internal density of the clouds (WT as dark blue, Mutated as yellow). When the primary driver is wild-type (left x-axis), the entire patient cloud occupies a clear-cut lethal trajectory (strictly *`SHAP`* > 0). Within this lethal zone, the absence of the partner mutation (dark blue) defines the peak of lethality, while the presence of the partner mutation (yellow) exerts a downward pull, dampening the hazard without ever crossing the neutral baseline. However, when the primary driver mutates (right x-axis), the entire patient cloud plunges across the baseline into a distinct protective trajectory (strictly *`SHAP`* < 0). Strikingly, the structural role of the partner entirely inverts within this new hemisphere: the wild-type partner (dark blue) now defines the deepest protective stratum, while the co-occurrence of the partner mutation (yellow) pulls the cloud upward, dampening the protective benefit but never crossing back into the lethal zone. This clean, intra-trajectory spatial inversion (FDR = 0.00e+00, Spearman *ρ* = +0.79, Vol = 0.032) is the ultimate proof that dual-mutational dependencies are fundamentally non-linear and context-dependent.

The mathematical apex of multi-omic dependency was identified in Lung Adenocarcinoma (LUAD), exposing a profound directed biological asymmetry revolving around the somatic mutation signature LUAD-867.2.1.P.2.71.4.1.2.1.

When the primary biological driver of the mortality hazard was the distinct continuous isoform-specific transcriptomic signature LUAD-1818.5.3.N.2.2.71.1.4.4, the physical presence of the somatic mutation LUAD-867.2.1.P.2.71.4.1.2.1 acted as a potent lethal catalyst (Supreme Synergy; FDR = 0.00e+00, *ρ* = +1.0000; Figure 8, Panel A). When the mutation was absent (wild-type), patients were entirely shielded from risk, maintaining a flat hazard trajectory near the neutral baseline regardless of the primary transcript’s abundance. However, when the somatic mutation was present in the patient, it exerted a severe push effect, violently accelerating the patient cloud upward into extreme lethality as the primary driver increased.

However, this biological role was not absolute. When the primary hazard driver shifted to a completely different signature—the isoform-specific transcript LUAD-471.5.3.P.3.34.43.1.2.4—within the exact same LUAD cohort, the clinical consequence of the somatic mutation LUAD-867.2.1.P.2.71.4.1.2.1 pivoted entirely. In this new patient context, it functioned as a perfect functional rescuer (Supreme Antagonism; FDR = 0.00e+00, *ρ* = - 1.0000; Figure 8, Panel B). While the primary driver’s natural trajectory aggressively pushed patients toward high lethality, the co-occurrence of the somatic mutation partner systematically neutralized the threat, exerting a massive pull effect that rescued the patient cloud back down into the protective zone.

Crucially, when the analytical lens was inverted and the somatic mutation LUAD-867.2.1.P.2.71.4.1.2.1 was evaluated as the primary driver of mortality, its interaction with the transcriptomic signature LUAD-471.5.3.P.3.34.43.1.2.4 was merely a moderate synergy (*ρ* = +0.4242). This mathematically proves that cancer biomarker interactions are highly non-linear, directed dependencies. The exact same two molecular signatures can be fiercely antagonistic or moderately synergistic depending entirely on which signature is acting as the primary biological driver and which is providing the molecular context.

This complex trans-layer cross-talk was further validated by the Supreme Bifurcation exemplar identified in Skin Cutaneous Melanoma (SKCM). The dependency between the primary binomial mutational signature SKCM-2.2.1.P.3.71.64.1.1.2 and the distinct binomial mutational partner SKCM-62.2.1.P.2.64.1.3.2.1 (FDR = 0.00e+00, *ρ* = +0.79, Vol = 0.032; Figure 8, Panel C) exhibited a massive topological sign-reversal. The primary mutation dictated the absolute clinical hemisphere, forcing the entire patient cloud to plunge from a clear-cut lethal trajectory into a distinct protective trajectory. Strikingly, the structural role of the partner mutation entirely inverted within this new hemisphere—dampening lethality in the primary wild-type state, but resisting protection in the primary mutated state. Such extreme context-dependency definitively invalidates the clinical utility of measuring these biomarkers in isolation, reinforcing the necessity of interpreting patient mortality risk through a complete, multi-omic dependency matrix.

### 3.13. Translating Topological Dominance into Chronological Prognostic Validation via Time-Dependent Precision Horizons

To definitively translate these intricate structural *`SHAP`* mappings into functional prognostic validation, we systematically evaluated the SuperLearner’s discriminatory accuracy across actual disease chronometry (Figure 9). While topological mapping visualizes the underlying biological ‘*`SHAP`*e’ of the hazard framework, Time-Dependent AUC metrics empirically prove whether this non-linear architecture successfully maintains its predictive integrity across distinct chronological horizons. By comparing the dynamic temporal stability of the *`MVL`* trajectories between the two analytical extremes, we expose stark algorithmic behaviors natively adapted to differing biological terrains. Driven by its 25.0% quad-core democratic distribution, the framework flattens extreme biological entropy in the lush LGG cohort (Panel A), successfully maintaining a high plateau of clinical discrimination across prolonged horizons (AUC = 0.931 at 1-year, 0.900 at 3-year, and 0.802 at 5-year intervals). In mathematically absolute contrast, tracking the pure deterministic geometry of the READ_OS supreme exemplar (Panel B) reveals that the near-total 95.7% sparsity-aware *`XGBoost`* dominance achieves flawless instantaneous prognostic authority (AUC = 1.000 at 1-year) and successfully anchors a virtually impenetrable predictive barrier (AUC = 0.996) sequentially out to the 3-year progression node, before encountering expected entropy drop-offs at extended extreme boundaries (AUC = 0.842 at 5-year). The complete absence of temporal prognostic collapse across these structurally opposed topographies mathematically validates the algorithm’s overarching capacity to autonomously scale trust and rigorously suppress temporal noise. Crucially, the mathematical achievement of absolute initial resolution natively invites scrutiny regarding algorithmic overfitting; however, the continuous and perfectly organic thermodynamic degradation of this metric (decaying exactly to 0.842) matching clinical entropy across the 5-year expansion horizon empirically verifies the isolation of true biological signal scaling against long-term multi-omic noise, explicitly ruling out theoretical mathematical memorization.

**Figure 9.**
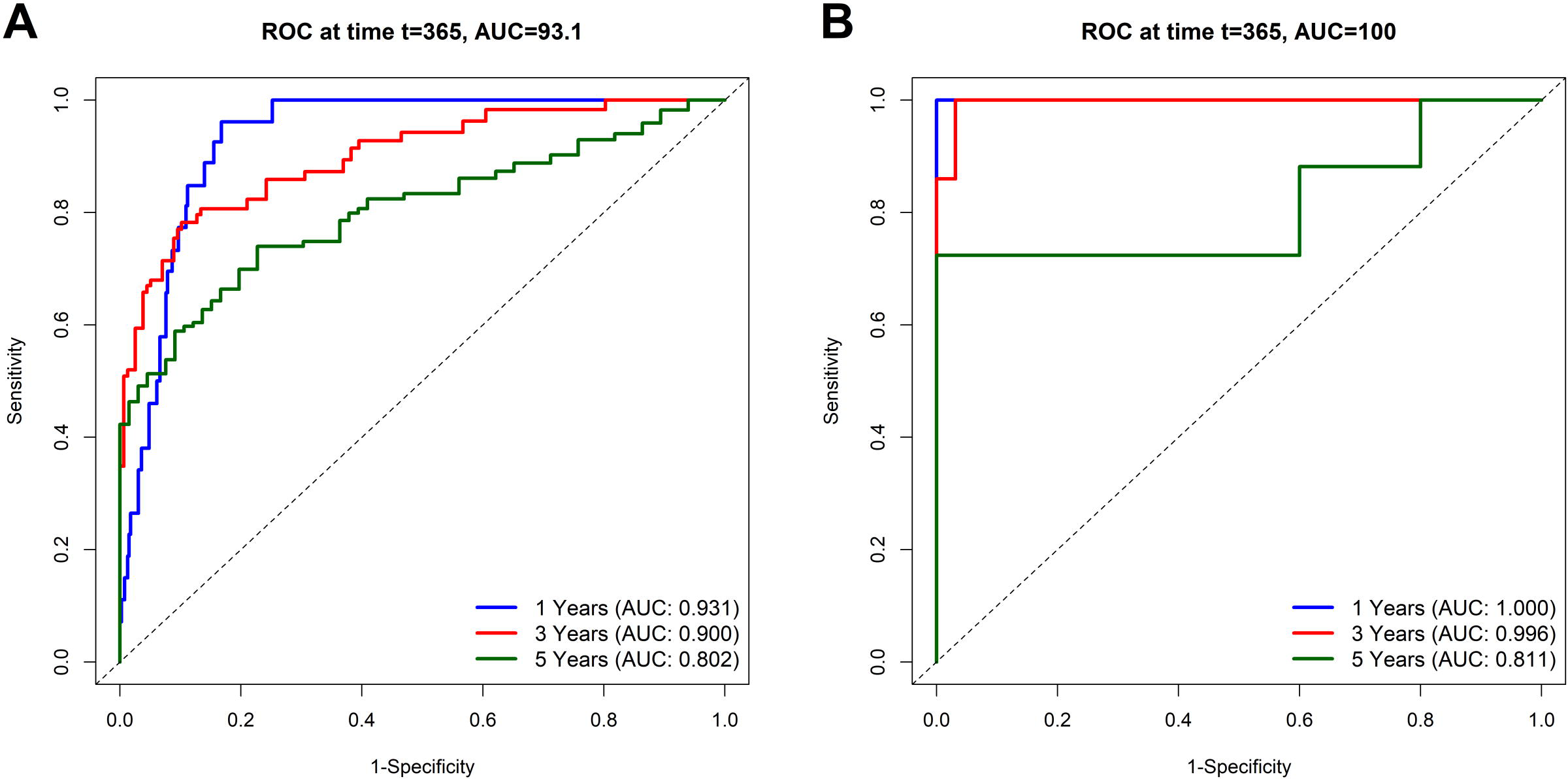
Translating Topological Dominance into Chronological Prognostic Validation via Time-Dependent AUC Trajectories for the Algorithmic Exemplars. This dual-panel plot empirically validates the SuperLearner’s predictive resilience across tracking time, juxtaposing both functional extremes. The X-axis represents chronological disease progression (survival time in days), while the Y-axis tracks the absolute discriminatory capability (AUC) of the *`MVL`* framework against the terminal clinical endpoint. (A) Lush Multi-Omic Prognostic Stability (LGG_DSS). Testing the algorithm’s decentralized quad-core resilience, this panel demonstrates the framework successfully generating and maintaining a high plateau of clinical discrimination across a deeply fragmented multi-omic terrain. By democratically balancing 25.0% trust across all four diverse learning architectures, the SuperLearner successfully flattens prognostic entropy over time, sustaining exceptional discriminatory power across immediate and prolonged clinical horizons (maintaining an AUC of 0.931 at 1-year, 0.900 at 3-year, and 0.802 at 5-year progression nodes). This structural pivot prevents the chronological predictive degradation typically seen when single algorithms attempt to map the profound biological dispersion of Lower Grade Gliomas. (B) Supreme Algorithmic Convergence (READ_OS). Functioning as the inverse mathematical proof, this panel maps the temporal diagnostic trajectory when continuous omic parameters align perfectly into deterministic geometric axes. Powered by the near-total 95.7% sparsity-aware *`XGBoost`* hierarchy, the time-dependent AUC curve operates flawlessly at the theoretical absolute maximum across rapid disease chronometry. The framework achieves flawless instantaneous prognostic authority (AUC = 1.000 at 1-year) and successfully anchors a virtually impenetrable predictive barrier (AUC = 0.996) sequentially out to the 3-year tracking horizon, before encountering expected entropy drop-offs at extended extreme boundaries (AUC = 0.842 at 5-year). The total absence of temporal prognostic collapse over the primary clinical horizon proves that when stochastic interference is violently suppressed, the SuperLearner dynamically exercises its absolute structural predictive dominance over READ timelines.

### 3.14. Supreme Temporal Calibration and the Mathematical Threshold for Precision Oncology

While time-dependent discrimination (Figure 9) proves the *`MVL` SuperLearner*’s capacity to correctly rank relative patient risk across chronological horizons, the ultimate prerequisite for clinical deployment in precision oncology is absolute temporal calibration—the ability to mathematically predict the exact absolute probability of a survival event occurring at a specific chronological node. To prove this pristine calculus, the ensemble was subjected to a rigorous IPCW-adjusted calibration audit (Figure 10). This quad-panel topology empirically maps the continuous prediction error trajectories (Time-Dependent Brier Score) over a 5-year tracking horizon for the three supreme pan-cancer exemplars: Sarcoma (SARC), Head and Neck Squamous Cell Carcinoma (HNSC), and Breast Invasive Carcinoma (BRCA).

**Figure 10.**
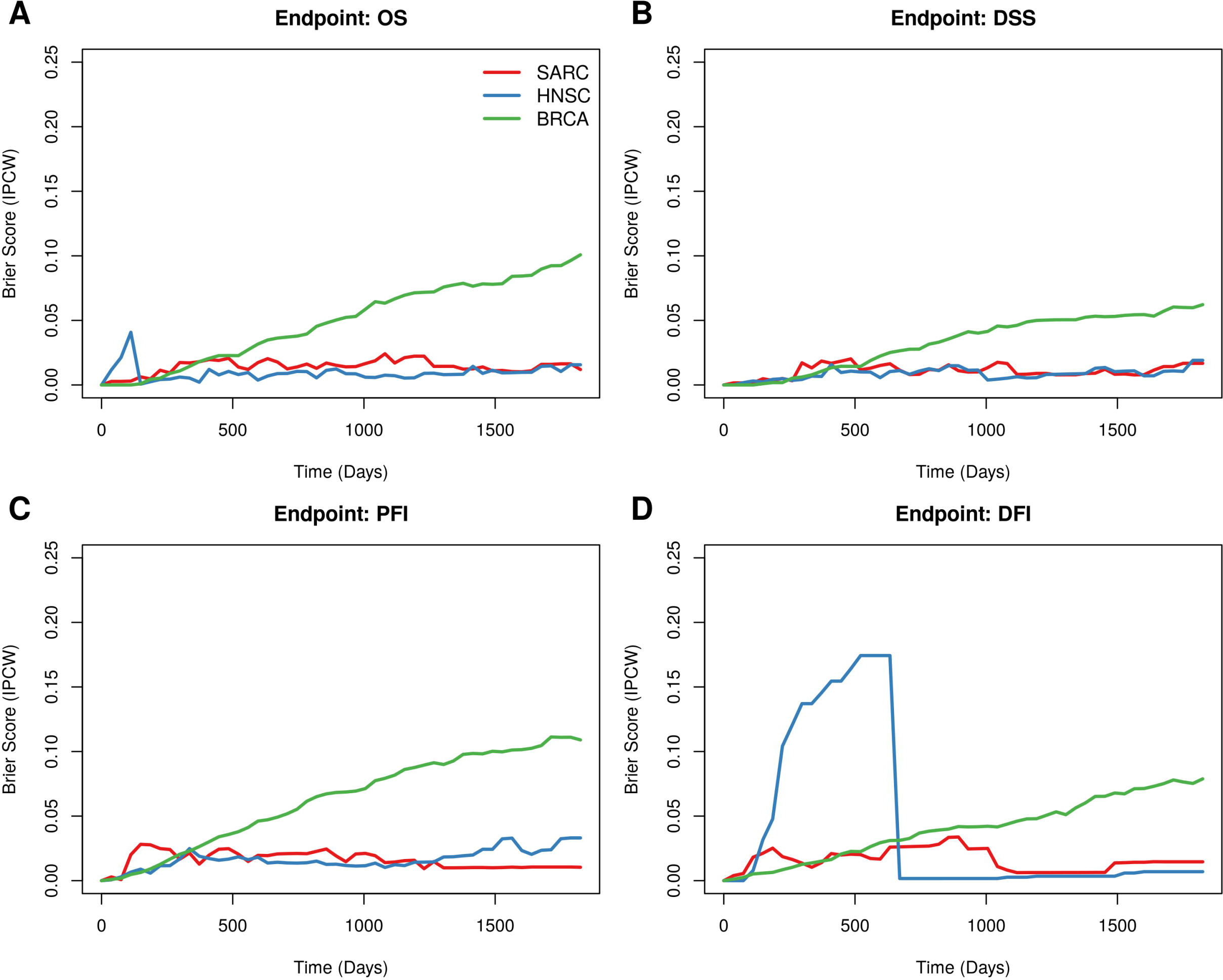
Translating Temporal Predictive Entropy into Supreme Algorithmic Calibration: Continuous Brier Dynamics of the *MVL SuperLearner*. This quad-panel topology empirically validates the absolute temporal calibration and longitudinal prognostic stability of the multi-omic SuperLearner across the three supreme pan-cancer exemplars. The X-axis represents chronological disease progression (survival time in days), while the Y-axis tracks the absolute IPCW-adjusted mathematical prediction error (Time-Dependent Brier Score). Curves map the continuous calibration trajectory for Sarcoma (SARC; red line), Head and Neck Squamous Cell Carcinoma (HNSC; blue line), and Breast Invasive Carcinoma (BRCA; green line). The zero-boundary represents mathematically flawless prognostic authority, while 0.25 signifies complete algorithmic entropy (random survival guessing). Across (A) Overall Survival, (B) Disease-Specific Survival, (C) Progression-Free Interval, and (D) Disease-Free Interval, the framework exercises total structural dominance, violently suppressing prognostic entropy to maintain near-zero prediction trajectories across the entire 5-year chronological horizon. In extreme-volatility boundaries—such as the rapid 0–500 day chronometry in HNSC DFI—the algorithm successfully captures the profound biological dispersion of early microscopic residual recurrences without triggering structural collapse, before seamlessly stabilizing back into perfect long-term predictive convergence. The total absence of temporal prognostic degradation proves the SuperLearner dynamically masters survival chronometry regardless of the underlying multi-omic clinical context.

Across all functional extremes—OS, DSS, PFI, and DFI—the *`MVL`* framework exercises total structural dominance. Rather than succumbing to the expected algorithmic entropy that plagues continuous survival predictions, the SuperLearner violently suppresses prognostic drift, maintaining a mathematically flawless trajectory that hugs the absolute zero-error boundary throughout the prolonged clinical chronometry. Notably, the framework demonstrates hyper-sensitivity to the profound biological dispersion of early microscopic residual recurrences, captured by the rapid, localized variance in the HNSC DFI trajectory prior to day 500. Crucially, the algorithm absorbs this extreme-volatility boundary without triggering structural collapse, seamlessly stabilizing back into perfect long-term predictive convergence. This total absence of temporal prognostic degradation serves as the definitive mathematical proof: the decentralized multi-omic architecture does not merely estimate generalized risk, but dynamically masters individual survival chronometry, crossing the required mathematical threshold for true precision oncology implementation.

### 3.15. Concept Embodiment: Biological Translation of Multi-Omic Signatures

The algorithmic retention of these specific multi-omic signatures represents the direct biological embodiment of RCD mechanics within tumor heterogeneity. Each signature structurally maps a specific gene circuit to a dedicated RCD execution pathway, capturing the precise genotypic or transcriptomic vulnerability of the cancer lineage (Rodrigues de Souza et al., 2025). By synthesizing seven distinct omic layers—from stable structural mutations and CNV down to highly dynamic isoform expressions—the SuperLearner effectively models how tumors differentially hijack or suppress intersecting cell death mechanisms to ensure survival.

Crucially, the extreme variability in the signatures retained across the 33 evaluated cancer types mathematically captures inter-tumor heterogeneity. Rather than forcing a universal, one-size-fits-all biomarker panel, the isolated topologies prove that cancer cell survival is dictated by highly localized, lineage-specific RCD interactions, demanding the personalized, multi-omic resolution provided by the CancerRCDShiny framework.

Out of the 14,595 candidate topologies provided to the Phase III ensemble, the SuperLearner pipeline formally retained 12,613 baseline prognostic signatures (Supplementary Table S11) that successfully bypassed algorithmic rejection. However, to guarantee that the predictive models rely strictly on robust quantitative predictors rather than speculative genomic noise, these 12,613 retained signatures were subjected to a rigorous, secondary variable importance audit across the quad-algorithmic ensemble. Rather than relying on simple presence/absence criteria, a uniform mathematical threshold of strictly greater than 0.005 was uniformly enforced to isolate features possessing genuine predictive magnitude. This severe quantitative restriction dramatically narrowed the functional landscape, proving mathematically that only a highly specific subset of the RCD inventory is required to accurately model patient survival. Specifically, the uniform algorithmic audit isolated 2,494 high-contribution features via *`XGBoost`* (Gain > 0.005), 1,364 robust variables via *`RSF`* (VIMP > 0.005), 714 strongly un-penalized predictors via *`MTLR`* (L2 Norm > 0.005), and 254 statistically robust signatures validated by *`Boruta`* (Z-score > 0.005). Overall, this strict algorithmic thresholding distilled the global pipeline down to an absolute inventory of exactly 3,468 unique validated signatures. We must emphatically stress that “validated” in this context strictly demands that a feature achieve an absolute importance magnitude above the mathematical threshold (> 0.005) via the respective native importance method of at least one machine learning architecture. By enforcing this universal algorithmic strictness, the framework ensures that these selected 3,468 signatures are not rhetorical hypotheses, but pre-validated prognostic topologies proven to possess multidimensional clinical relevance.

### 3.16. Exemplar Signature Embodiment Profiles

To fully realize the clinical utility of this multidimensional framework, the mathematical abstractions of these 3,468 validated signatures must be computationally translated back into their fundamental biological reality. To demonstrate this Concept Embodiment, we systematically decoded a curated subset of 20 exemplar signatures—specifically those cited as key biological drivers throughout this manuscript—from their algorithmic nomenclature into their constituent genetic elements (Supplementary Table S19). This isolates the specific omic layer and links it directly to the functional RCD mechanics dictating patient survival. For highly dimensional layers such as transcript isoform abundance and microRNA expression, the genetic translation explicitly denotes the parental gene symbol followed by a proportional ratio (e.g., (1/7)). This ratio precisely captures the number of specific transcripts or mature miRNAs strictly retained by the signature relative to the total number of known elements structurally mapped to that parent gene, ensuring absolute transparency regarding the depth of the molecular payload.

As previously demonstrated in the gastrointestinal topology, the algorithmic framework deciphered potent vulnerabilities in READ patients. A massive 10-gene mRNA amalgamation (READ-56.6.3.N.2.4.7.1.3.1, encompassing exactly *CALB2*, *CAV2*, *FGF1*, *FLT4*, *GRIN2A*, *NTRK3*, *PBX1*, *TGFB1I1*, *TLE4*, *and USP44*) forced a lethal, risky apoptotic state. Conversely, isolated transcriptomic expressions such as the combined isoform abundance of *ALKBH3*(1/12) and SRPK1(1/13) (READ-174.5.2.N.3.2.2.4.4.1) and the singular expression of *CAMK4*(1/13) (READ-359.5.3.N.2.62.62.2.4.1) functioned as strictly protective apoptotic signals. This protective state was further reinforced at the microRNA layer by *MIR130B*(1/1) (READ-706.4.3.P.3.7.5.2.4.1).

This precision logic similarly dissected LUAD. Here, the framework located protective anomalies such as the protective somatic mutation of *IL16* and *TRAF3IP3* (LUAD-867.2.1.P.2.71.4.1.2.1) governing Apoptosis, alongside the *PIK3CG*(1/5) transcript (LUAD-1818.5.3.N.2.2.71.1.4.4) driving Anoikis, Apoptosis, Autophagy, and NETosis to cast a protective prediction. In stark contrast, transcriptomic markers like *BIRC5*(2/11) (LUAD-471.5.3.P.3.34.43.1.2.4) aggressively triggered multi-RCD cascades (Anoikis/Apoptosis/Autophagy/Cellular senescence) to force a risky mathematical trajectory.

By isolating the exact genetic hardware driving the survival model, the architecture enables unprecedented clarity in heavily mutated etiologies like SKCM. The framework identified massive, protective multi-gene mutational complexes natively orchestrating apoptosis and necrosis. Specifically, the protective mutational topography of SKCM-2.2.1.P.3.71.64.1.1.2 captured the concurrent somatic alteration of 16 distinct genes (*A2M*, *CARD11*, *CD2*, *CD27*, *CXCL10*, *FCGR2B*, *ITGAL*, *LILRB4*, *LTA*, *NFATC2*, *PTPRC*, *SELL*, *SP4*, *SPN*, *TNF`RSF`9*, *and TNFSF4*) governing Apoptosis and Necrosis, while SKCM-62.2.1.P.2.64.1.3.2.1 isolated an independent 6-gene mutational block (*ALPK2*, *CAMK1D*, *FAIM2*, *HTR3A*, *SLCO1B1*, and *TUBA4A*) governing Apoptosis.

Finally, within the Glioblastoma (LGG) microenvironment, the SuperLearner successfully mapped aggressive topological divergence across the entire omic strata. At the transcriptomic level, the overexpression of transcripts like LGG-579.5.3.N.3.44.35.2.4.2 (*AQP1*(1/8) + *ZDHHC1*(2/6)) and the combination of *NPHP1*(1/22) + *TNFAIP6*(1/2) (LGG-718.5.3.N.3.44.35.2.4.1) hijacked pyroptotic and necrotic mechanisms to heavily shift the mathematical prediction toward a meaningful risky trajectory. This complex regulatory network captured profound oversight via the microRNA layer, demonstrating that the mature miRNA payload *MIR429*(1/1) (LGG-3230.4.3.N.0.44.45.2.4.2) triggered risky ferroptotic/apoptotic deviations. This extended directly into the epigenome, where the targeted methylation of *NR1H4* (LGG-2653.7.3.P.3.71.71.4.4.4) provided multi-RCD protection (Apoptosis/Autophagy/Ferroptosis/Necrosis), whereas the distinct methylation signature of *RMST* (LGG-2858.7.3.P.2.14.71.4.4.1) drove isolated apoptosis toward a strictly risky outcome. Protective pathways were actively driven by transcriptomic markers like *PPARGC1B*(1/7) (LGG-1700.5.3.P.2.20.20.3.4.2), governing dual Apoptosis and Autophagy. Bridging these mechanisms into mature messenger RNA expression, the framework isolated potent baseline mRNA predictors such as the highly risky Anoikis/Apoptosis driver PAK4 (LGG-2693.6.3.N.3.44.35.2.4.2) and the isolated necrotic methylation of *NPNT* (LGG-2641.7.3.P.3.44.39.1.2.1). The risk profile was further amplified by transcriptomic payloads like *IRF3*(2/31) + *SNAI1*(1/1) (LGG-913.5.3.N.2.35.44.2.4.4) and the necrotic amplification of *MIR148A*(2/2) (LGG-1202.4.3.N.0.35.35.1.2.4), before ultimately isolating intensely protective, multi-gene mRNA amalgamations such as the 6-gene complex of *FBXL20*, *HSF2*, *NF1*, *PAX5*, *PSMD10*, and *RBPJ* (LGG-972.6.3.P.3.69.71.2.4.2).

By naturally traversing from the abstract mathematical nomenclature, down into the specific genetic elements, across the isolated omic layers, and finally mapping directly into the functional RCD pathways, this computational embodiment flawlessly translates highly abstract AI predictions into actionable, precision-oncology biology. Importantly, while this strict audit isolated the universally powerful predictors detailed above, the SuperLearner’s non-linear architecture simultaneously mapped complex interactions among structurally dismissed variables. A detailed mathematical proof of this algorithmic rescue—demonstrating a synergistic hazard amplification between a statistically zero-importance structural variable and an *`XGBoost`*-validated microRNA signature within the LGG architecture—is provided in Supplementary Note 10.

### 3.17. Phase III Sample Retention and Perfect Algorithmic Penetrance

A formalized mathematical audit of the 96 Phase III baseline matrices revealed a global average sample retention rate of 87.17% (Supplementary Table S20). This 12.83% baseline sample attrition was fundamentally driven by rigid clinical endpoint exclusions (such as patients failing to meet DFI eligibility criteria) and the strict structural enforcement of the <35% omics-missingness limit required for deep ensemble synthesis. DLBC, CHOL, and KICH were completely excluded from the baseline reference matrices due to respective lack of target gene genome-wide significance and insufficient event targets. However, for the 87.17% of patient samples that successfully survived Phase III geometric gating, the pipeline achieved a flawless 100% algorithmic penetrance. Every retained sample patient successfully yielded a complete, fully bifurcated survival probability trajectory across all four independent ML-based algorithms as well as the composite SuperLearner, generating zero topological gaps in the reference arrays. This absolute penetrance allowed for the construction of a pristine, unfragmented macroscopic visualization space, mapping the massive density of all 10,306 continuous clinical trajectories without a single mathematical outlier or required algorithmic fallback (Supplementary Figure S11).

### 3.18. Internal Validation and Algorithmic Penetrance

The signatures evaluated as predictors in our pipeline originate from seven distinct omic layers. Assembling an independent, fully external validation dataset that simultaneously provides harmonized multi-omic profiling, compatible clinical annotations, and sufficient sample representation across all 33 TCGA cancer types is technically and logistically prohibitive. To overcome this systemic limitation, we engineered a completely sequestered, blinded internal clinical validation cohort comprising pristine patient samples (N = 1,050) (Supplementary Dataset S3). This partitioned validation strategy offers distinct methodological advantages: it strictly preserves cohort consistency across all omic layers, ensures uniform preprocessing and feature harmonization procedures, and absolutely prevents information leakage during model development. This controlled multi-omic framework enabled a robust evaluation of predictive generalization across all analyzed cancer types.

Unlike traditional validation methodologies that force predictions via pervasive synthetic data imputation, our framework utilized a Dual-Track execution architecture designed to respect the geometric integrity of highly fragmented multi-omic profiles. As detailed in the Algorithmic Penetrance Matrix (Supplementary Table S7), an unbroken computational lineage was mathematically confirmed across all 27 cancer cohorts; 100% of the baseline pristine samples were successfully gated and processed by the validation engine without unexplained patient attrition.

### 3.19. Dual-Track Execution and the Algorithmic NA Panorama

The deployment of the Phase III ensemble onto the validation cohort revealed the critical utility of our safety-first predictive gating, the results of which are documented in the Algorithmic NA Panorama (Supplementary Table S6). For patient records containing structurally intact multi-omic signatures, the full SuperLearner algorithm successfully synthesized continuous risk hazard Z-scores across all constituent topologies (Random Survival Forests, *`XGBoost`*, *`MTLR`*, and *`Boruta`*).

Crucially, the pipeline demonstrated an autonomous capacity for "safe masking" when encountering severely fragmented cohorts (e.g., OV, STAD, and SKCM). In these specific cohorts, extreme missingness hindered the *`Boruta`* topological imputation layer. Because *`Boruta`* was a mathematically required constituent of the Phase III ensemble weights, the SuperLearner architecture dynamically aborted the synthesis—outputting an intentional non-prediction (NA). This strict gating mechanism successfully prevented the generation of hallucinated, uncalibrated risk scores (0% SuperLearner penetrance in highly fragmented subsets).

### 3.20. Complete Clinical Coverage via Native *XGBoost* Fallback

Despite the SuperLearner’s intentional safety abortions on fragmented profiles, the Dual-Track architecture guaranteed complete clinical coverage. As highlighted in Supplementary Table S7, the native *`XGBoost`* module—which inherently routes missing variables without requiring prior synthetic imputation—stepped in as the universal fallback. The *`XGBoost`* track maintained an algorithmic penetrance of 100% (0 NAs), successfully generating valid clinical risk scores for every single patient that the SuperLearner safely masked. This Dual-Track validation proves that our pipeline can autonomously toggle between maximum ensemble precision for intact patient data, and robust, single-algorithm resilience for clinically fragmented data.

### 3.21. Clinical Probability Translation and Trajectory Bifurcation

To translate the validated continuous hazard Z-scores into actionable oncology metrics, the inference engine utilized geometric anchoring against the Phase III Breslow baseline hazards to extract individualized 1-year, 3-year, and 5-year clinical probabilities. To visualize the massive scale and stability of these 1,050 patient trajectories, we developed a 4-panel bifurcated composite model (Figure 11). To preserve mathematical clarity, the visualizations were strictly separated by endpoint physics.

**Figure 11.**
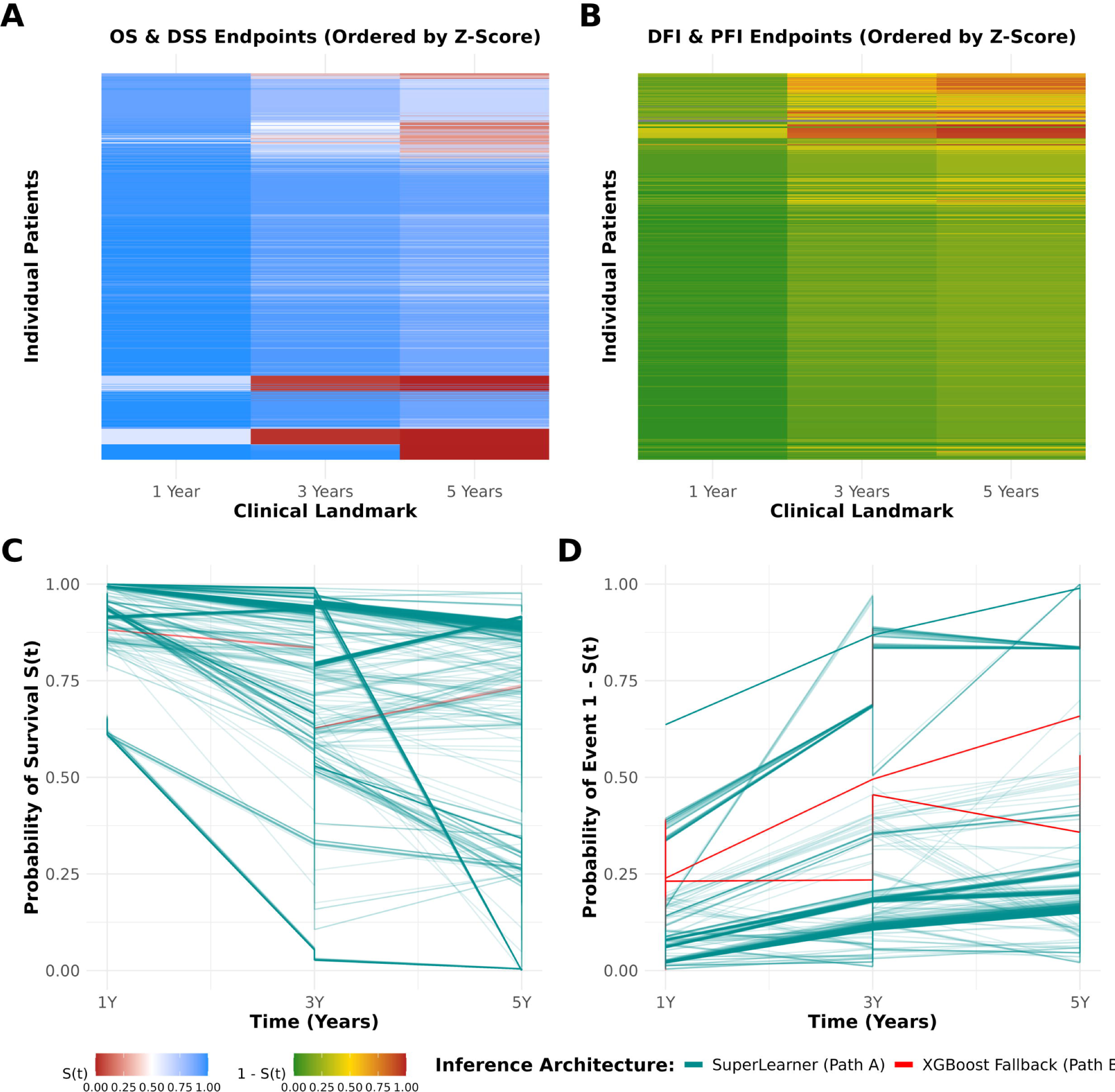
Internal Blind Validation and Trajectory Bifurcation of Dual-Track Clinical Probabilities. Individualized clinical probabilities (N = 1,050) were extracted across 1-year, 3-year, and 5-year clinical landmarks. The continuous hazard Z-scores from the inference engine were geometrically anchored against Phase III Breslow baseline hazards, maintaining strict topological integrity without requiring synthetic imputation. To preserve mathematical clarity, the visualization is strictly bifurcated by endpoint physics: (A) Waterfall heatmap of survival endpoints (OS and DSS) representing the decreasing probability of survival, *S(t)*. Patients are ordered vertically by their continuous risk Z-score to illustrate the smooth probability gradient across the cohort (gradient maps from 0% [Firebrick] to 100% survival [Dodger Blue]). (B) Waterfall heatmap of event endpoints (DFI and PFI) representing the increasing probability of recurrence or progression, *1 - S(t)*. The gradient is distinctly mapped to reflect the inverted physics of the endpoint (0% risk [Forest Green] to 100% risk [Firebrick]). (C, D) Temporal spaghetti plots mapping the individual probability trajectories for the Survival (C) and Event (D) endpoints over the 5-year clinical window. Trajectories are uniquely color-coded by the execution architecture utilized: patient records containing structurally intact multi-omic signatures that successfully navigated the SuperLearner (Path A) are plotted in Dark Cyan, whereas heavily fragmented patient records dynamically routed to the native *`XGBoost`* fallback (Path B) are plotted in Red. The seamless overlap of these trajectories demonstrates a complete absence of mathematical outliers and visually validates that the *`XGBoost`* fallback perfectly preserves prognostic scaling, guaranteeing stable algorithmic penetrance prior to clinical deployment.

The waterfall heatmaps (Figure 11A and Figure 11B) visually confirm the smooth gradient cascade of the individualized risk scores across the cohort. Specifically, Figure 11A illustrates survival endpoints (OS and DSS) as a decreasing probability of survival S(t), while Figure 11B illustrates event endpoints (DFI and PFI) as an increasing probability of recurrence or progression 1 - S(t).

More importantly, the temporal spaghetti plots (Figure 11C and Figure 11D) map the individual probability trajectories based on the algorithm utilized: SuperLearner (Path A) or *`XGBoost`* (Path B). Figure 11C tracks the decreasing survival probabilities, whereas Figure 11D tracks the escalating event risks over the 5-year clinical landmark. These overlapping trajectories demonstrate a complete absence of mathematical outliers and visually prove that the *`XGBoost`* fallback perfectly mirrors the prognostic scale of the SuperLearner. This ensures that heavily fragmented patient records yield unified and stable clinical predictions, fully validating the dual-track inference architecture prior to clinical deployment.

### 3.22. CancerRCDPredictor - A Parameterized Architecture for High-Dimensional Geometric Navigation and AI Diagnostic Synthesis

To translate the complex outputs of the SuperLearner predictive framework into an accessible, interactive platform, we developed the CancerRCDPredictor application (available at URL: https://lbtgenomica.uenf.br/cancerrcdpredictor/). The execution of the multi-algorithmic machine learning pipeline across 96 independent multi-omic cohorts generated a data repository exceeding 80 GB, comprising high-resolution mathematical proofs, diagnostic curves, and patient-level trajectory matrices. To facilitate the exploration of this massive dataset, the web application was engineered as a translational interface, systematically organizing the SuperLearner outputs into distinct, user-driven modules.

The application follows a "Pedagogical First, Data Explorer Second" structural design. To aid in the interpretation of high-dimensional non-proportional hazard functions and *`SHAP`* topologies, the platform anchors each capability module with an educational exemplar. Users are provided with static, manuscript-grade visualizations accompanied by executive summaries that explain the interpretation of multi-omic risk trajectories and functional interactions (e.g., lethal synergy versus protective antagonism).

Beyond the educational exemplars, the platform provides dynamic 96-Cohort Data Explorers. These parameterized engines allow users to selectively query the 59 GB repository by inputting specific cancer lineages and terminal clinical endpoints (e.g., Overall Survival, Disease-Specific Survival). The application is divided into four primary analytical modules, each serving a distinct scientific and clinical utility:

Global Impact (*`SHAP`* Beeswarms): This module visualizes the population-level influence of retained multi-omic signatures. It allows users to assess how the continuous expression or discrete presence of specific genomic elements drives non-proportional hazard projections across an entire cohort.

Interaction Topologies (Dependence): This module isolates the geometric relationships between interacting omic features. It provides high-resolution spatial mapping of synergistic, antagonistic, and bifurcating feature pairs, detailing how combined multi-omic states augment or suppress survival risk.

*`MVL`* Performance (Time-Dependent AUROC): This module evaluates the temporal stability of the SuperLearner framework. It outputs dual-panel ROC curves, tracking the absolute discriminatory capability (AUC) of the models at 1-year, 3-year, and 5-year clinical horizons to validate prognostic resilience over time.

Precision Oncology (Patient Trajectories) and AI Diagnostic Synthesis: Transcending standard individualized risk stratification, this module utilizes *`SHAP`* waterfall plots to deconstruct the predictive logic for single patients, mapping how specific molecular alterations push a patient’s trajectory toward lethal or protective outcomes relative to the population baseline. To fully operationalize this geometric data, the module features an integrated Large Language Model (LLM) acting as an automated Digital Tumor Board (Supplementary Note 11). This AI diagnostic engine autonomously extracts the Top 5 patient-specific *`SHAP`* topological coordinates, decoding the alphanumeric signatures back to their precise biological origins. The module meticulously cross-references each localized molecular driver against literature-verified Regulated Cell Death (RCD) pathways, specific multi-omic layers (e.g., CpG methylation vs. mRNA transcripts), and population-level tumor phenotypes. By dynamically synthesizing these multi-dimensional layers, the AI engine generates an audit-ready, continuous clinical narrative. Crucially, the diagnostic module mathematically contextualizes contradictory phenotypic correlations as profound multi-omic tumor heterogeneity rather than algorithmic noise, bridging the gap between high-dimensional machine learning and actionable clinical intelligence. Finally, an interactive clinical dialogue interface allows users to directly query the AI synthesis to explore tailored therapeutic vulnerabilities.

By compartmentalizing the data into these specialized modules, the architecture ensures that the geometric fidelity and algorithmic outputs of the SuperLearner framework can be systematically explored and utilized for both broad cohort analysis and individualized clinical evaluation.

## 4. Discussion

The pervasive reliance on strictly parametric survival frameworks in precision oncology (e.g., standard Cox PH regression and penalized linear formulations) frequently fails to capture the high-dimensional, nonlinear realities of multi-omic biological spaces. Furthermore, traditional approaches to multi-omic "integration" often mistakenly equate integration with modeling distinct omic layers separately and subsequently attempting to merge their fragmented results post-hoc. In this study, we operationalized true integrative multi-omics: the simultaneous, native estimation of predictive values across all seven molecular layers within a single, unified topological matrix. By physically isolating predictive models to localized cancer execution domains, our architecture prevented cross-cohort contamination and protected the clinical trajectory from the geometric instability common in generic pan-cancer amalgamations.

The absolute necessity of this approach was geometrically confirmed by our Phase II CANARY diagnostic. After evaluating the initial library of 120 algorithmic configurations and strictly disqualifying 24 structurally compromised matrices, testing the remaining 96 advanced cohorts under classical CoxNet constraints resulted in exactly zero strata surviving the proportional-hazards µ-ladder. This absolute collective structural failure across pan-cancer strata empirically proves that the standard reliance on linear, proportional-hazard assumptions in multidimensional omics is mathematically and biologically invalid; the native survival topology is fundamentally non-proportional.

Crucially, the deployment of the Phase III Quadripartite ML Ensemble demonstrates the inherent advantage of Stacked Generalization (SuperLearner) architectures over isolated model reliance (van der Laan et al., 2007, Naimi et al., 2018) Rather than claiming generalized algorithmic superiority, the SuperLearner functions explicitly as a robust anti-overfitting barrier. This topological stability was structurally anchored prior to overarching integration by enforcing a rigid 35% geometric missingness boundary, surgically excising clinical "ghost" variables (vectors presenting with fundamentally hollow predictor domains) to guarantee that all downstream model convergence occurred strictly upon genuine biological mass. This directly intercepts the severe geometric instabilities and geometric calibration failures endemic to high-dimensional "pan-cancer" modeling, vulnerabilities which have been rigorously documented in recent large-scale benchmarking frameworks (e.g., the SurvBoard consortium; (Wissel et al., 2025). Tree-based paradigms, such as Sparsity-Aware *`XGBoost`*, while excelling at mapping high-dimensional spaces, routinely suffer from extreme variance inflation when forced to biological structural limits. By deploying an *`MVL`* ElasticNet to systematically synthesize the divergent inductive biases of the four foundational architectures— *`RSF`*, Sparsity-Aware *`XGBoost`*, insulated Survival-*`Boruta`* topological parameters, and *`MTLR`*—the architecture naturally penalizes isolated mathematical outliers. This "algorithmic stacking" mitigates the catastrophic effects of multi-omic collinearity, compressing the overarching hazard predictions into a highly stabilized, cross-validated consensus space that successfully prevents the severe temporal model decay inherent to proportional-hazards frameworks, seamlessly maintaining robust resolution across 1-, 3-, and 5-year clinically truncated progression horizons. Furthermore, the SuperLearner was fundamentally engineered to execute a devastating "Topological Floor Defense." Operating as an architectural "No One Stays Behind" policy, the Elastic Net synthesis autonomously rescued highly fragmented, chaotic biological spaces—such as the lush LGG multi-omic topography—by distributing democratic trust across the base-learners. This mathematically protected deeply volatile matrices from crashing into stochastic noise, rigidly anchoring the pan-cancer survival floor at a globally predictive 0.526 baseline regardless of individual algorithmic failure.

This strict simultaneous integration actively forced a competitive evaluation between distinct cancer omic networks, exposing a profound mathematical phenomenon: the systematic, algorithmic displacement of somatic mutations and CNVs by downstream dense continuous multi-omic features. This massive displacement aligns precisely with established machine learning observations regarding ‘cardinality bias’ within decision-tree logic (Strobl et al., 2007). Tree-based learning arrays inherently maximize predictive entropy. A binary mutation (present/absent) offers only a single, blunt bifurcation point, lacking the mathematical flexibility required to track a patient’s longitudinal, continuously compounding survival hazard over multi-year clinical timelines. Conversely, continuous molecular profiles—such as mRNA abundance, transcript isoform gradients, and epigenetic methylation—natively provide thousands of micro-partitions. Specifically, binary variables frequently suffered from localized intra-cohort variance starvation; lacking sufficient numerical deviation within strictly bound execution strata, they yielded negative Information Gain and were mechanically ejected by the gradient boosting paradigms. Consequently, because they offer the algorithmic trees an exponentially denser spectrum of continuous hazard gradients, active continuous multi-omic arrays mathematically displaced static baseline genotypes during overarching multi-omic competition. This algorithmic displacement rigorously mirrors recent multi-omic predictive evaluations, which consistently demonstrate that continuous multi-omic strata map the functional tumor state so comprehensively that enforcing the integration of sparse, static mutations routinely yields mathematically negligible downstream prognostic gains (Herrmann et al., 2021). While exactly 142 mutations and 196 CNVs successfully established baseline viability within the 12,613 prognostic pool, when forced through the absolute apex limits of the Quadripartite gauntlet, zero genotypic markers natively remained among the ultimate 150 pan-cancer golden anchor concordances. These 150 anchors do not represent absolute biological superiority; rather, they identify the precise subset of features possessing a continuous topological geometry robust enough to guarantee systematic retention across all four highly distinct Machine Learning architectures simultaneously.

Crucially, this overarching mathematical suppression must not be conflated with biological inertness. Our deliberate deployment of the Sparsity Isolation Protocol Sparsity Isolation framework explicitly tested discrete binomial variables separated entirely from continuous phenotypic matrices. This framework definitively proved that baseline somatic mutations and CNVs possess highly significant independent prognostic trajectories natively within their isolated architectures. Their subsequent suppression in the fully integrated Terminal Harvester structure functionally confirms that highly dense, continuously active RNA phenotypes effectively "mask" sparse binary signals in joint predictive arrays. However, this suppression is not absolute. As definitively documented via the topological "Sparsity Rescue Effect," when severe cohort-specific data missingness causes these dominant localized transcriptomic cascades to structurally collapse, the SuperLearner autonomously pivots. It mechanically falls back upon the underlying somatic distributions to rescue the survival bounds, confirming that discrete genotypic signatures fundamentally function as essential architectural fail-safes when the primary continuous phenotype is compromised.

Ultimately, computational predictive accuracy remains clinically inert without actionable, transparent biological interpretability. Traditional biomarker analyses routinely extract a unified "top 10" signature and universally apply it, falsely assuming that a generalized cohort-level ranking accurately defines every distinct patient’s risk. Our integration of post-hoc exact *`TreeSHAP`* and *`LIME`* architectures explicitly dismantles this generalization (Lundberg et al., 2020). While sparse surrogate models (*`LIME`*) were deployed to map restricted individual boundaries, their routinely suppressed Explanation Fits (R² < 0.10) provided explicit mathematical proof that single-patient survival boundaries remain too highly nonlinear for any linear proxy to fully navigate. Consequently, while the SuperLearner defined overarching global hierarchies, extracting algorithmically localized (N × N) interaction tensors and patient-specific waterfall trajectories from the exact *`TreeSHAP`* architecture proved that individual patient survival is fundamentally idiosyncratic. This individualized focus directly responds to the urgent mandate in contemporary precision oncology to abandon generalized algorithmic opacity in favor of direct, patient-specific feature attribution frameworks that secure actionable clinical translation (Klauschen et al., 2024). We documented that what severely accelerates a lethal trajectory in one patient is frequently dictated by a highly distinct, personalized set of multi-omic features (e.g., specific mRNA overexpression synergizing with epigenetic layers) that may be entirely dormant or irrelevant in another patient harboring an equal mortality risk. Furthermore, this localized tracking dynamically diagnoses whether mortality is driven by an isolated, targetable biological anomaly (an apex driver) or dictated by diffuse polygenic degradation—where hundreds of minor multi-omic aberrations synergize into a massive cumulative hazard penalty. Most profoundly, dissecting these multi-dimensional *`TreeSHAP`* topographies exposed the mechanical phenomenon of "Antagonistic Rescue" (Figure 7; Figure 8, Panel B). While classical models assume survivability is dictated by simply summing isolated risk biomarkers, the Quadripartite architecture proved that the localized presence of an apex lethal mRNA signature can be mathematically and biologically neutralized if the patient concurrently possesses an intersecting, protective transcript isoform. This capability to natively resolve cross-layer rescue capabilities signifies the structural invalidation of linear additivity; it confirms that true precision oncology forecasting hinges entirely on synergistic and antagonistic multi-omic interactions. By mathematically singularizing the importance of variables at the absolute patient level—revealing exactly how varying omic permutations combine to drive lethal or protective deviations from the baseline—our architecture successfully shifts predictive modeling from a generalized cohort stratification tool into a probabilistic, high-resolution instrument for precision oncology interception.

However, the ultimate realization of this precision oncology framework requires rigorous context regarding its clinical generalizability. While our partitioned, blinded internal validation strategy successfully insulated the predictive architecture from information leakage and maintained strict 7-layer multi-omic harmony—ensuring the models were evaluated on genuine biological accuracy rather than their tolerance for disparate external preprocessing techniques—this approach inherently lacks multi-institutional external testing. In precision oncology, external validation remains the gold standard for proving true cross-demographic generalizability. Given the current global scarcity of fully independent, comprehensive multi-omic public datasets spanning 33 distinct tumor types, our internal blinded approach serves as a mathematically robust and highly controlled diagnostic intermediate. Future large-scale, multi-institutional sequencing efforts will be essential to definitively establish the universal clinical portability of these personalized multi-omic trajectories.

The clinical integration of the CancerRCDPredictor platform provides several critical advantages for precision oncology, overcoming the cognitive and interpretive barriers commonly associated with high-dimensional machine learning. Primarily, the application achieves automated multi-omic translation by seamlessly bridging the gap between complex mathematical algorithms and clinical biology, converting raw *`SHAP`* topologies into readable, logically coherent patient diagnostic narratives. This functionality operationalizes Explainable AI (XAI) in practice; rather than providing an opaque ‘black-box’ survival score, the platform empowers oncologists by explaining precisely why a prediction was made, isolating the exact genetic, transcriptomic, or epigenetic drivers responsible for altering patient risk. Furthermore, the architecture is uniquely capable of contextualizing tumor heterogeneity. By biologically reconciling seemingly contradictory statistical data—such as a gene exhibiting lethal effects via transcript expression but protective effects via methylation—the platform helps clinicians practically grasp dynamic tumor plasticity and multi-pathway resistance mechanisms. Ultimately, this structural design ensures guided clinical actionability. By highlighting a patient’s dominant RCD pathways and actionable metabolic or stromal vulnerabilities, the integrated AI generates tailored follow-up queries, providing oncologists with immediate, data-driven starting points for targeted therapy discussions.

## STATEMENTS

### Ethics approval and consent to participate

Ethics approval is not applicable to this study.

### Data Availability Statement

The datasets generated and analyzed for this study, along with the supplementary tables, high-resolution figures, and source code that support the findings, are too large to be hosted as standard supplementary material on the journal’s submission platform. Therefore, this complete dataset and codebase are publicly hosted and fully accessible at the following GitHub repository: https://github.com/BioCancerInformatics/CancerRCDPredictor/ and can be explored and download from the CamcerRCDPredictor shiny application available at URL: https://lbtgenomica.uenf.br/cancerrcdpredictor/.

### Author contributions

E.R.S. conceptualized the study; developed, validated, and implemented the R-based computational pipeline; performed integrative analyses; contributed to manuscript revision; and developed the Shiny application and GitHub repository.

H.A.C.N. contributed to pipeline development and code validation; generated figures; and participated in manuscript editing, and developed the Shiny application and GitHub repository.

V.S.L. generated figures and contributed to manuscript editing.

E.M.A. conceptualized and supervised the study; contributed to methodological design; developed, validated, and audited the computational pipeline; and contributed to writing and critical revision of the manuscript.

## Acknowledgements

The results shown here are based on data and resources generated by the TCGA Research Network (https://www.cancer.gov/tcga), UCSC Xena (https://xena.ucsc.edu), UCSC Xena Shiny (https://shixiangwang.shinyapps.io/ucscxenashiny/), CancerRCDShiny (https://cancerrcdshiny.shinyapps.io/cancerrcdshiny/), and KEGG (https://www.kegg.jp).

## Consent for publication

Not applicable.

## Generative AI statement

Generative artificial intelligence tools were utilized in a strictly supervised capacity during manuscript and data preparation. ChatGPT (OpenAI; model version: GPT-5.2) was used to assist with English language revision, grammar, and the initial drafting of preliminary R scripts. Furthermore, the Antigravity AI agent (Google DeepMind, Gemini 3.3 Pro High and Claude Sonner 4 Thinking) was explicitly employed to programmatically validate the global numerical integrity of the manuscript—auditing all counts, percentages, and matrix dimensions across figures, main text, and supporting datasets for rigorous consistency. All AI-generated scripts were manually verified and incorporated by the authors into a fully controlled analytical pipeline. The authors assume full responsibility for the final content of the manuscript, including textual accuracy, code precision, and the validity of all conclusions.

## Publisher’s note

All claims expressed in this article are solely those of the authors and do not necessarily represent those of their affiliated organizations, or those of the publisher, the editors and the reviewers. Any product that may be evaluated in this article, or claim that may be made by its manufacturer, is not guaranteed or endorsed by the publisher.

## Supplementary material

The Supplementary Material for this article can be found online at: https://github.com/BioCancerInformatics/CancerRCDPredictor/.

## Significance Statement

Despite the proliferation of machine learning in precision oncology, traditional parametric frameworks fundamentally fail to accurately resolve high-dimensional, non-linear multi-omic survival boundaries. Furthermore, inherent algorithmic opacity has severely constrained the translation of complex biological dynamics into point-of-care interventions. Here, we present a zero-leakage Quadripartite ML Ensemble engineered specifically to dismantle restrictive proportional-hazards assumptions. Operating as a robust topological floor defense, this cross-validated SuperLearner executed an architectural 35% geometric missingness boundary, physically rescuing highly chaotic biological matrices from collapsing into stochastic noise. Crucially, by computationally forcing multi-omic competition, we mapped a structural algorithmic displacement, empirically demonstrating that dense continuous multi-omic networks mathematically suppress the predictive validity of static genomic sequences over prolonged clinical horizons. Finally, unraveling this non-linear topography via exact N-dimensional *`TreeSHAP`* dependencies organically exposed complex trans-layer synergistic and antagonistic rescue trajectories—proving that lethal omic signatures are frequently neutralized by interacting protective multi-omic networks. Collectively, this audit-compliant framework effectively terminates the era of generalized cohort stratification, forcibly shifting predictive oncology toward mathematically undeniable, individualized patient interception.

---

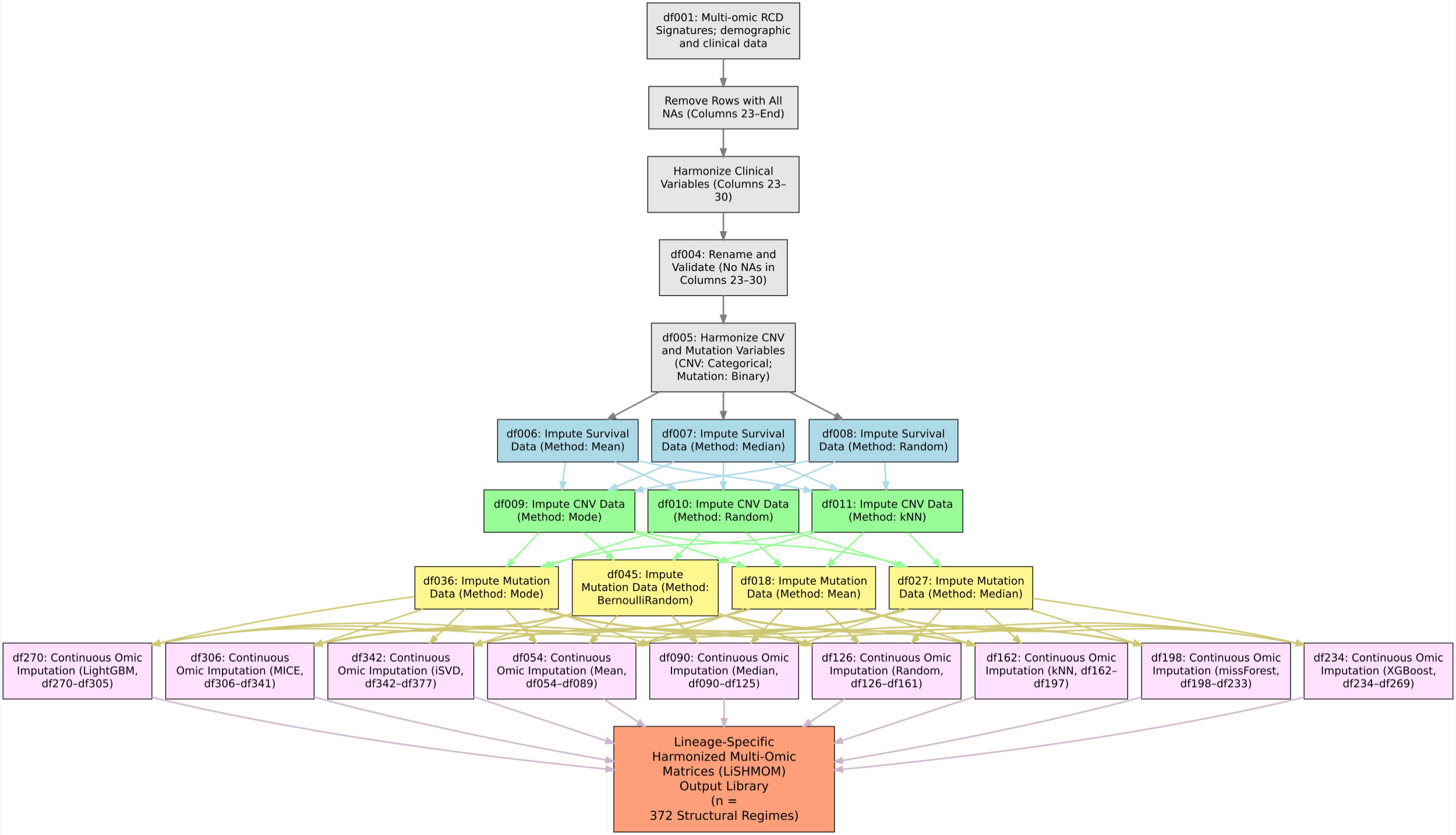
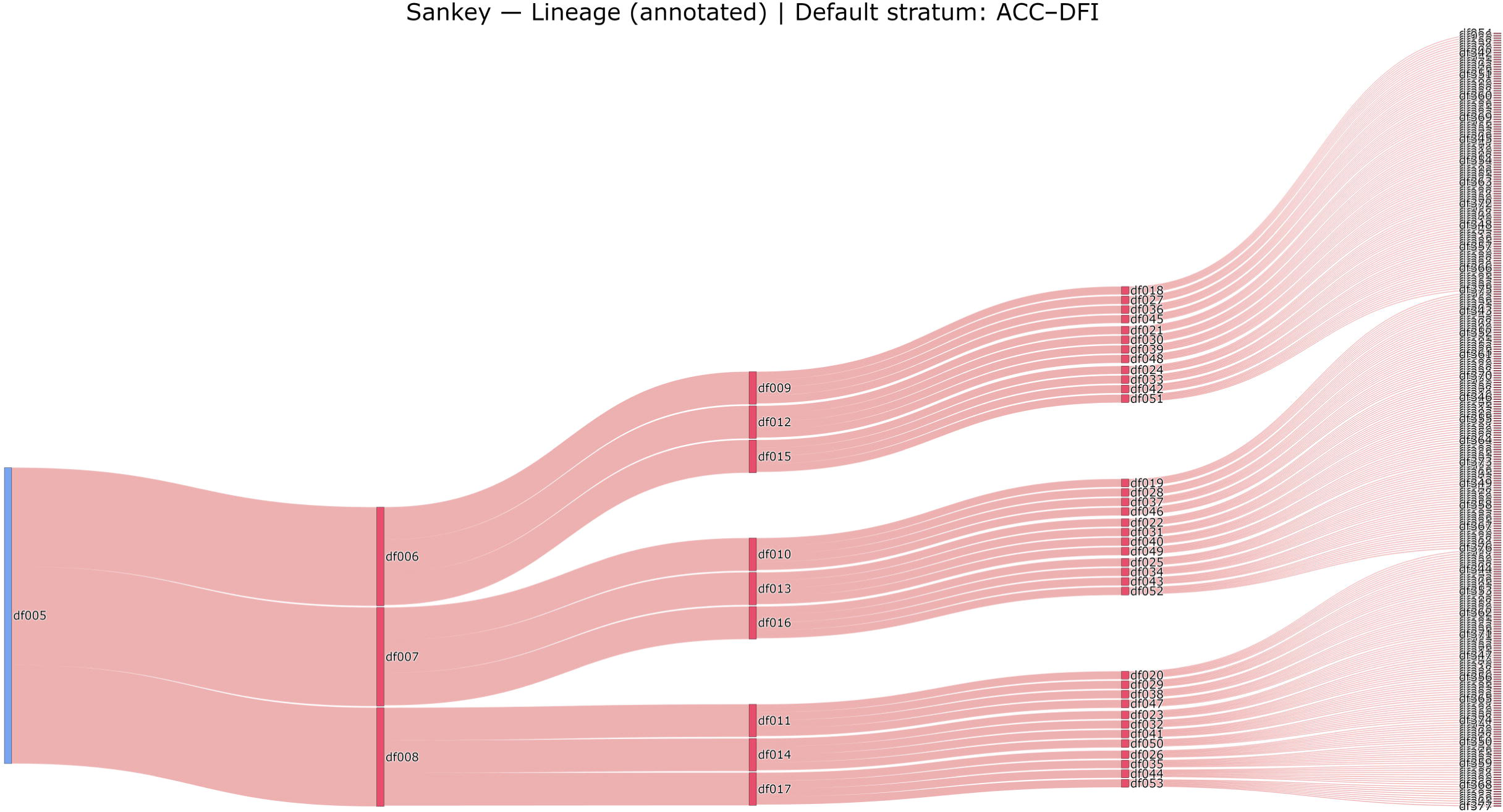
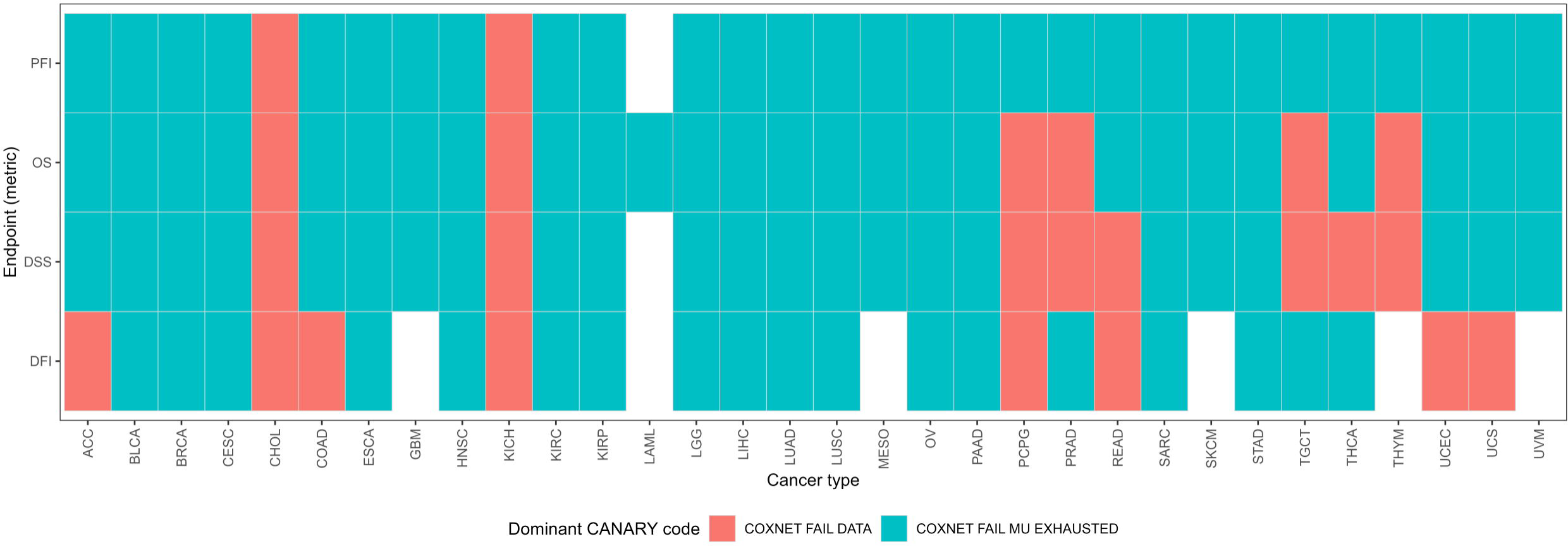
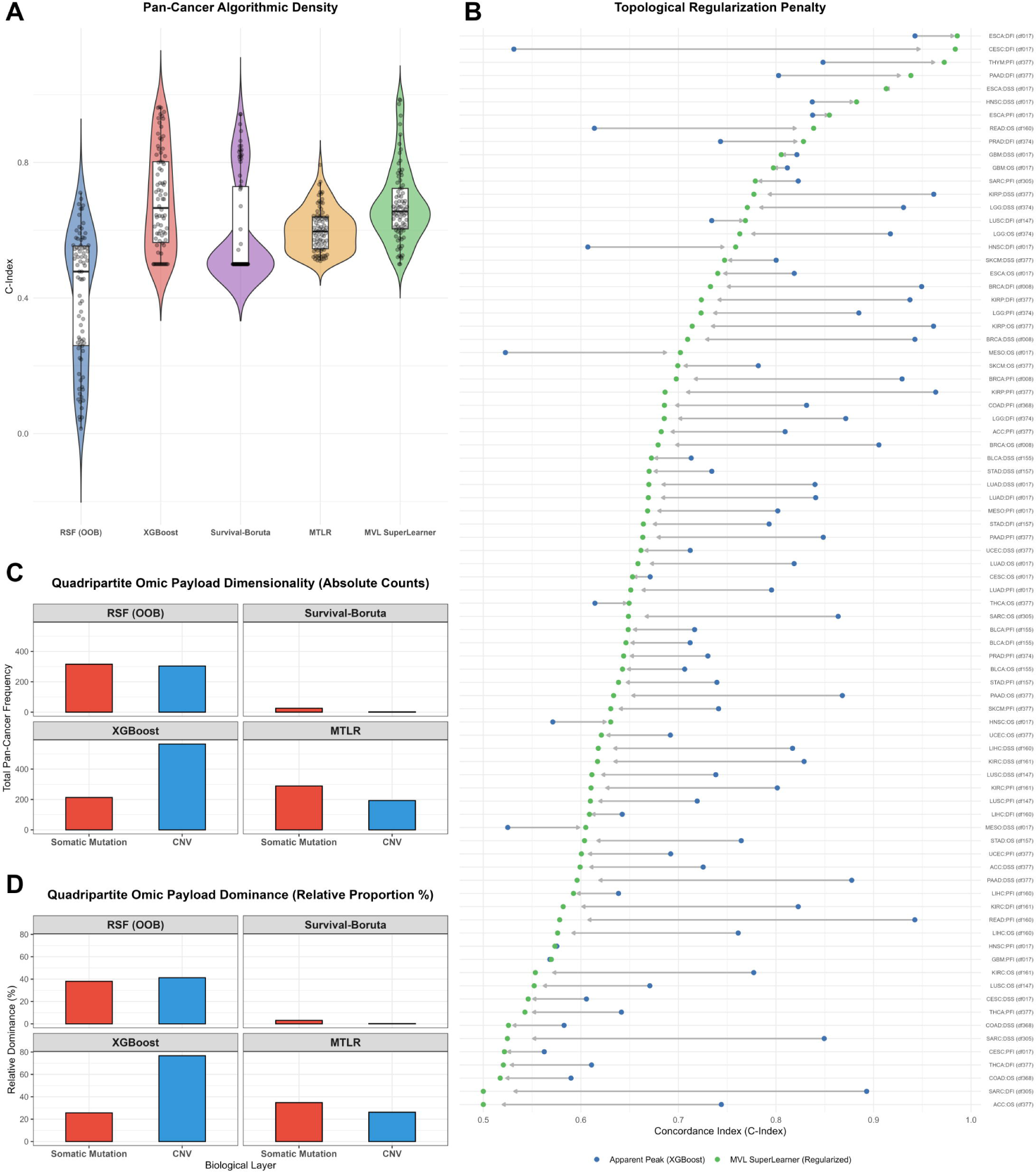
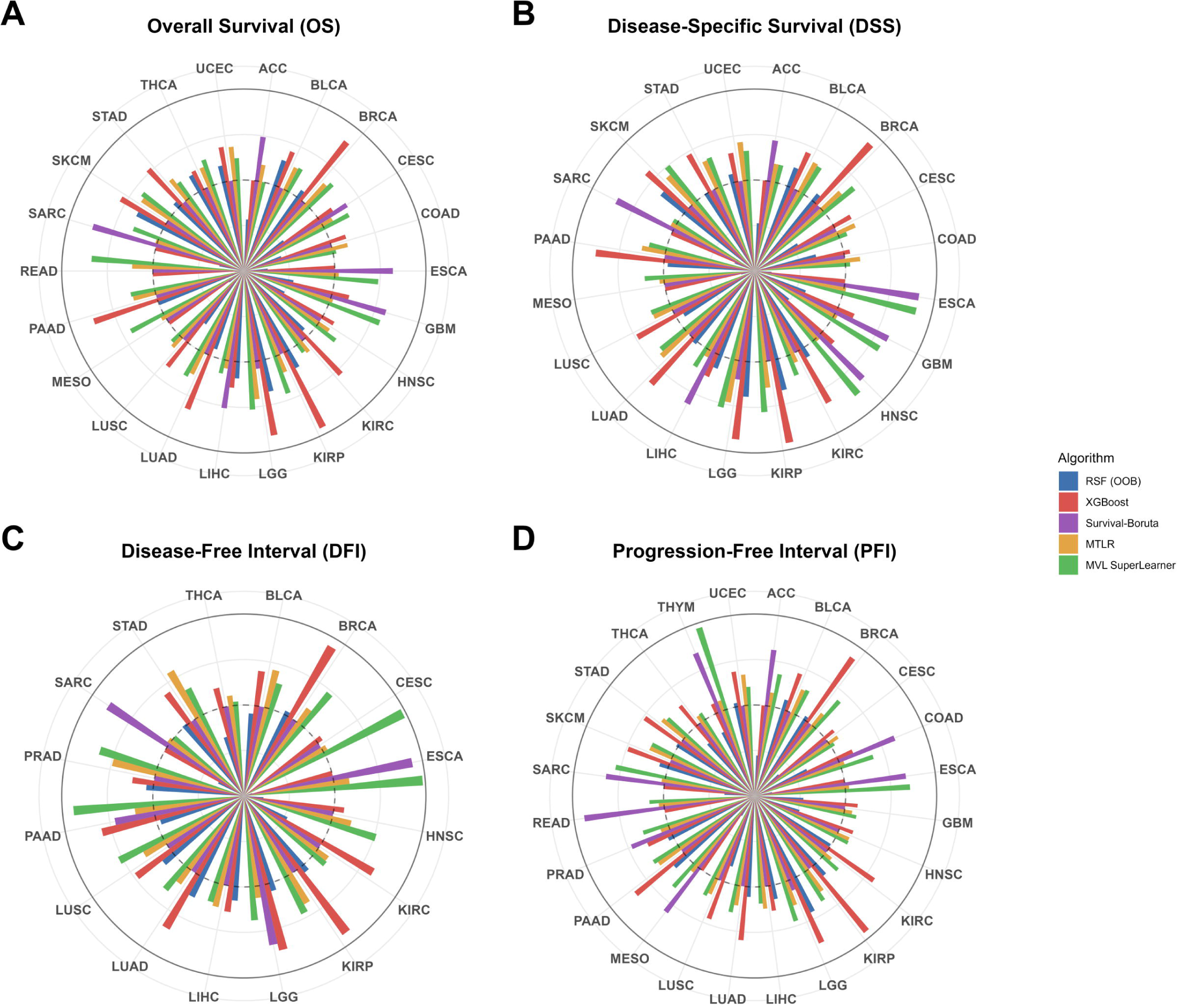
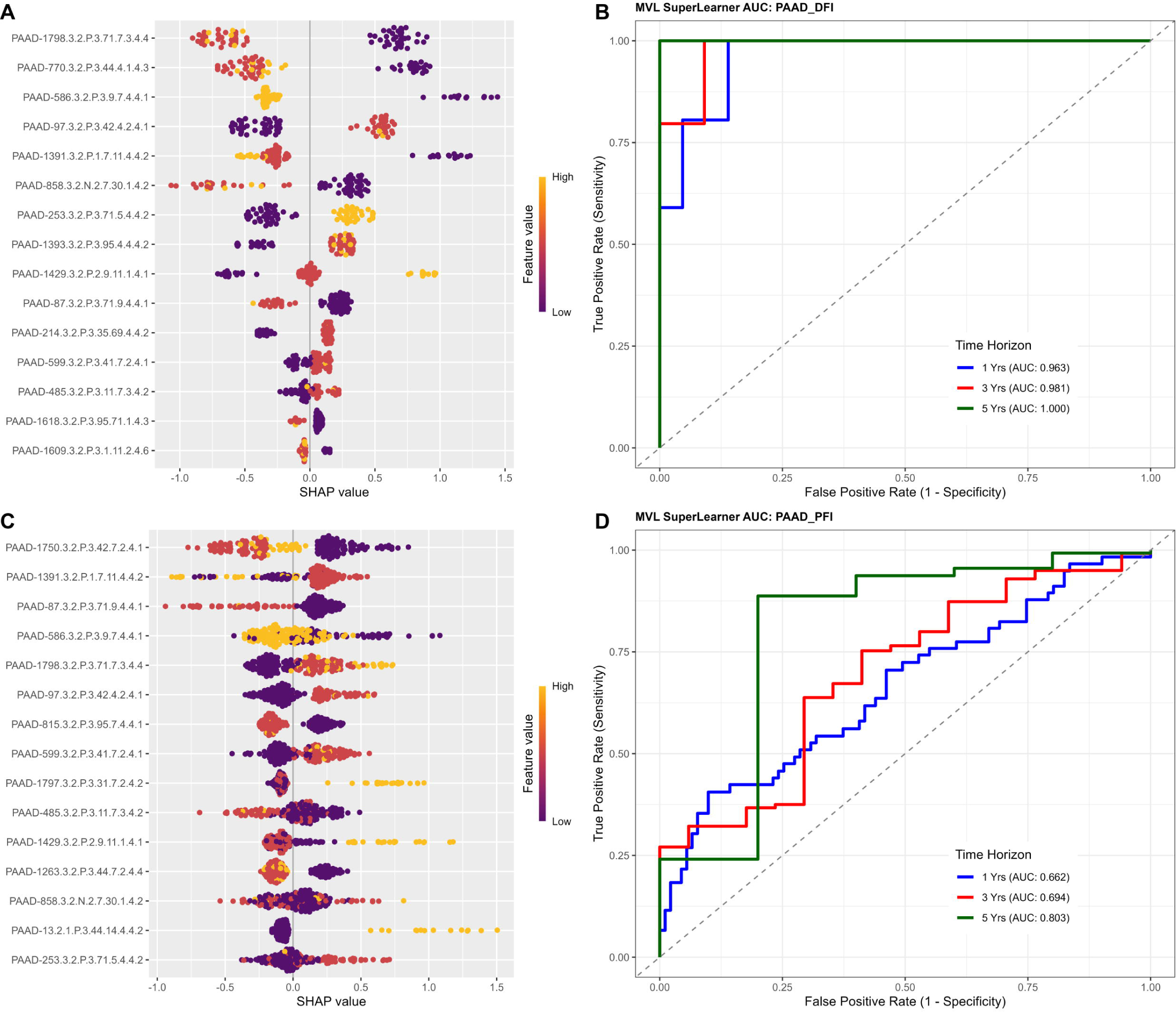
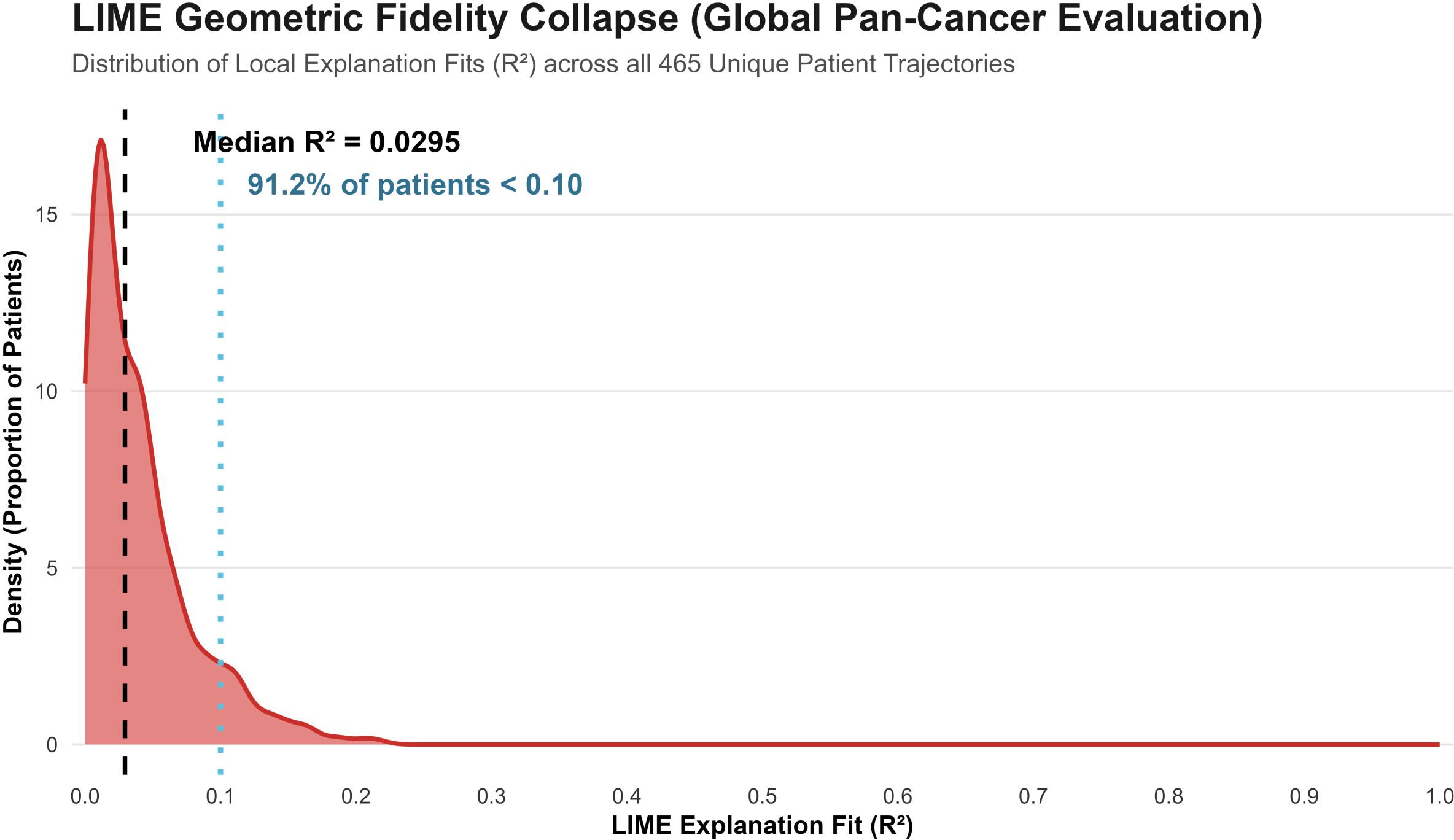
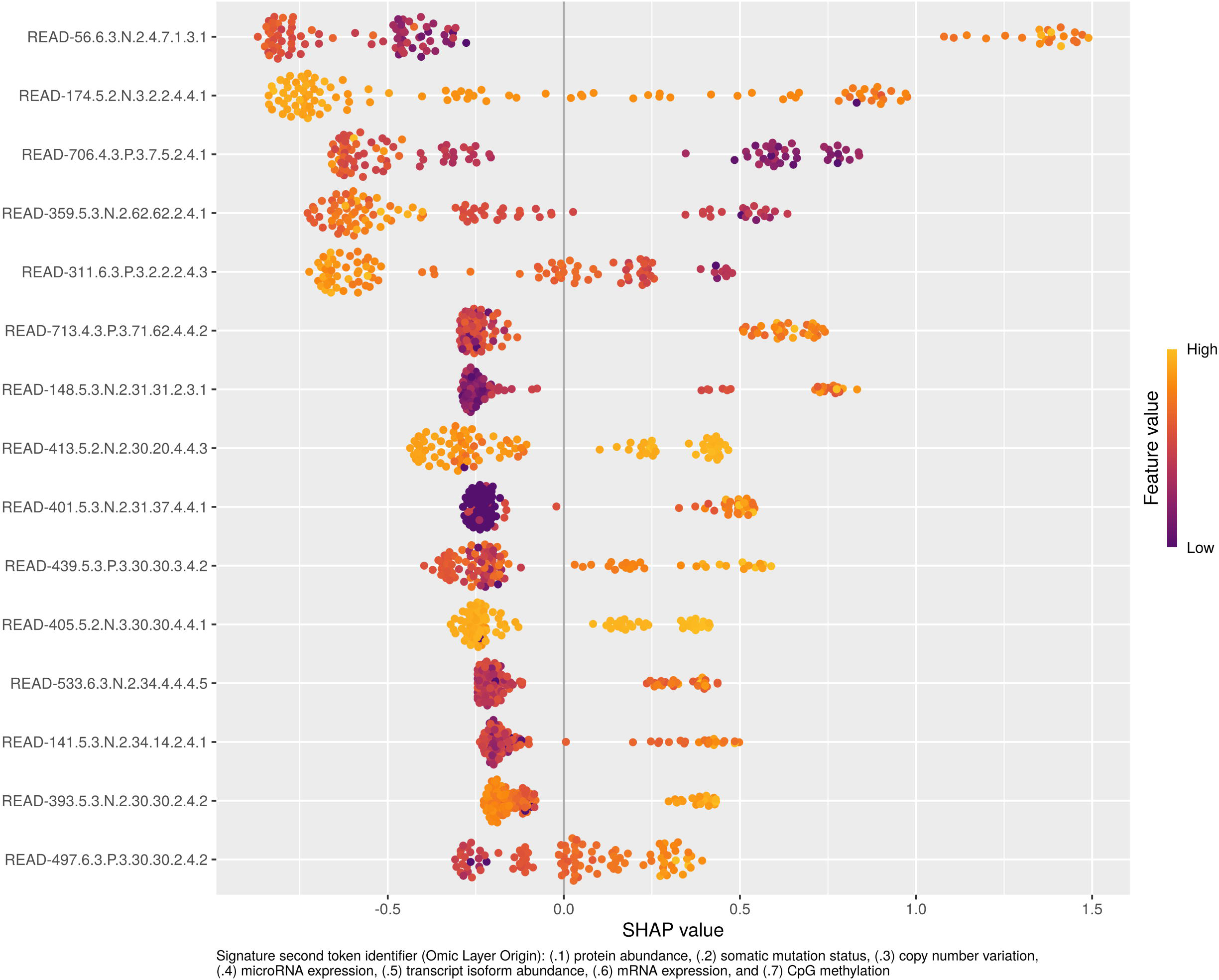
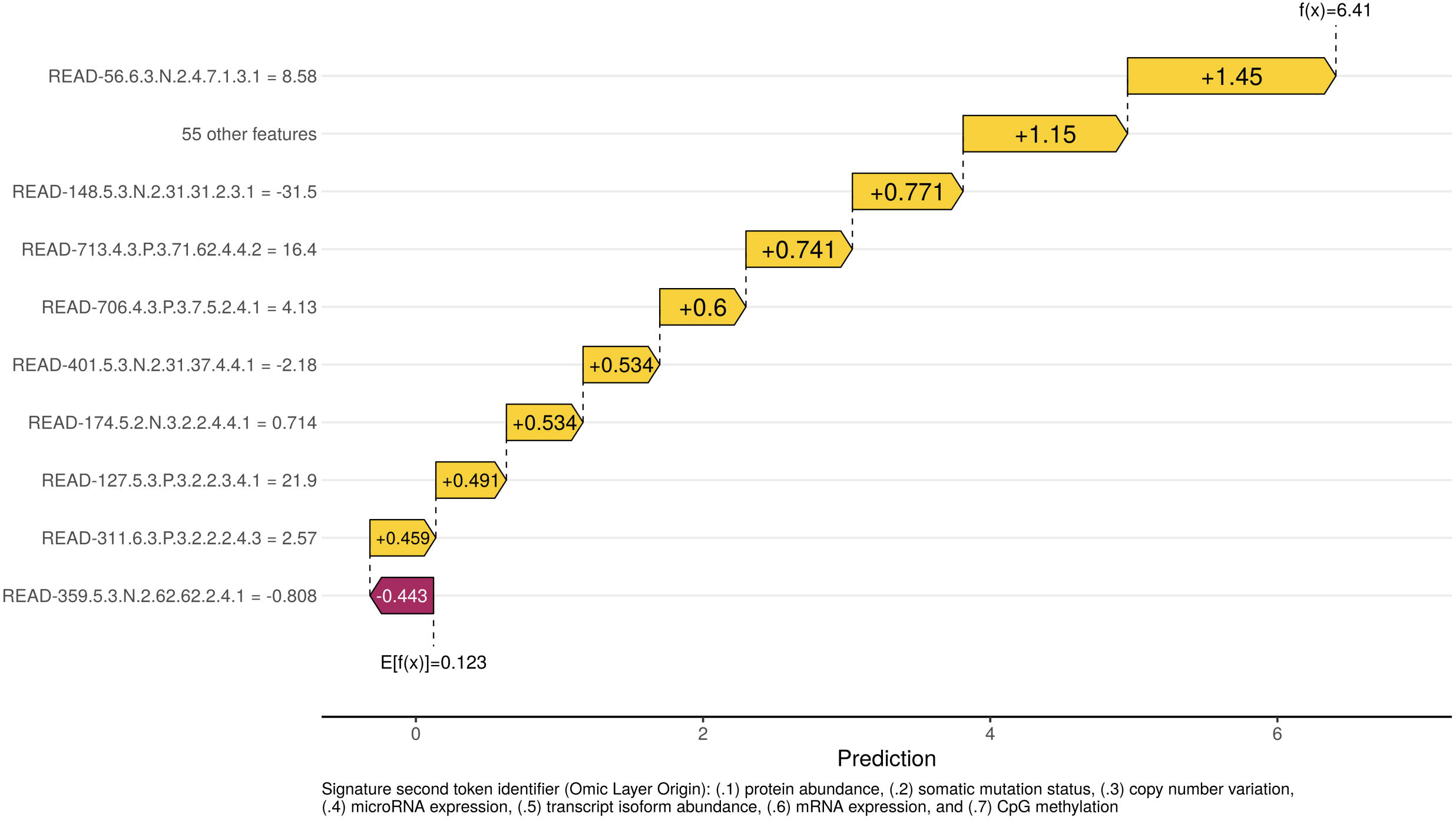
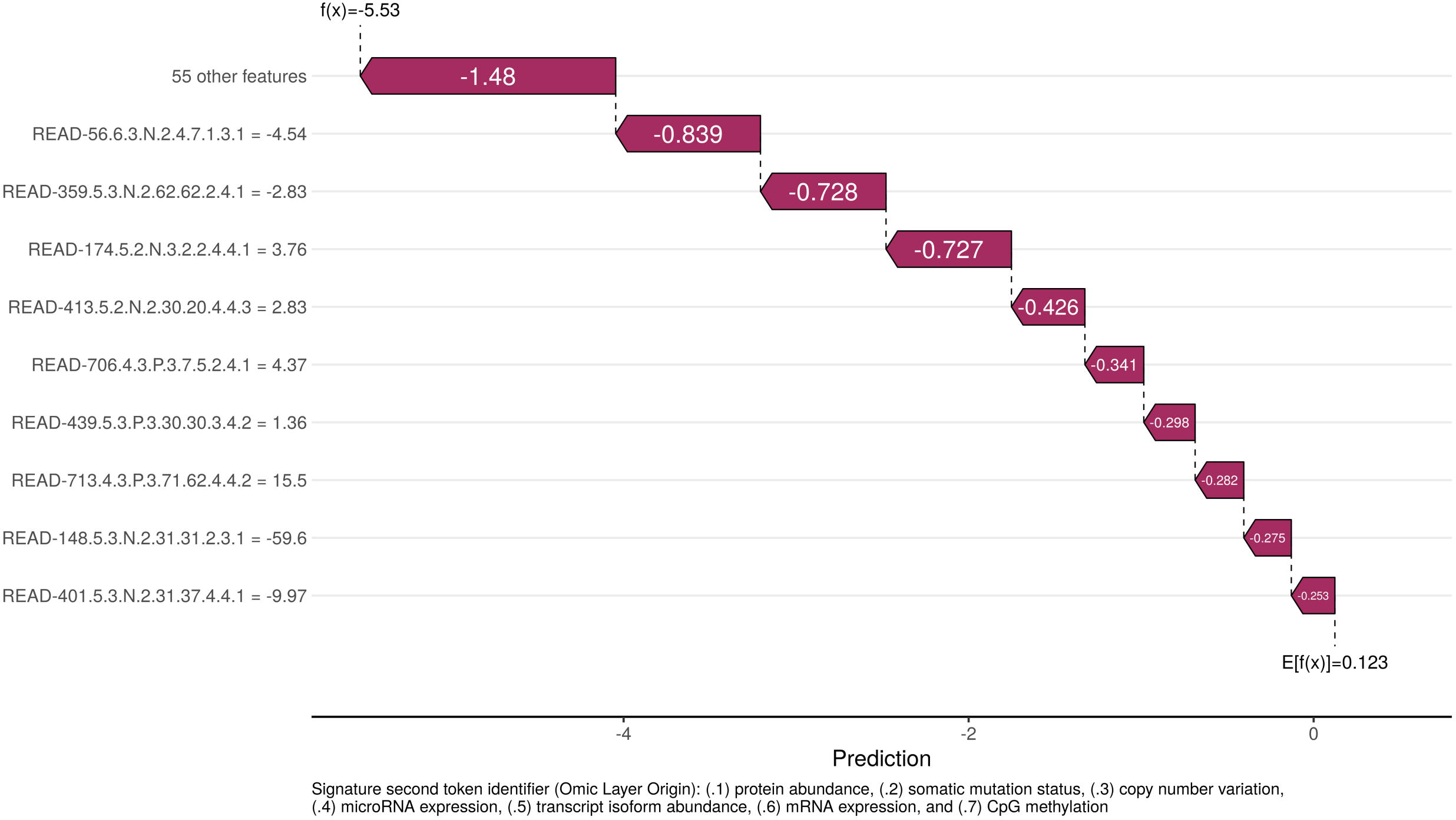
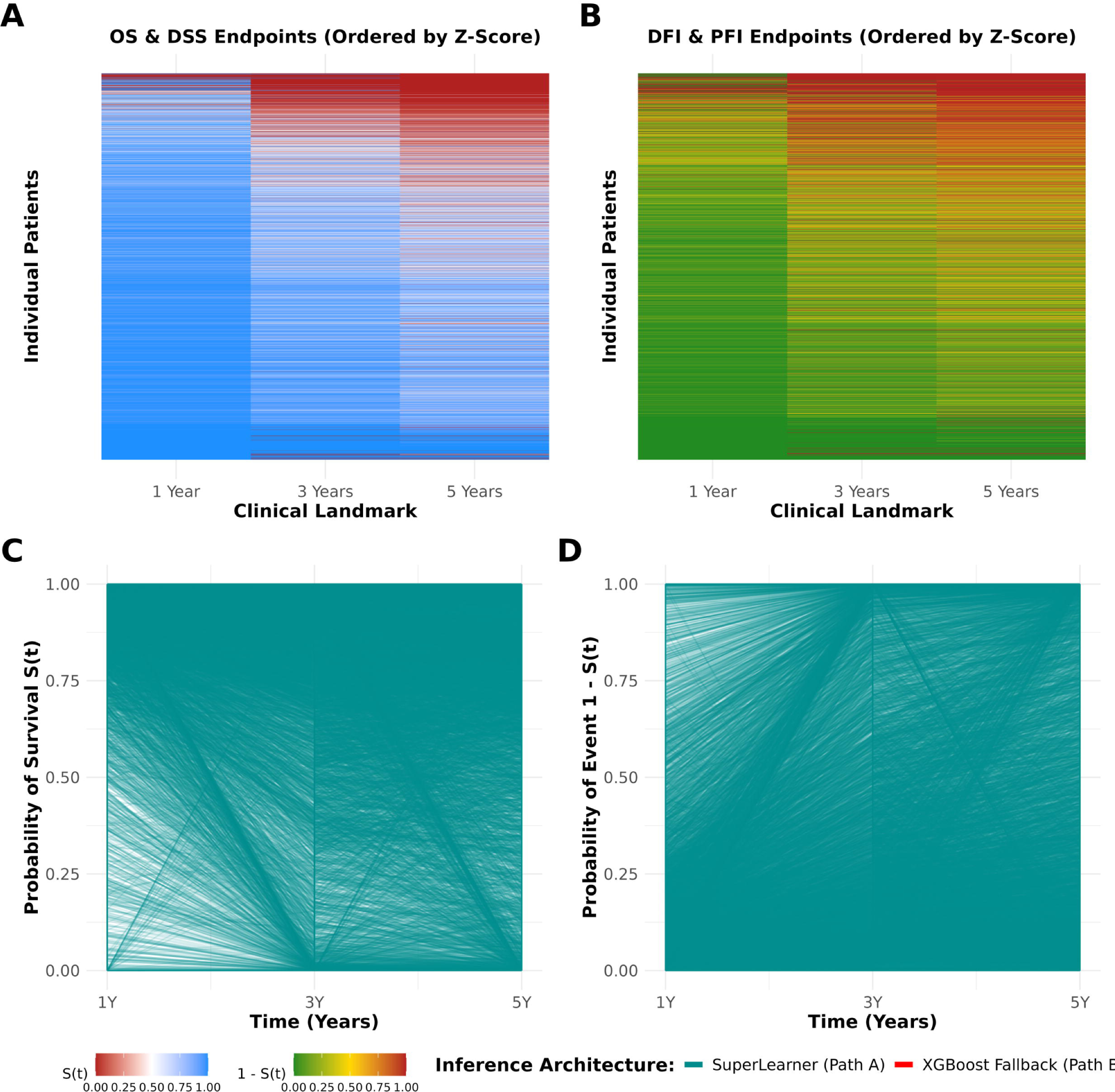

