## Supplementary Information for "A Pan-Cancer Multi-Omic SuperLearner for Regulated Cell Death Survival Topologies"

**Running Head:** Multi-Omic Precision Oncology Topologies

Emanuell Rodrigues de Souza, Higor Almeida Cordeiro Nogueira, Victor dos Santos Lopes and Enrique Medina-Acosta*

* **Corresponding author**: Enrique Medina-Acosta

**Address**: Laboratório de Biotecnologia, Centro de Biociências e Biotecnologia, Universidade Estadual do Norte Fluminense, Avenida Alberto Lamego 2000, Parque Califórnia, Campos dos Goytacazes, RJ, CEP 28015-602, Brazil.

| **Author** | **E-mail** |
|---|---|
| Emanuell Rodrigues de Souza | |
| Higor Almeida Cordeiro Nogueira | |
| Victor dos Santos Lopes | |
| Enrique Medina-Acosta* | |


### Supplementary Information

#### Supplementary Note 1. Methodological Framework for Multi-Omic and Multi-Optosis Signature Nomenclature

To ensure absolute precision and systematic retrieval of pan-cancer data, we adapted the standardized alphanumeric nomenclature system previously established by (Rodrigues de Souza et al., 2025). This structured taxonomy explicitly links the diverse multi-omic attributes of specific target genes to their respective phenotypic traits across 33 unique malignancies. The comprehensive identifier is constructed from an 11-part architectural array formatted as: **CTAB-GSI.GFC.PFC.SCS.TNC.HRC.SMC.TMC.TIC.RCD** (e.g., KIRP-107.3.2.N.1.44.44.1.1.2) (Figure 2).

The individual components of this topological array are defined as follows:

**CTAB**: A formalized three- or four-letter acronym designating the specific TCGA-derived cancer cohort (e.g., KIRP corresponding to kidney renal papillary cell carcinoma; refer to Supplementary Table S1 for the complete glossary).

**GSI**: A unique numerical identifier (ranging from one to four digits) assigned to distinct signatures within a given cancer type (e.g., 107).

**GFC:** The Genomic Feature Contexture, cataloging the biological layer of the signature: 1 (Protein expression), 2 (Somatic Mutations), 3 (Copy Number Variation [CNV]), 4 (microRNA expression), 5 (Transcripts), 6 (mRNA expression), and 7 (CpG Methylation).

**PFC**: The Phenotypic Feature Contexture mapping the biomarker to specific profiles: 1 (Tumor Mutational Burden [TMB]), 2 (Microsatellite Instability [MSI]), or 3 (Tumor Stemness Measure [TSM]).

**SCS**: The Spearman Correlation Sign, mathematically represented as either P (indicating a positive correlation) or N (indicating a negative correlation).

**TNC**: The Tumor versus Non-Tumor Expression Contexture, indicating comparative differential expression levels: 0 (insufficient data), 1 (stable/unchanged expression), 2 (significant underexpression), and 3 (significant overexpression).

**HRC**: The Hazard Ratio Contexture, formulated as a synthetic 1N2N3N4N array. The prefix digits (1 through 4) sequence the survival metrics—DSS, DFI, PFI, and OS, respectively. The suffix variable 'N' denotes the hazard magnitude, categorized as A (neutral/no effect), B (deleterious/risky), or C (favorable/protective).

**SMC**: The Kaplan-Meier Survival Distribution Contexture, which similarly adopts the 1N2N3N4N architecture across the four survival metrics. However, the qualitative classifiers (A, B, C, D) are uniquely calibrated according to the specific omic layer. The 'A' classification universally signifies "Not Significant" (NS) across all modalities. The 'B' classification denotes "High" expression for continuous modalities (Protein, miRNA, Transcript, mRNA, Methylation), but translates to "Mutant" (MT) for mutations and "Deleted" for CNV. Conversely, the 'C' classification indicates "Low" expression for continuous variables, "Wild Type" (WT) for mutations, and "Duplicated" for CNVs. The 'D' classifier is exclusively reserved for CNV signatures to represent a composite "Deleted/Duplicated" event.

(**Note**: The theoretical 128 topological combinations derived from the 1N2N3N4N mathematical arrays are programmatically reassigned to a continuous integer scale from 0 to 127 (Dataset S1J from (Rodrigues de Souza et al., 2025)). For example, an array of 1A2A3A4A (universal non-significance) is mapped to identifier 0, whereas 1A2A3A4B (exclusive risk in OS) maps to identifier 1).

**TMC**: The Tumor Microenvironment Contexture, evaluating the ecological niche: 1 (anti-tumoral), 2 (dual-role), 3 (pro-tumoral), and 4 (insufficient data).

**TIC**: The Tumor-Infiltrating Lymphocyte Contexture, characterizing immunological infiltration: 1 ("hot"/inflamed), 2 (variable), 3 ("cold"/desert), and 4 (insufficient data).

**RCD**: A one- or two-digit integer capturing the absolute number of distinct Regulated Cell Death (RCD) pathways associated with the signature.

**Supplementary Note 2: Structural Dimensionality and Mathematical Justification for Non-Proportional Modeling**

#### 1. Structural dimensionality of the multi-omic survival design space

Each TAR-admissible preprocessing regime represents a high-dimensional survival design matrix defined by the joint configuration of clinical covariates, multi-omic predictors, and endpoint-specific survival information. Across the study, analyses span 33 cancer types and four survival endpoints (OS, DSS, DFI, PFI), yielding cancer–endpoint strata that differ substantially in sample size, event burden, censoring structure, and predictor dimensionality. Crucially, all dimensional configurations and subsequent structural evaluations were executed strictly groupwise (isolated by cancer type and endpoint) to guarantee zero inter-lineage biological leakage or spatial covariance dilution. The baseline reference dataset (df005), which anchors all downstream preprocessing regimes, already comprises thousands of molecular predictors distributed across seven omic layers alongside clinical covariates, with substantial heterogeneity in missingness patterns across both variables and cancer types.

Predictor dimensionality and information content vary not only across cancers and endpoints, but also across preprocessing regimes. Different omic layers contribute markedly different numbers of predictors, exhibit distinct missingness profiles, and encode signals with varying degrees of collinearity and cross-layer dependence. Continuous expression layers (mRNA, transcript isoforms, protein abundance, methylation) are high-dimensional and strongly correlated, whereas mutation and CNV layers are sparse, discrete, and cancer-type dependent. Even prior to imputation, these characteristics induce non-trivial covariance structure and effective dimensionality that cannot be reduced to simple feature counts.

Missing data are likewise heterogeneous: individual predictors and entire omic layers may be well measured in some cancer types and nearly absent in others. Consequently, each preprocessing regime yields a distinct predictor geometry defined not only by retained feature counts, but by correlation structure, effective rank, and interaction with endpoint-specific survival information. Under these conditions, assumptions of sparsity, linear hazard structure, and proportionality cannot be presumed to hold uniformly across regimes or strata.

Taken together, these properties imply that proportional-hazards feasibility is a structural hypothesis rather than a default modeling assumption in this setting. The combination of high dimensionality, heterogeneous missingness, multi-layer collinearity, and endpoint-specific censoring necessitates an explicit, auditable assessment of whether a given cancer–endpoint stratum admits a stable sparse linear hazard representation.

#### 2. CoxNet as a structural CANARY model

During Phase II execution, Elastic Net–regularized Cox regression was evaluated under fixed identifiability thresholds and a deterministic regularization ladder as an initial feasibility probe of proportional-hazards structure. Across cancer–endpoint strata, this procedure systematically

revealed distinct and interpretable failure modes, including early data infeasibility and exhaustion of the regularization ladder without achieving stable convergence.
These empirical behaviors motivated the interpretation of CoxNet as a CANARY model, whose role is not to serve as a competitive survival predictor, but to diagnostically probe the geometric admissibility of sparse proportional-hazards structure prior to any downstream estimation or inference.

Under this interpretation, CoxNet was evaluated exclusively for structural feasibility—namely convergence, coefficient stability, and the absence of degenerate risk scores. No performance optimization, hyperparameter tuning, or model comparison was performed. Successful feasibility indicates that the stratum admits a sparse, approximately linear hazard representation compatible with proportional-hazards assumptions under auditable constraints. Conversely, systematic failure modes—including regularization exhaustion, instability across the μ ladder, or insufficient event information—were interpreted as evidence that the underlying survival signal exhibits dense, nonlinear, time-varying, or information-deficient geometry incompatible with sparse PH modeling.

Importantly, these outcomes were treated as intrinsic diagnostic properties of the data rather than as modeling deficiencies. The CANARY stage therefore functions as an explicit eligibility gate, determining whether downstream analysis proceeds under proportional-hazards models, alternative non-PH frameworks, or purely descriptive survival summaries.

#### 3. Why CoxNet was not pursued as the final modeling framework

The absence of sparse proportional-hazards feasibility across most strata should not be interpreted as modeling failure. CoxNet was intentionally deployed as a structural probe rather than as a final predictive framework. Elastic Net–regularized Cox regression imposes strong geometric constraints—namely sparsity and approximate linear hazard structure—that serve as a rigorous diagnostic of proportional-hazards admissibility under auditable regularization. The predominance of μ-ladder exhaustion indicates that, while survival information was present in most strata, its structure was incompatible with sparse linear PH assumptions within the deterministic constraint regime. This finding motivated the use of alternative modeling families in downstream analyses, including non-proportional and nonlinear survival frameworks, rather than further relaxation or optimization of CoxNet. The CANARY stage therefore functioned as a principled geometry filter, not as a model competition st

#### 4. Phase I–III Constitutional Contracts

##### 4.1. Contract 1 — Strictly groupwise analysis by cancer type with cancer-prefix predictors

All operations must be executed strictly within cancer-specific strata. Predictor variables are selected locally using a deterministic naming rule that matches the cancer-type prefix embedded in predictor names. No global predictor universe is constructed, no cross-cancer predictor intersection is computed, and no predictor set is shared across cancers. Each cancer type therefore defines its own admissible predictor namespace, potentially spanning multiple omic layers.

**Audit requirement:** For each execution unit, record the cancer-type prefix rule used, the number of predictors selected, and the exact predictor list hash.

##### 4.2. Contract 2 — Strictly endpoint-scoped cohorts (OS, DSS, DFI, PFI) with no global outcomes

All cohort construction and modeling must be performed separately for each survival endpoint. The four endpoints (OS, DSS, DFI, PFI) define independent cohorts, and no global completeness constraint across endpoints is permitted. Missingness in one endpoint must never exclude a sample from analysis of another endpoint. No rule is allowed to "borrow" survival information, thresholds, or decisions across endpoints.

**Audit requirement:** For each execution unit, record cohort size and event count for that endpoint only, and prohibit intersection checks across endpoints.

##### 4.3. Contract 3 — No sample reduction driven by predictors

Within each execution unit, sample identity is preserved. Cohort masking is permitted **only** on the basis of endpoint survival feasibility (time/event present and schema-valid). Predictor missingness must never trigger row deletion. Complete-case filtering is prohibited. If a modeling algorithm requires a finite matrix, resolution must occur via feature-level action and/or deterministic, outcome-independent imputation, but never by excluding samples due to predictor missingness.

**Audit requirement:** For each execution unit, record (i) number of rows before survival masking, (ii) number of rows after survival masking, and (iii) confirm that no subsequent predictor-driven row deletion occurred.

##### 4.4. Contract 4 — Predictor exclusion must be explicit, local, and reason-coded

Any predictor exclusion must be executed only within the local execution unit and must be accompanied by an explicit, reason-coded audit record. Predictor removal is permitted only for numerically or structurally admissible reasons (e.g., zero variance, non-numeric type where numeric required, all-missing, infinite values, or exceeding a fixed missingness threshold if applicable). Predictor exclusion must never be justified by survival outcome behavior, model performance, or cross-unit considerations.

**Audit requirement:** Maintain a per-unit "predictor exclusion ledger" that records predictor name, exclusion reason code, and pre/post predictor counts; include a stable hash of the final predictor set.

#### 5. Phase II Addendum — Structural Identifiability Thresholds (*Emin* = 20, *Nmin* = 50)

To guarantee an absolute barrier against degenerate survival geometry and model hallucination, all strata were subjected to strict minimum statistical identifiability constraints during the Phase II CANARY diagnostic. Cohorts were deemed strictly data-infeasible if they possessed fewer than 20 valid survival events (*Emin* < 20) or fewer than 50 total viable samples (*Nmin* < 50). These exact thresholds were selected based on the architectural prerequisites of the downstream Phase III topology. The *N*min ≥ 50 boundary constitutes the absolute mathematical floor necessary to secure stable bootstrapping and meaningful Out-Of-Bag (OOB) variance estimations within tree-based ensembles (`Random Survival Forests`, `XGBoost`). Concurrently, statistical power in survival models is driven exclusively by observed spatial events, not total sample dimension. The *E*min ≥ 20 boundary rigorously prevents the calculation of "flat" or degenerate Concordance Indices. In a traditional k-fold cross-validation schema, 20 total events guarantee a sufficient non-zero event distribution per validation fold, ensuring the mathematical gradient of calculated Time-Dependent AUCs is driven by genuine clinical mortality and progression rather than artifactual censoring noise.

#### 6. Phase III Addendum — Execution Unit Integrity

For Phase III specifically, all model fitting and prediction must occur within the execution unit:
(cancer type c, endpoint m, dataset variant d, algorithm a)(cancer\ type\ c,\ endpoint\ m,\ data set\ variant\ d,\ algorithm\ a)(cancer type c, endpoint m, dataset variant d, algorithm a)
No decision (feature selection, repair action, seed, parameter choice, or eligibility status) is permitted to operate outside this unit, and no Phase III model output is permitted to retroactively alter TAR admissibility or Phase II feasibility classifications.
**Audit requirement:** Log the full execution-unit key, deterministic seed, and output paths; prohibit any write-back to upstream artifacts.

#### 7. *RSF* implementation note

*RSF* can tolerate missing predictors, which helps Contract 3, but it does not exempt us from Contract 4: if we ever drop predictors (e.g., illegal types, infinite values), we must reason-code every exclusion. Likewise, Contract 1 requires that the *RSF* predictor set is defined by cancer-prefix matching, not by global feature availability.

#### 8. Universal Resume Engine

The Universal Resume Engine for continuous omic variable imputation was implemented in R, providing a robust, parallelized, and fault-tolerant pipeline for high-throughput missing data imputation in multi-omic datasets. The engine supports 12 distinct imputation methods, ranging from simple statistical approaches (mean, median, random sampling) to machine learning-based techniques (k-nearest neighbors, *missForest*, *XGBoost*, *LightGBM*, *MICE*) and advanced matrix completion algorithms (*iSVD*). Each method was applied in a groupwise manner, ensuring that imputation was performed independently within predefined cancer-type subgroups to prevent data leakage.

The pipeline was designed with strict adherence to 16 operational audit criteria, ensuring reproducibility, recoverability, and memory safety. Key features included automated detection of the last valid checkpoint file (.rds) for resuming interrupted runs, dynamic thread allocation based on available system RAM (2–12 CPU cores), and comprehensive logging of all imputation steps. Parallel execution was implemented using the foreach and doParallel packages, with explicit memory cleanup (*rm()*, *gc()*) enforced after each iteration to prevent resource exhaustion.

For methods requiring multiple input features (e.g., *kNN*, *missForest*, *MICE*), a deterministic dummy variable was injected when only a single omic feature was available within a cancer-type subgroup. This ensured compatibility with dimensionality requirements while maintaining biological plausibility. The dummy variable was automatically removed post-imputation, and its usage was logged for downstream auditing.

Diagnostic profiling of missing data burden was performed prior to imputation, stratifying variables by cancer type and omic layer (e.g., mRNA, miRNA, methylation). Variables were then imputed in order of increasing missingness to prioritize high-quality features. Execution details, including the imputation method used, variable counts, and timestamps, were recorded in a centralized audit log (*output_name_table_all.tsv*).

The *iSVD* method was executed via the softImpute package, employing low-rank matrix approximation with iterative refinement.

Error handling was enforced throughout the pipeline using tryCatch blocks, ensuring that failures in individual imputation tasks did not halt overall execution. Corrupted intermediate files were automatically detected and regenerated, while recursive self-invocation logic allowed the pipeline to resume iteratively until completion. The implementation was validated against all predefined operational standards, confirming full compliance with reproducibility, memory safety, and groupwise imputation constraints.

All code, along with detailed documentation, audit logs, and diagnostic reports, was archived to support reproducibility. The pipeline was designed for deployment in large-scale batch processing environments, with modular components allowing for future extensions or modifications.

##### 9. Fault tolerance, auditability, and execution guarantees

All imputation procedures in Phase I were executed using a fault-tolerant, audit-oriented execution engine designed for large-scale, groupwise multi-omic processing. The engine enforces cancer-type isolation, deterministic checkpointing, and full provenance tracking for every generated preprocessing variant. Intermediate outputs are materialized as indexed RDS objects and registered in a centralized audit table, enabling exact recovery, reproducibility, and downstream traceability. Execution failures are isolated at the variable or method level and do not interrupt global pipeline completion. This design ensures that all preprocessing variants entering Phase II are fully auditable, reproducible, and free of cross-cohort contamination.

#### Supplementary Note 3: Operational Architecture of the Multi-View Meta-Learning ($\mathcal{MVL}$) SuperLearner

##### 1. Distinction between the Quadripartite Base-Learner Ensemble and the $\mathcal{MVL}$ SuperLearner

The Phase III predictive framework comprises two structurally distinct and hierarchically nested computational layers, whose separation is fundamental to both the mathematical validity and the interpretability of the resulting survival predictions.

The Quadripartite Base-Learner Ensemble constitutes the first layer. It consists of four independent, non-linear, non-proportional survival algorithms — Random Survival Forest ($RSF$),

Extreme Gradient Boosting (*XGBoost*), Multi-Task Logistic Regression (*MTLR*), and Survival-*Boruta* Independent — each derived in strict isolation on the full multi-omic predictor matrix for a given cancer-endpoint-preprocessing cohort (c, m, d). Critically, each algorithm operates on the same raw omic feature space but applies fundamentally different inductive biases, dimensionality strategies, and splitting or optimization topologies. Consequently, each base learner produces a distinct, algorithm-specific summary of the underlying multi-omic survival landscape, encoded as a single continuous mortality risk score per patient.

The Multi-View Meta-Learner (*MVL*) SuperLearner constitutes the second, superior layer. Rather than receiving the raw omic feature matrix as input, the *MVL* receives an entirely new — and dramatically lower-dimensional — matrix of dimension N × 4, where each column corresponds to the patient-level risk score generated by one of the four base learners. This meta-matrix is standardized prior to submission. The *MVL* then fits an Elastic Net-regularized Cox proportional hazards model (α = 0.5; λ selected by cross-validated penalization) on this meta-matrix, learning the optimal survival-weighted linear combination of the four base-learner risk scores. The resulting super-risk score is the final individualized survival prediction of the SuperLearner architecture.

This two-layer nesting is illustrated schematically below:

```
RAW OMIC MATRIX (N × P, where P = thousands of multi-omic features)
       │
       ├──→ RSF           → OOB mortality risk          [N × 1]
       ├──→ XGBoost       → Apparent log-hazard risk    [N × 1]
       ├──→ MTLR          → Cumulative apparent risk    [N × 1]
       └──→ Boruta-RSF    → OOB mortality risk          [N × 1]
                            │
                            ▼

              META-MATRIX (N × 4, standardized)

              ┌────────────┬───────────┬──────────┬────────────┐
              │    RSF     │ XGBoost   │   MTLR   │ Boruta-RSF │
              └────────────┴───────────┴──────────┴────────────┘
                            │
                            ▼

           Elastic Net Cox Meta-Model (α = 0.5, CV-λ)

           SuperRisk =
           β1·RSF + β2·XGBoost + β3·MTLR + β4·Boruta-RSF
                            │
                            ▼

                SUPER-RISK SCORE [N × 1]

                → C-Index
                → AUC(t) at 1, 3, and 5 years
```

#### 2. What Each Layer Knows

The five computational components of the SuperLearner architecture receive fundamentally different inputs, apply distinct analytical logics, and produce incommensurable outputs. The table below summarizes the input space, analytical scope, and output of each layer:

| Layer | What it sees | What it learns | Output |
|---|---|---|---|
| *RSF* | All omic features (full X matrix) | Which features best split survival trees | OOB mortality risk score + VIMP |
| *XGBoost* | All omic features (full X matrix) | Which features drive the log-hazard | Apparent log-hazard risk score + *SHAP* + *LIME* + Gain |
| *MTLR* | *Boruta*-selected features only | Which features predict survival time intervals | Apparent cumulative risk score (no post-hoc feature attribution) |
| Survival-*Boruta* | All omic features (full X matrix) | Which features are statistically confirmed relevant | OOB risk score (on confirmed features) + attStats decisions |
| *MVL* SuperLearner | 4 risk scores only (N × 4 meta-matrix) | Which algorithm is most trustworthy per cohort | Fused super-risk score + β weights |

*Note on interpretability scope: Post-hoc interpretability tools (SHAP, LIME) are applied exclusively to XGBoost due to its native compatibility with exact TreeSHAP computation and the LIME framework. RSF and Survival-Boruta carry their own native importance metrics (VIMP and attStats, respectively). MTLR has no dedicated post-hoc interpretability layer — its feature dimensionality is constrained upstream by the Boruta gate.*

The *MVL* SuperLearner operates exclusively in algorithm-prediction space, learning optimal survival-weighted combinations of base-learner risk estimates rather than individual omic features. Consequently, post-hoc interpretability of omic-level predictive drivers was performed independently at the base-learner level via *TreeSHAP* (*XGBoost*), out-of-bag Variable Importance (VIMP, *RSF*), and *Boruta* feature confirmation — each capturing a distinct, algorithm-specific view of the multi-omic feature landscape driving patient survival.

#### 3. What the *MVL* SuperLearner Learns — and What It Does Not

The *MVL* SuperLearner operates exclusively in algorithm-prediction space, not in omic-feature space. Its β coefficients do not describe the importance of any omic signature. Instead, they represent the relative predictive trustworthiness of each base learner for a given cohort —

specifically, to what degree each algorithm's implicit encoding of the multi-omic survival landscape should be weighted in the final fused prediction.

This distinction has a critical interpretive consequence: the *MVL* SuperLearner cannot and does not identify which omic features drive patient survival. Omic-level feature importance is determined exclusively at the base-learner layer, through algorithm-specific tools applied strictly to their respective models:

| Interpretability Tool | Base Learner | What It Reveals |
|---|---|---|
| *TreeSHAP* Beeswarm & Dependence Plots | *XGBoost* only | Global and local omic feature impact on log-hazard |
| *SHAP* Waterfall (Patient-Level) | *XGBoost* only | Per-patient omic feature contribution to individual risk |
| *LIME* Local Attributions | *XGBoost* only | Linearized neighborhood approximation of individual predictions |
| Out-of-Bag VIMP | *RSF* only | Variable importance for survival tree splitting |
| *Boruta* Feature Decisions | Survival-*Boruta* only | Statistical confirmation of omic features as globally relevant |
| (No post-hoc feature attribution) | *MTLR* | Feature selection delegated upstream to *Boruta* gate |

The *MVL* operates as a synthesis layer, not an interpretability layer. It answers the question "which combination of algorithm views optimally predicts survival?", while *SHAP*, *LIME*, VIMP, and *Boruta* answer the question "which omic features actually drive it?"

#### 4. Why the *MVL* Uses Elastic Net and Not LASSO

The use of Elastic Net regularization (α = 0.5) in the *MVL* meta-model reflects a deliberate and methodologically consistent architectural decision. During Phase II, Elastic Net-regularized Cox regression (CoxNet) was applied directly to the high-dimensional raw multi-omic feature matrices. The overwhelming prevalence of μ-ladder exhaustion across the pan-cancer matrix (Supplementary Figure S7) demonstrated that linear proportional-hazards geometry was structurally incompatible with the multi-omic survival signal in the vast majority of strata. This finding constituted the formal CANARY rejection of LASSO-class linear models as predictive frameworks for this dataset.

The *MVL* Elastic Net is architecturally non-redundant with this rejection for the following reason: it is not applied to the raw omic feature space — in which LASSO-type models were formally disqualified — but to the low-dimensional (N × 4) space of base-learner predictions produced by four independently validated non-linear survival algorithms. The input dimensionality, the nature of the features, and the geometric properties of the meta-matrix are fundamentally different from those of the Phase II CoxNet inputs. The *MVL* Elastic Net therefore does not violate the Phase II rejection, but operates in a complementary and structurally distinct domain.

The choice of $\alpha = 0.5$ (balanced Ridge–LASSO mixture) is further motivated by the nature of the meta-matrix: with only four input columns, pure LASSO ($\alpha = 1$) risks pathological sparsity by zeroing out one or more base-learner contributions, while pure Ridge ($\alpha = 0$) enforces equal shrinkage regardless of relative predictive contribution. The balanced Elastic Net penalization allows the *MVL* to dynamically suppress informationally redundant base-learner views while preserving complementary predictive signals across non-redundant algorithms.

#### 5. Relationship to the Phase III Constitutional Contracts

The two-layer nesting of the SuperLearner is fully consistent with the Phase III Constitutional Contracts governing cohort isolation (Contract 1) and endpoint-scoped analysis (Contract 2). Both the base-learner derivation and the *MVL* fitting are executed strictly within each cancer-endpoint-preprocessing unit (c, m, d), with no information shared across strata. The *MVL* does not aggregate information across cancer types or endpoints; it synthesizes information exclusively within the cohort-specific risk prediction space.

The *MVL* β-coefficient weights are therefore cohort-specific and are not interpretable as universal algorithm rankings. A base learner that receives high weight in one cancer-endpoint stratum may receive zero weight in another, reflecting genuine structural heterogeneity in the relative predictive capacity of different algorithmic views across the pan-cancer survival landscape.

#### 6. Architectural Fallback Defenses: The "No Cohort Stays Behind" Policy

To ensure robust evaluation across all 96 cancer-endpoint strata without dropping mathematically challenging cohorts, the Phase III framework implements two strictly algebraic, "No One Stays Behind" fallback defenses. Neither defense utilizes Bayesian imputation; rather, they rely on deterministic Z-score properties and random chance baselines to prevent singular matrix crashes.

6.1. The *Boruta* Coerced_0.5 Resolution: If the Survival-*Boruta* algorithm evaluates a highly noisy matrix and returns zero statistically confirmed features, it mathematically cannot map a survival forest. Rather than crashing the master loop, the architecture resolves this by defining a neutral baseline hazard vector (absolute zeros), assigning a structural C-Index of 0.5 (random chance probability), and flagging the execution log with "Coerced_0_Features". This accurately penalizes *Boruta*'s performance metric in that specific cohort while preserving the integrity of the 96-cohort evaluation.

6.2. The *MVL* SuperLearner Missingness & Variance Resolution: Within the Elastic Net SuperLearner (*MVL*), missing base-learner predictions (NAs) or zero-variance predictions (which trigger singular matrix failures) are resolved strictly in standardized space:
6.2.1. Z-Score Mean Imputation: The *MVL* first standardizes the available predictions forming the meta-matrix (scale()). Any NA values resulting from base learner convergence failures are then explicitly replaced with 0. Because the matrix operates in Z-score space, a value of 0 rigorously corresponds to the population mean risk. This algebraically neutral imputation ensures matrix convergence without pulling the meta-prediction toward an artificially high or low risk.
6.2.2. Micro-Jitter Variance Injection: If an algorithm generates a constant risk score for all patients (variance of zero), the *MVL* injects an innocuous microscopic uniform noise (runif between -1e-6 and +1e-6). This noise breaks the mathematical tie necessary for Elastic-Net execution while remaining entirely inert regarding the biological topology and survival ranking.

**7. Sparsity Architecture and Clinical Audit Generation**

In advanced gradient boosting frameworks and tree algorithms, systemic sparsity (e.g., patient data where critical predictive molecular biomarkers are unmeasured) forms distinct routing pathways. Mathematical analyses of the Phase I and II feature spaces identified an ascertainment bias paradigm, wherein complete localized absence of sequenced predictors generated "default-path" algorithmic mapping independent of verifiable biology. By establishing a rigid ≥ 35% geometric exclusion parameter, the SuperLearner architecture computationally excised "ghost" clinical patients whose predictive structures were fundamentally hollow.

During parallel processing sweeps across all Phase III endpoint scenarios, the ensemble algorithm autonomously generated precise execution logs (_Excluded_Patients_Geometric.tsv). This diagnostic tracing recorded the exact biological strings (TCGA Subject IDs) of the excluded participants, validating that 98% of mathematical exclusions presented with exactly 100% missing functional omic profiling. Furthermore, bounded tracking logged all algorithmically retained cases (_Retained_Partial_Missing.tsv) that possessed intermediate sparsity (>0% and <35%). This independent catalog ensures definitive downstream transparency for interpreting the structural logic underpinning post-hoc decision trajectory algorithms (e.g., *TreeSHAP* and *LIME* implementations).

**Supplementary Note 4: Mathematical Framework of the Brier Calibration Audit**

**Native Risk Extraction and Topological Synchronization**

Unlike standard linear predictive models where error is algebraically trivial, evaluating complex survival ensembles post-hoc requires extracting continuous risk proxies and projecting them temporally. To strictly preserve the exact computational state of the Phase III models, risk vectors were natively extracted from the saved model bundles across all five architectural dimensions:

**Random Survival Forest (*RSF*):** Extracted via native Out-Of-Bag (OOB) ensemble mortality predictions (predicted.oob).

**Extreme Gradient Boosting (*XGBoost*):** Extracted via row-wise *SHAP* (*SHAP*ley Additive exPlanations) margin summations to definitively bypass environment-dependent C++ pointer corruption.

**Survival-*Boruta*:** Extracted by dynamically projecting the independent noise-separated topological subset (Confirmed and Tentative features) into an independent *RSF* proxy framework, governed by identical Phase III execution constraints (ntree = 500, logrank splitting).

**Multi-Task Logistic Regression (*MTLR*):** Extracted via native apparent risk indices.

***MVL* Super-Learner**: Reconstructed dynamically using the optimal Elastic-Net tuning dimensions (e.g., Lambda) established during Phase III derivation, projecting a weighted, highly polarized risk continuum derived from the Quadripartite predictions.

A critical challenge in cross-algorithm extraction is implicit internal geometry dropping. Specifically, *RSF* C-backends executing native topological imputation (na.impute) implicitly drop records where Time <= 0, resulting in mathematical length discrepancies between the predicted risk vector and the source clinical dataframe. To preserve exact data provenance to the TCGA patient barcodes, a dynamic geometric mapping bypass (safe_assign_risk) was enforced. This topological lock verified the dimensionality of the algorithm's output, isolated the indices of valid patients (Time > 0), and structurally mapped the truncated prediction vectors back onto the full clinical manifest, padding exclusions with NA to prevent array corruption.

Clinical Endpoint Synchronization: Mortality vs. Progression Probabilities It is critical to delineate the nomenclature and clinical interpretation of the extracted probabilities prior to calibration, as they must accurately reflect the specific clinical endpoint under evaluation.

For models predicting OS and DSS, the underlying Cox Breslow estimators natively derive the Probability of Survival, denoted mathematically as *S(t)*. Reporting *S(t)* is the universal convention in clinical oncology for mortality endpoints, representing the probability that a patient will survive (i.e., not experience the terminal event) beyond a specific temporal landmark *t*.

Conversely, for models evaluating the Disease-Free Interval (DFI) and Progression-Free Interval (PFI), clinical convention defines the hazard as the cumulative risk of recurrence or progression (Probability of the Event). Mathematically, the probability of the event occurring is the exact complement of the survival function: P(Event at time t) = *1 - S(t).*

Because the Brier Score calculates the Mean Squared Error of these probabilities, the mathematical penalty is perfectly symmetric whether evaluating the error of surviving versus the error of progressing. Therefore, the computational pipeline natively processes the strict Probability of Survival, *S(t)*, across all clinical endpoints. This topological symmetry guarantees that the Brier Score identically captures the true predictive error against the correct clinical context, whether measuring a patient's survival chance (OS/DSS) or their inverse risk of relapse (PFI/DFI).

**Inverse Probability of Censoring Weighting (IPCW) Calibration**

The absolute accuracy of the probability outputs was evaluated using the Brier Score (Mean Squared Error for probabilities). For a specific time point t (e.g., 365, 1095, or 1825 days), the Brier Score measures the algorithm's predicted survival probability against the true binary clinical status. To account for patients whose follow-up ended before time t without an event (right-censoring), the predictions were weighted using the IPCW estimator $G(t)$:

$$BS(t) = (1 / N) * SUM \left[ (Y_i(t) * (0 - p_i(t))^2 ) / G(T_i) + ( (1 - Y_i(t)) * (1 - p_i(t))^2 ) / G(t) \right]$$

Where: N is the total cohort size. $p_i(t)$ is the algorithm's predicted probability of survival, S(t), identically evaluated across all endpoints. $Y_i(t)$ is an indicator of the known clinical status of patient i at time t. G(t) represents the Kaplan-Meier estimate of the censoring distribution at time t. $G(T_i)$ represents the Kaplan-Meier estimate of the censoring distribution at the patient's observed event time.

Continuous Temporal Profiling (Prediction Error Curves) To evaluate algorithmic decay over time, this exact IPCW Brier Score calculation was iteratively extended across 50 dense temporal slices spanning the entire 5-year clinical follow-up period. This dynamic mapping allowed for the generation of continuous Prediction Error Curves. Crucially, this temporal profiling was applied uniformly across all four non-linear base-learners (*RSF*, *XGBoost*, Survival-*Boruta*, *MTLR*) and the overarching *MVL* Super-Learner. The area under these curves was subsequently calculated via trapezoidal integration to compute the final Integrated Brier Score (IBS), representing the absolute cumulative predictive error for each specific algorithm.

**Super-Learner Polarization**

The resulting patient-level survival probability trajectories ($p_i(t)$) computationally highlighted a defining mathematical property of the *MVL* Synthesis approach. While base-learners like *XGBoost* produced smooth, conservative probability decay curves as time advanced, the *MVL* Elastic-Net Super-Learner exhibited distinct mathematical polarization. By aggregating the risk vectors of the Quadripartite ML Ensemble, the *MVL* acts as an amplifying judge: when the base models agree on a high-risk topography, the *MVL* aggressively drives the survival probability toward 0%; when the models mathematically agree on biological stability, the probability is locked near 100%. This polarization mathematically validates the Super-Learner's structural utility in definitively separating clinical outcome trajectories over the 5-year landmark period.

**Supplementary Note 5: Surrogate Mapping and Geometric Fidelity in *LIME* Individualized Risk Attributions**

**1. Localized Surrogate Geometries and Point-of-Care Explanations**

While *SHAP* provides exact marginal contributions mapped across the entire cohort topology, Local Interpretable Model-agnostic Explanations (*LIME*) is deployed exclusively to dissect "point-of-care" individualized hazard boundaries. Rather than attempting to decompile the global architecture of the nonlinear SuperLearner matrix, `LIME` mathematically isolates a microscopic geometric "neighborhood" surrounding a single patient's clinical coordinate. Within this highly restricted boundary, `LIME` constructs a sparse, penalized linear surrogate model to approximate the localized decision logic of the underlying complex algorithm.

The resulting visualization plots the exact linear regression coefficients of this localized model (plotted on the X-axis as Weight). These discrete weights explicitly map how the biological presence or abundance of a targeted multi-omic signature shifts the patient's localized prediction relative to the localized algorithmic baseline.

- **Supports (Blue Bars):** Features classified as "Supporting" mathematically elevate the predictive output of the algorithm. Operating natively in the survival domain, these omic signatures actively accelerate the individualized log-hazard ratio, driving the patient strictly toward increased relative mortality risk.
- **Contradicts (Red Bars):** Features classified as "Contradicting" depress the predictive output. These act as localized protective factors, mathematically pulling the patient's individual trajectory down and mitigating the primary survival hazard.

**2. Explanation Fit ($R^2$) as a Diagnostic Proof of Severe Non-Linearity**

The `LIME` algorithm accompanies each individualized model with an Explanation Fit metric, representing the Goodness of Fit (R2) of the localized linear surrogate against the absolute predictions generated by the underlying `XGBoost` survival network.

Crucially, within the highly interactive multi-omic topographies of the 96 evaluated cohorts, localized Explanation Fits frequently yielded exceptionally low boundaries (median $R^2 = 0.029$, with 91.2% of patients falling below $R^2 < 0.10$). Rather than signifying a failure of the analytical execution, these suppressed fit metrics function as a profound structural diagnostic. They provide immediate, mathematical proof that `LIME`'s linearized surrogate models simply cannot handle the extreme non-linearity of complex multi-omic spaces. Consequently, this geometric limitation absolutely validates our decision to forcefully anchor the primary interpretability of the workflow to the topologically exact, non-linear `TreeSHAP` architecture.

If an individual patient's localized survival space was governed by additive, independent biological factors, the sparse `LIME` linear surrogate would easily approximate the boundary and yield a high Explanation Fit. However, low fit values unequivocally demonstrate that even when zooming into the narrowest geometric coordinate of a single patient, the localized survival boundary remains so violently nonlinear, synergistic, and interactive that a linear proxy structurally shatters. Consequently, this metric mathematically validates the clinical indispensability of the Quadripartite ML Ensemble; it explicitly proves that traditional, strictly parametric linear survival frameworks (e.g., standard Cox Proportional Hazards models) are structurally incapable of navigating these complex, interactive omic topologies.

**Supplementary Note 6: Mathematical Formulation of the Internal Blind Validation Inference Engine Geometrical Preservation and Zero-Information Penalties**

To simulate a true prospective diagnostic environment without data leakage, the internal validation matrix (comprising N = 1,050 pristine patient vectors; Supplementary Dataset S3) was mathematically forced into absolute geometrical parity with the Phase III derivation matrix (Supplementary Dataset S2). To establish the final pristine cohort, the extracted validation matrix was subjected to a strict biological exclusion filter: any patient vector that exhibited 100% missingness (NA) across its entire cancer-specific multi-omic signature was mathematically purged from the cohort, ensuring that only biologically viable patient records advanced to the inference engine (e.g., resulting in the total exclusion of the severely fragmented UVM and DLBC cohorts).

To prevent the artificial escalation of risk scores caused by remaining topological missingness, deterministic algorithms were protected by a mathematical baseline fallback. Specifically, for algorithms requiring complete matrices, missing variables were anchored using Phase III topological imputation matrices. Conversely, for sparsity-aware algorithms such as *XGBoost*, feature absence exerted a strictly neutral, zero-information penalty. If a validation vector $x_i$ contained a missing variable $x_{ij}$, the sparsity-aware split-finding algorithm mathematically routed the observation down the optimal default branch $h(x_i)$ established during Phase III derivation, thereby minimizing the Cox partial likelihood loss without hallucinating synthetic values. The exact multi-omic dimensionality and the outcome of the biological exclusion filter are detailed in Supplementary Table S5.

**Quadripartite Risk Generation and Continuous Hazard Extraction**

For structurally intact patient signatures, the validation engine calculated continuous hazard vectors across four distinct algorithmic topologies:

*Random Survival Forests* (*RSF*): Aggregated cumulative hazard functions $H(t|x_i)$ derived from an ensemble of decision trees utilizing log-rank splitting rules to maximize survival separation at internal nodes.

*MTLR* (Multi-Task Logistic Regression): Modeled the survival curve by discretizing the time domain into discrete intervals, utilizing a penalized maximum likelihood function to estimate the probability of the event occurring within each interval.

*Boruta*: Generated continuous risk signatures derived from topologically validated, dimensionality-reduced feature sets.

*XGBoost*: Extracted gradient-boosted hazard predictions by iteratively minimizing the negative log partial likelihood of the Cox proportional hazards model, L(beta).

**Elastic Net Synthesis and Dual-Track Safe Masking (Path A)**

The Phase III SuperLearner architecture synthesized the four independent hazard vectors into a singular, continuous risk Z-score, $Z_{SL}$, via a rigid linear combination:

$$Z_{SL} = \sum [ \beta_k \cdot R_k(x_i) ] \text{ for } k \in \{RSF, XGBoost, MTLR, Boruta\}$$

where $R_k(x_i)$ represents the continuous risk score from algorithm k, and $\beta_k$ represents the exact penalized elastic net coefficient frozen during Phase III derivation.

However, when the inference engine encountered highly fragmented validation records, the *Boruta* topological imputation layer mathematically failed to converge. Because *Boruta* was a strictly required dimension ($\beta_{Boruta}$ != 0), the geometric matrix collapsed. The architecture autonomously recognized the missing scalar $R_{Boruta}$ and safely aborted the linear combination, intentionally returning Z_SL = NA. This strict gating mechanism prevented the synthesis of hallucinated linear combinations. The frequency of these algorithmic safety abortions is mapped in Supplementary Table S6.

**Fallback Extraction via Sparsity-Aware Routing (Path B)**

To guarantee clinical predictive coverage without patient attrition, the Dual-Track architecture implemented a programmatic fallback (Path B). When Z_SL -> NA, the pipeline isolated the pure `XGBoost` risk component, R_XGB(x_i). Because the native `XGBoost` module handles missingness via mathematical default-branch routing rather than linear imputation, it successfully generated valid continuous hazards for the fragmented records. As verified in Supplementary Table S7, this architecture achieved 100% computational penetrance, ensuring every sequestered patient received an algorithmically derived risk score.

**Baseline Probability Extraction and Topological Convergence Limits**

While the Dual-Track architecture successfully achieved 100% predictive penetrance in generating continuous hazard Z-scores across all sequestered patients, the subsequent translation of these scores into absolute clinical probabilities exposed the structural limits of highly sparse topological strata. Specifically, when anchoring the validation hazard vectors against the Phase III Breslow baseline survival estimator (basehaz()), the inference engine encountered localized topological crashes—manifesting as infinite partial likelihood coefficients—exclusively within severely fragmented validation cohorts (e.g., specific LGG endpoints). These localized mathematical singularities necessitated explicit UI masking during the final clinical probability deployment to prevent graphical hallucination. The exact frequencies and cohort distributions of these infinite-coefficient convergence failures are formally audited and quantified in the Baseline Convergence Audit (Supplementary Table S21).

#### Supplementary Note 7: Mathematical Validation of Algorithmic Displacement and Local Explainer Nullification

**Overview:** The Sparsity Isolation Protocol specifically investigated the isolated predictive limits of baseline ML algorithms when forcefully restricted to sparse, non-continuous feature vectors geometries (Somatic Mutations and CNVs). The raw performance outputs detailed in Supplementary Table S10 exhibit severe, systemic algorithmic breakdowns across independent `RSF` and `XGBoost`.

**The Mechanism of Displacement**: The most critical validation of this spatial degradation mechanism occurred organically during the automated explainability extraction protocols via `SHAP` and `LIME`. When evaluated on the primary "Supreme" sparsity exemplar (ESCA-DFI-df017), the `XGBoost` model mathematically surrendered at an absolute C-Index of 0.500, indicating mathematical equilibrium and flatlined risk stratifications. Because the underlying C++ logic generated zero substantive tree splits on the binary fields, the algorithm yielded identical risk constants transversally across the cohort.

Consequently, when directed to compute localized feature-attribution variance, both `SHAP` and `LIME` native engines safely triggered singularity bypass protocols. Because no multi-dimensional spatial gradients or mathematical decision splits physically existed in the localized survival surrogate model, it was computationally impossible to draw interactive dependence vectors. The organic nullification of these specific visual artifacts physically validates algorithmic displacement: highlighting that localized feature explanations require, as an absolute prerequisite, the continuous variance and topographical depth inherently provided by full spectrum multi-omic integration.

**Supplementary Note 8: Phase III Mathematical Rejection Audit (The 1,982 Null Cohort)**
To empirically prove why the 1,982 multi-omic signatures were ejected natively by the Phase III machine learning pipelines (as comprehensively cataloged in Supplementary Table S16), we benchmarked their structural topology focusing strictly on non-linear entropy constraints (*Random Survival Forests* and *XGBoost* paradigms), directly analyzing them against the 12,613 variables that successfully passed algorithmic extraction.

Our zero-leakage pipeline dictates that features are only retained if they exhibit autonomous, non-random predictive entropy (Information Gain) strictly inside isolated cancer subgroups. When interrogating the 1,982 rejected variables within their locked etiology matrices, two lethal non-parametric thresholds were consistently violated:

1. The Localized Information Gain Deficit In gradient boosting (*XGBoost*) and *RSF* architectures, algorithms rely on Gini Impurity (or Survival Variance) to execute decision-tree bifurcations. Mathematically, a feature is only selected for a node split if it provides a strictly positive Information Gain ($\Delta G > 0$). During group-wise isolation (e.g., testing a rejected Transcriptomic feature strictly within isolated Glioblastoma OS = GBM), over 98% of the localized permutations for these 1,982 features resulted in $\Delta G \leq 0$ or Negative VIMP. This definitively proves that within their native cellular environment, these variables exhibited stochastic, multidimensional noise. They fundamentally failed to reduce the geometric uncertainty of patient mortality, rendering them mechanically invisible to the predictive algorithms.

2. Native Intra-Cohort Sparsity (Zero-Variance Splitting Limits) Predictive AI algorithms actively assess base variance prior to execution to prevent infinite mathematical recursion. In *XGBoost*, parameters such as min_child_weight define the absolute minimum sample variance required to authorize a topological branch. Upon localized benchmarking, the 1,982 rejected features routinely victimized themselves via extreme variance starvation. Because their variation (especially rare mutation vectors or sparse transcripts) was heavily clustered globally, partitioning the dataset structurally isolated these features and drove their intra-cohort variance to $\sigma \approx 0.00$. Without localized numerical deviation natively present to feed the split threshold, the $\Delta G$ equation structurally collapses into an undefined state (attempting to split identical values). Because the decision trees mathematically cannot bifurcate the uniform patient plane, the features possessed zero predictive entropy, ensuring immediate programmatic ejection.

**Supplementary Note 9: Interpretability Mechanics of *TreeSHAP* Decision and Waterfall Topographies**
**Algorithmic Aggregation of Long-Tail Omic Features**

Within the Phase III Quadripartite ML Ensemble, point-of-care interpretability is resolved via `TreeSHAP` decision and waterfall visualizations. These plots map the exact localized multi-omic variables responsible for shifting an individual patient's hazard ratio from the population baseline (Ef(x)) to their final specific prediction (*f(x)*).

Due to the extreme high-dimensionality of the underlying LiSHMOM matrices, plotting the individualized exact marginal contributions of all evaluated features simultaneously generates severe visual cognitive load and obscures dominant therapeutic targets. To resolve this, a mathematical plotting constraint is enforced: the visualizations explicitly isolate and rank only the absolute most dominant apex drivers (e.g., the top 10 most impactful features) contributing to the individualized survival outcome.

All remaining features are algorithmically grouped into an aggregate "*X other features*" category (e.g., "*90 other features*"). Crucially, this grouped label is not a statistical approximation; it represents the precise arithmetical sum of the localized `SHAP` values for the entire remaining long-tail subset. This strictly preserves the mathematical integrity of the localized hazard calculation while shielding the overarching clinical signal from the biological noise of hundreds of highly correlated, low-impact variables.

Dynamic Y-Axis Hierarchical Ranking The visualizations enforce a dynamic hierarchical ranking on the Y-axis, ordering the displayed features—including the aggregated "X other features" bucket—strictly by the absolute magnitude of their localized `SHAP` contribution for that specific patient. Consequently, the position of the aggregated "other features" bucket fluctuates deterministically from patient to patient, operating as a distinct biological indicator:

- **Subservient Rank (e.g., 3rd or lower):** When the "other features" category appears below specific individual drivers, the patient's survival topology is governed by profound, isolated **apex aberrations** (e.g., an extreme, localized epigenetic hypermethylation event or targeted somatic mutation). In these coordinates, the sheer magnitude of the primary multi-omic aberration mathematically dominates the cumulative background noise of the long-tail network.
- **Apex Rank (Highest Position):** Conversely, if the cumulative sum of the "other features" ranks at the very top of the hierarchy, it signifies a survival topology driven by massive, diffuse polygenic degradation—often characterized as a "death by a thousand cuts." In these localized terrains, no single therapeutic target dominates; rather, dozens of minor, distributed omic shifts interact synergistically to generate a massive cumulative hazard penalty that outweighs any individual driver.

Ultimately, this dynamic aggregation mechanism ensures that the visualizations accurately identify whether an individual patient's risk is functionally driven by targetable apex nodes or determined by a vast, distributed regulatory cascade.

**Supplementary Note 10: Non-Linear Synergy and Algorithmic Rescue**

The biological reality of these mathematical topologies is profoundly demonstrated within the LGG architecture. The automated extraction of complex signatures, such as LGG-972.6.3.P.3.69.71.2.4.2, does not represent a singular prognostic marker, but rather the concordant mRNA layer amalgamation of *FBXL20, HSF2, NF1, PAX5, PSMD10, and RBPJ*. According to the foundational database variables (Supplementary Table S8), this specific algorithmic node mathematically embodies the Overexpression of these genes and a direct, positive correlation (P) with the TSM phenotype. Biologically, this implies that the expression of this signature drives an increased stemness potential (ergo, dedifferentiation) within the tumor. However, by capturing the intense crosstalk strictly between the Apoptosis and Autophagy pathways, this multi-element amalgamation acts collectively as a highly Protective overall clinical outcome. Crucially, this is precisely where the SuperLearner's rigorous validation demonstrates its value: despite its strong bivariate protective association at the population level, the quad-algorithmic ensemble assigned this specific signature a strict importance score of zero (Supplementary Table S8), effectively rejecting it as a structurally redundant variable when analyzed within the full multivariate landscape.

Similarly, the isolation of the epigenetic signature LGG-1202.4.3.N.0.35.35.1.2.4 directly maps the concordant prognostic association of hsa-miR-148a-3p and hsa-miR-148a-5p within the TSM phenotype. In stark contrast to the mRNA signature, this topology exhibits a negative correlation (N) with TSM, implying its expression is linked to decreased stemness potential. Furthermore, this topology spans an extraordinarily broad RCD spectrum, regulating the critical threshold across four simultaneous modalities: Apoptosis, Autophagy, Necroptosis, and Necrosis. Ultimately, the multi-omic architecture identifies this negatively correlated, multi-element microRNA node as a fundamentally Risky prognostic indicator. In this case, the algorithmic audit explicitly confirmed the pre-existing population-level finding, with the SuperLearner categorizing it as a mathematically critical predictor of lethal trajectories and officially validating it via non-linear *XGBoost* extraction (Gain = 0.0059).

Crucially, the relationship between these two specific signatures perfectly illustrates the fundamental necessity of non-linear machine learning. As established, the SuperLearner assigned the protective mRNA amalgamation (LGG-972.6.3.P.3.69.71.2.4.2) a main-effect importance of zero, while independently validating the risky microRNA amalgamation (LGG-1202.4.3.N.0.35.35.1.2.4) via *XGBoost*. A classical linear model would simply discard the mRNA signature as statistically useless. However, the subsequent Tree*SHAP* topological analysis revealed a profound mathematical reality: these two signatures actively interact. Despite lacking an independent main effect, the unvalidated mRNA amalgamation acts as a potent non-linear modulator, combining with the validated microRNA signature to trigger a statistically undeniable SYNERGY (Hazard Amplification). Classical additive proportional-hazards models frequently average these disparate, intersecting effects toward zero, resulting in prognostic failure. The Quadripartite Ensemble bypasses this limitation by treating these RCD pathways not as isolated variables, but as colliding survival topologies. The synergistic hazard amplifications and antagonistic prognostic protections mapped by Tree*SHAP* are the direct mathematical representations of these physical RCD modalities interacting in vivo. Ultimately, by reverse-engineering these physiological circuits from raw multi-omic mass, the SuperLearner establishes that precision oncology cannot rely on isolated biomarkers; it requires the deterministic mapping of the entire non-linear RCD landscape.

**Supplementary Note 11. Algorithmic Architecture and Operational Inference of the CancerRCDPredictor Application**

**Mathematical and Structural Framework of the Patient Trajectory Engine**

The CancerRCDPredictor application (https://lbtgenomica.uenf.br/cancerrcdpredictor/), deployed via the R/Shiny framework, functions as the translational interface for the Pan-Cancer Multi-Omic SuperLearner. The structural dimensionality of the application relies on the deterministic integration of the pre-computed Quadripartite Base-Learner Ensemble (*Random Survival Forests*, *XGBoost*, insulated Survival-*Boruta*, and Multi-Task Logistic Regression) and the overarching Elastic Net Multi-View Meta-Learner (*MVL*). The analytical engine isolates individual patient configurations, computing real-time, time-dependent survival estimates at 1-year, 3-year, and 5-year horizons natively derived from the *MVL*. Crucially, the underlying metric intelligence core explicitly bifurcates mathematical scaling: global survival metrics (OS and DSS) compute probability decay mathematically mapping to a deleterious trajectory, whereas event-based recurrence metrics (DFI and PFI) compute cumulative incidence progression, wherein escalating probabilities map to lethal recurrence risk.

**High-Dimensional Topological Navigation and Personalized *SHAP* Decoding**

To achieve mathematically grounded Explainable AI (XAI), the application bypasses universal feature importance rankings in favor of localized, patient-specific topological deconstruction. This operation extracts post-hoc *SHAP*ley Additive exPlanations (Tree*SHAP*) geometry exclusively from the *XGBoost* sub-model array. The decoding algorithm isolates the Top 5 unique multi-omic coordinates dominating the individual's specific survival topology.

To bridge this abstract geometry with biological mechanism, the pipeline employs a deterministic string-matching protocol. Each extracted alphanumeric signature (e.g., LGG-2641.7.3.P.3.44.39.1.2.1) is queried against the master target repository. This programmatic parsing dynamically retrieves: (1) the decoded genetic elements and their literature-established functional definitions; (2) the exact biological omic layer encoding the risk (e.g., transcript isoforms vs. CpG methylation); (3) validated Boolean relationships to 25 distinct Regulated Cell Death (RCD) pathways; and (4) population-level Spearman correlation arrays mapping the topological coordinate to generalized tumor phenotypes (TMB, MSI, and TSM).

**Algorithmic Payload Assembly and Localized LLM Inference**

To synthesize these high-dimensional parameters into clinical intelligence without compromising auditability or violating patient data privacy, the platform integrates a strictly guarded Large Language Model (LLM) framework. The architectural payload relies on the httr2 package in R to negotiate asynchronous API requests directed to a completely localized, offline inference engine (Ollama) running the quantized qwen3:8b model.

The system bypasses stochastic zero-shot generation by systematically compiling a heavily structured JSON payload that concatenates the topological constraints into a localized context window. The LLM is forced via "Critical Ensemble Architecture" prompt engineering to recognize the absolute hierarchical authority of the *MVL* for the final survival outcome, relegating the *XGBoost SHAP* parameters strictly to localized molecular drivers. Crucially, the algorithmic guardrails explicitly prohibit the model from hallucinating functional biology to force narrative alignment between contradictory population-level correlations (e.g., an inverse TSM association on an mRNA feature paired with a direct TSM association on a methylation feature of the same locus). Instead, the model is mathematically mandated to interpret these multi-omic dualities strictly as systemic tumor heterogeneity. The resulting autonomous output generates a fully auditable clinical synthesis that resolves the mathematical geometry, concluding with targeted therapeutic vulnerabilities derived exclusively from the injected *SHAP* topology.

#### Supplementary Figure Legends

**Supplementary Figure S1. Phase I Multi-Omic Harmonization and Combinatorial Imputation Architecture.** Schematic representation of the determinist algorithmic workflow utilized to process the baseline Pan-Cancer data into the 372 distinct Lineage-Specific Harmonized Multi-Omic Matrices (LiSHMOM). The directed acyclic graph (DAG) illustrates the systematic, node-wise expansion of data handling steps, where distinct imputation algorithms bifurcate to address varying layers of missingness. Gray Nodes (df001–df005): Quality control protocols executing unrecoverable row exclusion and primary harmonization on continuous clinical features strictly prior to omic intervention. Blue & Green Nodes (df006–df011): First-order targeted imputations addressing discrete endpoints, applying baseline arithmetic (Mean, Median, Random) to clinical survival variables, and topological metrics (Mode, Random, *kNN*) to categorical Copy Number Variation (CNV) matrices. Khaki Nodes (df018–df045): Binomial feature resolution executing Bernoulli distribution mapping and probabilistic modeling to correct categorical somatic mutation voids. Thistle Nodes (df054–df342): The terminal, high-dimensional expansion step addressing continuous multi-omic matrices (e.g., protein expression, miRNA, CpG methylation). This layer deploys an integrated battery of nine distinct continuous algorithms, spanning deterministic algebraic baselines (Mean, Median, Random) and iterative machine learning vectors (*kNN*, *missForest*, *XGBoost*, *LightGBM*, *MICE*, and *softImpute iSVD*). Terminal Library (Light Salmon): The convergence of all permutations into exactly 372 distinct LiSHMOM geometric regimes, guaranteeing strict mathematical reproducibility for downstream Phase II CANARY screening. Arrow colors map directly to their originating imputation parent nodes to define structural lineage paths.

**Supplementary Figure S2. Acyclic Data Provenance and Imputation Lineage.** Targeted Sankey hierarchical visualization tracing the absolute computational provenance of the generated Localized in Silico Human Multi-Omic Matrices (LiSHMOM). The directed acyclic flow maps the exact mathematical lineages linking the terminal dataset identifiers (e.g., df nomenclatures) directly back to their specific cancer cohort boundaries and the distinct localized imputation algorithms applied across the seven multi-omic predictor layers. By visually mapping the branching deployment of conventional algebraic and advanced non-parametric data-handling techniques, this unbroken provenance architecture ensures total methodological transparency, documenting precisely how native structural sparsity was computationally resolved prior to the execution of the Phase II survival geometry audits. A dynamic, interactive, plot is available as Figure S2.HTML via GitHub repository: (https://github.com/BioCancerInformatics/CancerRCDPredictor/).

**Supplementary Figure S3. Pervasive Structural Rejection of Linear Proportional-Hazards Across Pan-Cancer Strata.** A pan-cancer feasibility matrix detailing the algorithmic outcomes of the Elastic Net–regularized Cox regression (CoxNet) 'CANARY' diagnostic. The tile map illustrates the structural admissibility of sparse, linear proportional hazards (PH) geometry evaluated across 33 distinct cancer lineages (X-axis) and four discrete survival endpoints: OS, DSS, DFI, PFI (Y-axis). Color mapping denotes the intrinsic terminal failure mode for each targeted stratum. Salmon tiles (COXNET FAIL DATA) represent early model infeasibility driven strictly by insufficient structural survival events or inadequate outcome resolution. Turquoise tiles (COXNET FAIL MU EXHAUSTED) demarcate severe μ-ladder exhaustion, confirming that despite possessing sufficient baseline survival data, the intrinsic multi-omic geometry was massively incompatible with stable, sparse PH convergence. The overwhelming preponderance of μ-ladder exhaustion across the pan-cancer matrix mathematically justifies the immediate disqualification of traditional linear predictive inference in favor of non-proportional, non-linear Phase III topological analyses. White intersections indicate unavailable or non-evaluated cohort strata.

**Supplementary Figure S4. Sparsity Isolation Protocol Panorama and Genotypic Algorithmic Benchmarking.** This composite visualization evaluates the global structural predictability of the survival models synthesized exclusively from strictly isolated binary variables (Somatic Mutations and CNVs) across the 82 computationally viable cancer-endpoint topologies. (A) Algorithmic Geometry Comparison (Density vs Dispersion). Strict statistical density curves (Raincloud distributions) contrasting the rigorous topological stability of the *MVL* ElasticNet SuperLearner against the extreme, heavily dispersed variance of isolated base-learners (*RSF*, *XGBoost*, Survival-*Boruta*, and *MTLR*) when forced to operate on natively sparse, non-continuous clinical architectures. (B) Topological Regularization and Mitigation of Sparsity-Induced Overfitting. A multi-dimensional Dumbbell Trace evaluating the mathematical penalty enforced by the SuperLearner. The blue loci designate the maximum Apparent Peak achieved by the isolated base learners, while the green loci represent the terminal, synthesized predictions constructed by the *MVL* ElasticNet. The bridging segments visually map the absolute shrinkage penalty enforced to stabilize discrete geometric gradients into a biologically plausible cross-validated stability space. (C) Absolute Genotypic Dimensional Payload. Absolute retention frequencies of the strictly binary multi-omic topological features extracted from the native architectures of each independent algorithm. (D) Proportional Retention of the Genotypic Universe. Relative proportional dominance representing the percentage of unique strictly-binary signatures retained by each machine learning topology relative to the absolute pan-cancer universe of available initial predictors (831 Somatic Mutations and 737 CNVs). When strictly isolated from continuous transcriptomic masking, the architectures maintain robust proportional integration of Copy Number Variations.

**Supplementary Figure S5. Sparsity Isolation Protocol Panorama and Algorithmic Concordance Benchmarking.** This multi-dimensional visualization evaluates the strict geometric predictability of the survival models synthesized exclusively from isolated binary variables (Somatic Mutations and CNVs) across the 82 remaining computationally viable cancer-endpoint topologies. Mirroring the primary Phase III architecture, the 2x2 Radial Atlas charts the isolated algorithmic concordance (C-Index) footprints generated by the four Base-Learners when forcibly shielded from continuous multi-omic data: *Random Survival Forests* (blue), Extreme Gradient Boosting (red), Survival-*Boruta* (purple), and Multi-Task Logistic Regression (gold). Each quadrant explicitly isolates a distinct clinical tracking objective: (A) Overall Survival (OS), (B) Disease-Specific Survival (DSS), (C) Disease-Free Interval (DFI), and (D) Progression-Free Interval (PFI). The radial axes define isolated performance trajectories mapped against specific TCGA cohorts, allowing immediate visual diagnosis of the extreme systemic collapse experienced by continuous algorithms operating on binary landscapes (visible as deep, artifactual contractions toward the 0.500 baseline). By mapping these discrete failures against the terminal Multi-View Meta-Learner (*MVL* ElasticNet SuperLearner, mapped in green), the visualization empirically demonstrates how the overarching regularized ensemble structurally overrides localized algorithmic hallucination to construct secure, mathematically stabilized predictive boundaries across purely genotypic terrain.

**Supplementary Figure S6. Non-Linear Signature Topologies and Temporal Performance of Sparse Genotypic Exemplars in Pancreatic Adenocarcinoma (PAAD).** This composite visualizes the structural extraction of prognostic covariance derived exclusively from binary markers across the PAAD Disease-Free Interval (DFI) and Progression-Free Interval (PFI) survival domains. (A) *PAAD-DFI Sparse Multi-Omic SHAP Topology.* A native *SHAP* (*SHAP*ley Additive exPlanations) Beeswarm depicting the global hierarchical importance and directional impact of the retained genotypic signatures (CNVs and Somatic Mutations) natively extracted from the non-linear *XGBoost* architecture. (B) *Time-Dependent MVL Risk Horizon (PAAD-DFI).* Evaluation of the synthesized *MVL* ElasticNet SuperLearner survival predictability across continuous temporal milestones. (C) *PAAD-PFI Sparse Multi-Omic SHAP Topology.* Global *SHAP* Beeswarm illustrating the topological prioritization of distinct binary markers (e.g., CNV and Somatic Mutation interplay) in predicting longitudinal disease progression. (D) *Time-Dependent MVL Risk Horizon (PAAD-PFI).* Corresponding continuous predictive resilience of the overarching sparse *MVL* ensemble.

**Supplementary Figure S7. Post-Hoc Interpretability: *LIME* Geometric Fidelity**. Global evaluation of Local Interpretable Model-agnostic Explanations (*LIME*) deployed across 465 unique patient trajectories (extracted via a controlled sampling stratum of 5 representative risk trajectories per surviving model). The global density distribution mapping the proportion of patients (Y-axis) against their localized Explanation Fit (X-axis, $R^2$) demonstrates a catastrophic linear collapse, proving the violent non-linearity of the overarching SuperLearner matrix. The vertical black dashed line denotes the global median Explanation Fit ($R^2$ = 0.0295), while the blue dotted threshold mathematically bounds the 91.2% of evaluated patients whose linear geometry completely shattered below an $R^2$ of 0.10. Due to its rigid linear approximation constraints, *LIME* fails to adequately trace the deeper synergistic complexity natively captured by multi-dimensional Tree*SHAP* interaction metrics, serving here purely as absolute mathematical proof of non-linear structural dependence.

**Supplementary Figure S8. Global Topological Distribution of Survival Risk via *XGBoost SHAP* Beeswarm Mapping for the READ_OS Supreme Exemplar.** This macroscopic plot aggregates millions of individualized decision interactions to expose the global algorithmic logic directing the READ_OS supreme exemplar landscape. The Y-axis ranks the global topological importance of the top survival determinants in descending order across the cohort. Enforcing a strict top-down hierarchy of survival importance across the pan-omic space, the model identifies the apex drivers across three completely distinct physiological layers: an mRNA network (READ-56.6.3.N.2.4.7.1.3.1) commands primary risk evaluation, immediately followed by specific transcript isoform variability (READ-174.5.2.N.3.2.2.4.4.1) and microRNA interaction parameters (READ-706.4.3.P.3.7.5.2.4.1). The X-axis measures the exact algorithmic log-hazard shift applied against the population survival baseline, with geometric displacement to the right (>0) directly escalating mortality and displacement to the left (<0) acting as a protective physiological layer. Every dot geometrically represents a single READ patient evaluated across strict boundary conditions. The color topography serves as a deterministic indicator tracking the standardized magnitude of the specific biological variable physically recorded within the patient interaction model (Orange equating to mathematical maximum abundance relative to the cohort; Blue equating to profound depletion). Crucially, the mathematical mapping dynamically traces complex, continuous clinical divergence. For example, highly escalated physiological distributions along the dominant mRNA node (READ-56.6.3.N.2.4.7.1.3.1), indicated by the intense orange aggregation stretching strictly to the right axis, invariably operate as a lethal, progressive engine escalating absolute log-hazard. Conversely, tracking distinct color topographies down the hierarchy demonstrates mathematically inverse operations where minimum node abundances (blue) forcefully safeguard survivability.

**Supplementary Figure S9. Local Interpretability of Personalized Overall Survival (OS) Risk via *TreeSHAP* Waterfall Visualization for the Supreme Lethal Trajectory (TCGA-AH-6547-01).** This decision plot decompiles the exact predictive logic of the non-linear SuperLearner architecture for a single representative high-risk Rectal Adenocarcinoma patient (TCGA-AH-6547-01), strictly evaluated against the OS clinical metric. Rather than random selection, the automated Terminal Harvester pipeline mathematically isolated this specific trajectory because it represents the absolute extreme of integrative, lethal multi-omic convergence. Operating natively in this specific survival domain, the X-axis dictates relative OS log-hazard ratios. The predictive process begins at the mathematically calculated population baseline (E[f(x)]). The Y-axis documents the patient's exact biological abundance (mRNA, isoform expression, or miRNA abundance) values for the multi-omic signatures most heavily weighted toward OS outcomes. Crucially, the biological origin of each signature is defined by its Secondary Token identifier (e.g., .4 = miRNA; .5 = transcript isoform abundance; .6 = mRNA expression). Furthermore, in strict adherence to continuous non-linear topography, the algorithm dynamically avoids classical, binary "high/low" classification thresholds. As shown, the exact, individualized omic profile of this patient acts as the deterministic mathematical anchor. Crucially, the mathematical architecture proves that lethal trajectories are not solely driven by extreme biological accumulations (positive values) but by complex, multi-directional dysregulation. For example, alongside positive physiological accumulations imposing profound OS hazard shifts, the algorithm specifically identifies the absolute structural depletion of the transcript isoform signature (READ-359.5.3.N.2.62.62.2.4.1 = -0.808). This specific continuous value acts functionally as a deterministic scalar forcing a consecutive, aggressive OS log-hazard penalty (indicated by orange bars). Purple bars indicate omic values pushing OS risk structurally lower. Through strict cumulative arithmetic of these continuous omic values seamlessly integrating, this patient's specific OS hazard trajectory shifts structurally forward. Finally, the plot visually compresses a vast remainder of evaluated variables into a single block representing roughly 150 other lower-level omic dimensions. The *XGBoost* framework accurately assesses these native biological variables but confirms they collectively represent inert topological noise contributing virtually zero survival deviance compared to the massive dominant predictive power of the lethal signatures. This overrides the population baseline to yield a final, heavily elevated personalized OS mortality prediction located at the apex of the waterfall. This massive Delta physically defines the multi-omic lethal burden identified by the pipeline.

**Supplementary Figure S10. Local Interpretability of Protective Overall Survival (OS) Resolution via *TreeSHAP* Waterfall Visualization for the Supreme Protective Trajectory (TCGA-DC-6683-01).** In stark contrast to Supplementary Figure S9, this plot demonstrates the autonomous capability of the SuperLearner to resolve inverse topological protective states natively within a distinct READ patient profile (TCGA-DC-6683-01). Again, the algorithmic extractor specifically identified this patient to physically uphold the absolute maximum integration of multi-omic protective shielding. While mapping against the identical underlying population baseline (E[f(x)]), this patient demonstrates that the SuperLearner concurrently utilizes multiple distinct omic layers to scientifically drive hazard downward based on rigid mathematical profiles. For example, specific continuous values assigned to profound physiological modulations across targeted transcript isoforms and absolute depletions in top miRNA drivers massively suppress baseline risk. These biological profiles act functionally as deterministic values forcing deep negative log-hazard pushes (indicated by purple bars). Through the strict multi-omic accumulation of transcriptomic and epigenetic continuous values across multiple integrative dimensions, this specific patient plunges structurally backward into a highly protective survival prognosis. Similarly to Supplementary Figure S9, the terminal block compresses roughly 150 other features, representing the mathematical compression of all remaining evaluated omic dimensions across the landscape which were confirmed to be safely reduced without skewing terminal validation. The massive negative Delta generated here explicitly upholds the dual-directional entropy and deep mathematical integration capabilities of the 0.990 absolute compliance architecture.

**Supplementary Figure S11. Phase III Baseline Trajectory Bifurcation of SuperLearner Clinical Probabilities**. Individualized clinical probabilities for the retained baseline derivation cohorts (N

= 10,306 patient trajectories) were extracted across 1-year, 3-year, and 5-year clinical landmarks. The continuous hazard Z-scores were natively generated by the Phase III SuperLearner ensemble and geometrically anchored via Cox-Breslow baseline hazards, achieving 100% algorithmic penetrance without synthetic data imputation. To preserve mathematical clarity, the visualization is strictly bifurcated by endpoint physics: (A) Waterfall heatmap of survival endpoints (OS and DSS) representing the decreasing probability of survival, S(t). Patients are ordered vertically by their continuous risk Z-score to illustrate the smooth probability gradient across the cohort (gradient maps from 0% [Firebrick] to 100% survival [Dodger Blue]). (B) Waterfall heatmap of event endpoints (DFI and PFI) representing the increasing probability of recurrence or progression, 1 - S(t). The gradient is distinctly mapped to reflect the inverted physics of the endpoint (0% risk [Forest Green] to 100% risk [Firebrick]). (C, D) Temporal spaghetti plots mapping the massive density of individual probability trajectories for the Survival (C) and Event (D) endpoints over the 5-year clinical window. Trajectories are uniquely color-coded by the execution architecture utilized: because 100% of the retained Phase III patient records possessed structurally intact multi-omic signatures and successfully utilized the primary SuperLearner (Path A) without requiring algorithmic fallback, all 10,306 patient trajectories are homogeneously plotted in Dark Cyan. This massive, unfragmented probability density cloud visually confirms the mathematical stability of the baseline architecture and serves as the pristine geometric reference required to anchor and rescue fragmented patient profiles during downstream clinical deployment.

#### Comprehensive Legends for Supplementary Matrices and Diagnostic Tables

**Supplementary Table S1. TCGA Lineage Abbreviations and Tumor Classifications.** This table serves as the definitive structural dictionary mapping standard diagnostic acronyms defined by The Cancer Genome Atlas (TCGA) to their complete anatomical and histopathological classifications. To streamline complex multi-omic and geometric evaluations throughout the study, these 33 standardized abbreviations are utilized constantly across all Phase I–III matrices, performance benchmarks, and post-hoc topological interpretability models (e.g., `TreeSHAP` and `LIME`).

**Supplementary Table S2. Data Imputation Strategies and Phase III Harmonization Arrays.** This table summarizes the systematic generation of the 372 candidate preprocessing regimes (arrays df006 through df377) derived from the strict, un-imputed baseline reference topology (df005). To exhaustively evaluate the influence of topological preprocessing on downstream machine learning fidelity, sequential and combinatorial imputation regimens were systematically executed across disparate multi-omic layers (e.g., survival endpoints, CNV distributions, somatic mutation rates). Each distinct algorithmic permutation—utilizing kNN, arithmetic means, positional medians, and randomized distributions—spawned a discrete, harmonized LiSHMOM array. This combinatorial network serves as the comprehensive Phase I baseline pool subjected to subsequent Target-Adversarial Resolution (TAR) viability screening prior to Phase II analysis.

**Supplementary Table S3. Canonical Nomenclature and Phase I–III Terminology Protocol.** This table formalizes the computational architecture and attrition checkpoints of the study. Specifically, it distinguishes between Tier 1 Structural Viability (datasets excluded locally prior to Phase II due to absolute omic sparsity or inherent violations of the CANARY N≥50 policy, such as DLBC and CHOL) and Tier 2 Attrition (datasets computationally rejected during CANARY execution due to target scarcity). Furthermore, it formally defines the Mu-Exhausted state—a mathematically critical distinction delineating cohorts with sufficient survival events that failed linear constraints (CoxNet) but were sequentially admitted to resolve non-linear geometries acting as Viable Survival Pathways within the ultimate Quadripartite ML Ensemble.

**Supplementary Table S4. Global Cross-Cohort Omics Missingness and Predictor Sparsity Diagnostics.** This diagnostic table quantifies the exact structural sparsity across the 33 evaluated cancer cohorts, stratifying patient samples by the quantitative severity of missing data within their explicitly assigned, lineage-specific predictive variables. It acts as the mathematical justification for the 35% geometric exclusion boundary implemented during the Phase III SuperLearner execution. Conclusively, the matrix proves that "partial" missing data drop-offs are an artificial phenomenon; over 98% of the mathematical exclusions mapped globally across the survival topologies (712 out of 727 dropped patients) presented with exactly 100% missing omic predictors, constituting clinical "ghosts." By actively auditing and excluding these hollow vectors, the Phase III architecture guarantees that downstream gradient boosting split-finding pathways are constructed upon genuine biological mass.

**Supplementary Table S5. Pristine Lineage Distribution Ledger: Accounting of sequestered validation patients and multi-omic predictive variable dimensions per cancer cohort.** This table details the exact composition and dimensional scale of the internal blind validation matrix. By cross-referencing the root multi-omic database against the Phase III model derivation dataset, exactly 1,050 unique patient records spanning 27 distinct cancer lineages were identified as strictly absent from the derivation process and safely sequestered. The ledger formally quantifies the distribution of these pristine patient samples and their corresponding multi-omic predictor variables per cohort, documenting the total biological exclusion of the severely fragmented UVM and DLBC cohorts from the validation environment. This dataset guarantees a mathematically pristine test environment, physically withholding the sequestered records from all algorithmic hyperparameter optimization and Phase III elastic net synthesis.

**Supplementary Table S6. Algorithmic NA Panorama: Landscape of topological missingness and autonomous SuperLearner safe-masking frequencies during internal blind validation.** This diagnostic matrix maps the precise frequency of intentional safety abortions executed by the Multi-View Meta-Learning (`MVL`) SuperLearner during clinical deployment on the validation cohort. When deployed against severely fragmented cohorts (e.g., OV, STAD, and SKCM), the native `Boruta` topological imputation layer within the ensemble mathematically failed to converge. Because *Boruta* was a rigidly required constituent of the overarching Phase III ensemble architecture, the SuperLearner dynamically aborted the risk synthesis for these records, generating an intentional non-prediction (NA). This table quantifies these specific masking frequencies across all 27 cohorts, proving that the algorithm's autonomous gating mechanism successfully prevented the generation of hallucinated, uncalibrated risk scores in highly fragmented patient subgroups.

**Supplementary Table S7. Algorithmic Penetrance: Validation lineage verification and quantification of the 100% Dual-Track predictive penetrance (SuperLearner vs. *XGBoost* fallback).** This ledger provides mathematical confirmation of the unbroken computational lineage maintained across the Dual-Track execution architecture during blind validation. It tracks the pipeline's autonomous routing capability: while structurally intact patient records successfully synthesized continuous risk Z-scores via the full SuperLearner (Path A), all safe-masked records documented in the NA Panorama were seamlessly routed into the native *XGBoost* fallback layer (Path B). The table confirms that the *XGBoost* layer—which natively handles missing variables without synthetic imputation—stepped in to evaluate the structurally fragmented subgroups, independently calculating continuous hazard vectors. Conclusively, this matrix proves that the Dual-Track architecture achieved an absolute 100% predictive penetrance (0 NAs), generating valid clinical risk scores for every single sequestered patient across four distinct survival endpoints (OS, DSS, PFI, and DFI) without any unexplained patient attrition.

**Supplementary Table S8. Phase II CANARY Benchmarks and Tier 2 Survival Eligibility Assessments.** This table details the Phase II computational triage of 120 baseline survival combinations across TCGA pan-cancer cohorts. All execution units were subjected to unconstrained, fully-dimensional Cox proportional hazards (CoxNet) profiling. Cohorts marked with Target Scarcity failed to clear strict statistical minima (N≥50 or E≥20) and were algorithmically decommissioned (Failed). Conversely, cohorts that satisfied survival availability prerequisites but mathematically resisted linear optimization due to high-dimensional complexities or strict proportional hazards violations were identified as Mu-Exhausted. These combinations were flagged as structurally valid and definitively Admitted to Phase III, mandating the deployment of advanced, non-linear machine learning architectures (*XGBoost*, *RSF*, Survival-*Boruta*, and *MTLR*) to resolve their geometric topologies.

**Supplementary Table S9. Baseline Macro-Level Sample Diagnostics and Omic Density Mapping (Tier 1).** This table outlines the absolute macro-level distribution of patient samples per cancer lineage alongside the corresponding density of predictive signatures, stratified by specific omic layers within the baseline unimputed df005 reference matrix. To ensure structural transparency, omic predictor variables are explicitly compartmentalized by their discrete numerical tokens into seven biological layers: protein abundance (.1), somatic mutation status (.2), copy number variation (.3), microRNA expression (.4), transcript isoform abundance (.5), mRNA expression (.6), and CpG methylation (.7). Crucially, this diagnostic matrix provides the empirical basis for Tier-1 cohort attrition, explicitly detailing the severe quantitative limitations—such as total sample paucity (e.g., CHOL, N=45) and pervasive structural omic opaqueness (e.g., DLBC, 0 omic variables; UCS, 1 omic variable)—that trigger immediate protocol ejection prior to Phase II analysis.

**Supplementary Table S10. Quadripartite ML Ensemble Concordance and Temporally Truncated AUC Configurations.** This comprehensive metric log tracks the predictive parity and divergent topological evaluations between the nested base-learners (`XGBoost`, `RSF`, `MTLR`, `Boruta`) versus the overriding `MVL` ElasticNet SuperLearner, mapping spatial stabilization specifically across 1-, 3-, and 5-year clinically truncated survival horizons.

**Supplementary Table S11. Quadripartite Concordance and Algorithm-Specific Importance Metrics (Node Gain & Gradient Norm).** This granular evaluation decompiles the predictive superiority of the baseline prognostic features. It establishes individual algorithmic weight assignments, contrasting explicit `XGBoost` Node Gain against `RSF` spatial proximity and `MTLR` gradient norm metrics to assert absolute non-linear feature dominance.

**Supplementary Table S12. Master Matrix: Topologically Validated Golden Anchor Features and RCD Signatures.** Housing the fully deconvoluted Phase III multi-omic payloads, this global hierarchical database traces thousands of highly dominant predictor variables back to their exact transcriptomic, somatic, or genotypic origins. It enumerates precise biological attributions, validating the specific pathways dictating the prognostic architecture globally.

**Supplementary Table S13. Genotypic Superiority and Sparsity Rescue Effect Outcomes.** This topological breakdown identifies cohorts uniquely governed by the *Sparsity Rescue Effect*. It mathematically documents where localized transcriptomic cascades collapsed, forcing the SuperLearner to rely on specific underlying somatic and CNV mutations to rescue the clinical survival bounds against the dominant RNA-centric baseline.

**Supplementary Table S14. Sparsity Isolation Reject Ledger.** Functioning as a strict structural quarantine report, this table details the absolute mathematical suppression of strictly binary genomic signatures (Somatic Mutations and Copy Number Variations) evaluated during the Sparsity Isolation Protocol. It documents the exact global missingness and sparsity ceilings for the specific discrete markers that completely failed to provide independent topological variance in the absolute absence of continuous transcriptomic masking.

**Supplementary Table S15. Quadripartite Sparsity Concordance and Algorithm-Specific Importance Metrics for Somatic Mutation and Copy Number Signatures (Node Gain & Gradient Norm).** This granular evaluation decompiles the absolute retention of the 1,253 somatic mutation and copy number signatures in the absolute absence of continuous omic features. It establishes individual algorithmic weight assignments, contrasting `Boruta` mean importance and explicit `XGBoost` Node Gain against `RSF` spatial proximity and `MTLR` gradient norm metrics to assert absolute non-linear feature dominance.

**Supplementary Table S16. Phase III Pan-Omic Ejected Signatures and Node-Splitting Geometric Exclusions.** Functioning as a diagnostic quarantine report for the global pan-omic ensemble, this table lists the explicitly isolated omic variables (spanning all seven molecular layers) that catastrophically failed the minimum topological split boundaries during unrestricted model competition. It documents the exact missingness and sparsity rates that triggered algorithmic ejection to protect the finalized computational geometry.

**Supplementary Table S17. *SHAP* Interaction Exemplars: Extreme Multi-Omic Drivers and Convergence Trajectories4** This final matrix traces the extracted patient-level precision trajectories. By mathematically anchoring localized *TreeSHAP* derivations to the SuperLearner, it captures the exact non-linear, multi-omic synergisms dictating extreme patient mortality boundaries beyond the standard population topology.

**Supplementary Table S18. Phase III Global Trans-Signature Dependencies and Mathematical Interaction Archetypes.** This table catalogs the 20,604 statistically significant trans-signature dependencies extracted from the Phase III precision oncology interaction analysis. From an initial universe of 37,890 audited multi-omic interactions across 93 isolated survival cohorts, dependencies were filtered using a rigorous joint statistical and effect-size threshold (Benjamini-Hochberg False Discovery Rate [FDR] $< 0.01$ and absolute Spearman correlation [$|\rho|$] $\geq 0.30$ within a specific molecular gradient). The table details the directed biological asymmetry of each dependency, specifying the Primary Signature (the principal biological driver of lethality or protection) and the color var Partner (the modulating signature that structurally interacts with the driver). The specific biological layers of these features are formally identified by their Primary Omic Token and Partner Omic Token. Crucially, the table provides the mathematical proof defining the non-linear geometry of each interaction:
Interaction Strength: A geometric proxy representing the spatial volume and magnitude of the cross-talk topology.
Spearman HighZone/MidZone CrossTalk: The directed correlation metric that defines the dependency archetype. A positive correlation defines Synergistic hazard amplification (where the partner exerts a lethal push effect), whereas a negative correlation mathematically defines Antagonistic functional rescue (where the partner exerts a protective pull effect).
Mathematical Classification: The formal integration of these metrics into distinct dependency archetypes (e.g., Synergy, Antagonism, and Bifurcation).

**Supplementary Table S19. Primary Signature Embodiment Profiles and Mathematical Interaction Context.** This table provides the biological translation (Concept Embodiment) for the 20 supreme multi-omic exemplars highlighted in the manuscript (Section 3.14.1). Each primary signature's algorithmic nomenclature is systematically decoded into its fundamental genetic elements, explicitly isolating the targeted omic layer and linking it directly to the functional Regulated Cell Death (RCD) mechanics driving patient survival. To ensure absolute fidelity to the non-linear interaction topologies proven in the manuscript (e.g., Figure 8), the table rigorously preserves the exact geometric context of each dependency. Every primary exemplar is strictly mapped to the specific clinical cohort and survival metric in which its dependency was audited. Furthermore, the corresponding modulating partner signature is fully decoded (including its constituent genetic elements and RCD pathways) and aligned with the definitive mathematical classification (Synergy, Antagonism, or Context-Dependent Bifurcation) that dictates the hazard amplification or functional rescue. This structural mapping provides a comprehensive, audit-ready index translating abstract AI predictions into actionable precision oncology biology.

**Supplementary Table S20. Phase III Sample Retention & Algorithmic Penetrance: Formal audited accounting of baseline cohort gating constraints, statistical exclusions, and quantification of the 100% 5-model algorithmic penetrance across the 96 Phase III derivation cohorts.** This matrix details the absolute geometric boundaries and retention rates of the primary Phase III reference architecture. The audit reveals a global average sample retention rate of 87.17% across the baseline derivation matrices. The ledger mathematically attributes the 12.83% baseline sample attrition exclusively to rigid clinical endpoint exclusions and the strict structural enforcement of the <35% omics-missingness threshold required to sustain deep ensemble synthesis. Furthermore, it logs the complete global exclusion of cohorts lacking genome-wide significance or sufficient event targets (e.g., DLBC, CHOL, and KICH). Crucially, the table verifies that for the 87.17% of patient samples that successfully survived this geometric gating, the pipeline achieved a flawless 100% algorithmic penetrance. Every retained patient within the baseline derivation cohorts successfully yielded a complete, fully bifurcated survival probability trajectory across all four independent ML-based algorithms and the composite SuperLearner. This mapped a pristine macroscopic visualization space of 10,306 continuous clinical trajectories without generating a single mathematical outlier or topological gap.

**Supplementary Table S21. Baseline Convergence Audit of the Clinical Blind Validation Cohort:** Documentation of topological singularities and convergence boundaries during internal blind validation. This diagnostic ledger quantifies the localized mathematical failures—specifically, infinite partial likelihood coefficients—encountered when anchoring highly fragmented validation signatures (e.g., specific LGG cohorts) against the continuous Phase III Breslow baseline survival estimator (basehaz()). The matrix details the exact frequency, clinical endpoints, and cohort distribution of these topological crashes that necessitated explicit UI masking during the final clinical probability deployment. By auditing these events, this table provides complete mathematical transparency regarding the structural boundaries of the inference engine when operating under conditions of extreme multi-omic sparsity.
